# Investigation of RNA metabolism through large-scale genetic interaction profiling in yeast

**DOI:** 10.1101/2020.10.04.325191

**Authors:** Laurence Decourty, Christophe Malabat, Emmanuel Frachon, Alain Jacquier, Cosmin Saveanu

## Abstract

Gene deletion and gene expression alteration can lead to growth defects that are amplified or reduced when a second mutation is present in the same cells. We performed 154 genetic interaction mapping (GIM) screens with mutants related with RNA metabolism and measured growth rates of about 700 000 *Saccharomyces cerevisiae* double mutant strains. The screens used the gene deletion collection in addition to a set of 900 strains in which essential genes were affected by mRNA destabilization (DAmP). To analyze the results we developed RECAP, a strategy that validates genetic interaction profiles by comparison with gene co-citation frequency, and identified links between 1 471 genes and 117 biological processes. To validate specific results, we tested and confirmed a link between an inositol polyphosphate hydrolase complex and mRNA translation initiation. Altogether, the results and the newly developed analysis strategy should represent a useful resource for discovery of gene function in yeast.

## Introduction

The process of assigning function to a gene involves switching it off, partially or totally, and evaluating a phenotype. A major limitation of this approach is that genes do not function in isolation and evolved from other genes, sometimes following cataclysmic events, such as whole genome duplication, or more restricted chromosome segment duplication (reviewed by Dujon, 2010; Marsit *et al*, 2017). As a consequence, removal or alteration of a gene from a duplicated pair might show no effect under standard culture conditions. The presence of duplicated genes can increase fitness, a phenomenon that was confirmed by testing single gene deletion mutants in yeast (Gu *et al*, 2003), by performing experimental evolution under random mutagenesis conditions (Keane *et al*, 2014) or by comparing the effect of duplicated gene pair deletion in comparison with singletons in *S. cerevisiae* versus *S. pombe* (Qian & Zhang, 2014). Gene duplication is just the simplest illustration of how cells can adapt to mutations. In many other cases, the flexibility and robustness of cellular pathways allows adaptation of cells to gene loss.

A way to identify and study gene redundancy and robustness against mutations is to combine perturbations for several genes in a single strain and look at the resulting phenotype. This strategy worked well for studies such as the identification of genes involved in the secretory pathway (Kaiser & Schekman, 1990). It only became a systematic way to study gene function when methods to identify and quantify growth of combinations for thousands of mutants became available (Tong *et al*, 2004; Decourty *et al*, 2008; Pan *et al*, 2004; Schuldiner *et al*, 2005; reviewed in Dixon *et al*, 2009). Simultaneous perturbation of two genes can result in various effects on growth. Sometimes, the combination is neutral, sometimes it leads to a strong growth inhibition (synthetic lethality) and sometimes one mutation can hide or overcome the effects of the other (reviewed in Costanzo *et al*, 2019). Altogether, these effects are covered by the convenient umbrella term of ‘genetic interactions’ (GI).

The behavior of a gene variant over many screens establish a GI profile (Schuldiner *et al*, 2005; Decourty *et al*, 2008; Costanzo *et al*, 2010). A similarity of GI profiles can predict physical interactions of the corresponding proteins in complexes and subcomplexes. For example, the analysis of proteasome component mutants, allowed to correctly assign proteins to the corresponding proteasome sub-complexes (Breslow *et al*, 2008). Large scale double mutant screens can also associate previously uncharacterized genes with specific pathways. For example, the RNA exosome co-factor Mpp6 was identified on the basis of the observed synthetic lethality between its gene deletion and the absence of the nuclear exosome component Rrp6 (Milligan *et al*, 2008). Thus, description of GIs serves several goals. It can identify the potential function of genes and find combinations of mutants that uncover phenotypes otherwise hidden by gene redundancy. It can also help in understanding the evolutionary trajectory of duplicated genes towards redundancy or towards unrelated cellular processes (for example, Kuzmin *et al*, 2020). These goals require high quality and validated large scale results, based on independent studies performed under a variety of culture conditions.

Early systematic gene deletion combination screens were restricted to the study of non-essential genes. To investigate essential gene mutants, several strategies have been used, including mRNA destabilization, the study of mutations leading to thermosensitivity, CRISPR genome editing and transposon insertion analysis. The “decreased abundance by mRNA perturbation”, DAmP, strategy was the first to be used for systematic investigation of hypomorphic alleles of essential genes in yeast and is based on the addition of a long extension downstream the stop codon position of targeted genes. This extension leads to mRNA destabilization through nonsense-mediated mRNA decay (NMD, Schuldiner *et al*, 2005). Three independent systematic yeast libraries were built using variations of this strategy, for large scale genetic or chemogenomic screens. One did not include molecular barcodes in the strains (Breslow *et al*, 2008) and can not be directly used for growth estimation in pooled mutant assays. For such assays, a second collection was generated in which “molecular barcodes”, unique artificial short sequences flanked by universal sequences allowing their amplification, were included at a specific genomic locus for each strain (Yan *et al*, 2008). However, since the modified locus and the barcode are not physically linked, this second collection was not usable for genetic interaction mapping (GIM) screens, which depend on co-segregation of mutant and barcodes in a pooled population of mutants (Decourty *et al*, 2008). To solve this problem, we generated a third DAmP collection, where barcodes are present at the modified locus. These strains can be used both for measuring cell numbers in GIM screens and for transcript quantitation, by reverse transcription and barcode amplification (Decourty *et al*, 2014).

In addition to DAmP essential gene perturbations, recent methods that are able to generate collections of mutants analyzed by DNA sequencing became available. For example, CRISPR interference was used to generate new collections of yeast mutants (Smith *et al*, 2017) and was adapted to the study of genetic interactions under several growth conditions (Jaffe *et al*, 2019). Alternatively saturated transposon insertion coupled with sequencing allows the exploration of a broad spectrum of mutations, including protein truncation or transcription deregulation, and can be used to characterize the function of essential genes (Michel *et al*, 2017). These new methods remain technically challenging and have not yet been used on a large scale. Thus, results about GIs from systematic large-scale studies using essential gene variants in yeast are, for the moment, restricted to thermosensitive (TS) and DAmP alleles under the specific conditions of the synthetic genetic array, SGA (Tong *et al*, 2004), screens (Costanzo *et al*, 2016).

The modest overlap between SGA results and those obtained for the same pairs of mutant genes by CRISPRiSeq (Jaffe *et al*, 2019) confirmed previous demonstrations that culture conditions might be crucial for the detection and measurement of GIs (Martin *et al*, 2015; St Onge *et al*, 2007). In this respect, the GIM screens (Decourty *et al*, 2008) performed under selection with antibiotics that affect mRNA translation, are particularly good at detecting GIs for factors involved in RNA metabolism. For example, the strong effect observed in GIM screens for double deletions involving components of the ribosome quality control complex and the SKI complex (Brown *et al*, 2000), was validated on individual strains only in the presence of low concentrations of hygromycin B, a translation inhibitor (Defenouillère *et al*, 2013).

The specific conditions of GIM screens that had the potential to identify new GIs, and the availability of the barcoded DAmP collection, compatible with these screens (Decourty *et al*, 2008, 2014), motivated us to generate a new set of large-scale GIs in yeast. We selected 154 genes, mostly related with RNA metabolism, and tested their GIs when combined with the 5500 gene deletions and 900 DAmP alleles for essential genes.

The size of the generated data set and the fact that RNA metabolism perturbation directly or indirectly affects most cellular processes ensured that our results cover a large variety of functions. A major challenge was extracting meaningful information from the obtained results. SAFE, a recently developed method that is specific to GI networks (Baryshnikova, 2016), uses the local neighborhood in complex networks to identify enrichment for specific annotations. We present here a different approach, called RECAP for “**R**ational **E**xtension of **C**orrelated **A**nnotations and GI **P**rofiles” that starts instead from links between genes inferred from co-occurrence in publications, based on the set of scientific articles curated by the Saccharomyces Genome Database (Cherry *et al*, 2012). This approach uses well annotated gene groups in combination with GI profile similarity to find which mutants behave “as expected” from previous studies. Only the validated mutants were then used to extend the network of related genes and predict the potential association of hundreds of genes with specific cellular processes or multiprotein complexes.

## Results

### Correcting for pleiotropic behavior improves the specificity of GIs

The choice of the 154 mutants used to query the collections of gene deletion and DAmP strains was guided by published GI results and gene annotations, with a focus on RNA-related processes, as summarized in **Fig. 1 A** and listed in **Supplementary Table 1**. In addition to factors affecting RNA transcription, export and degradation, a set of 25 mutants included genes that were either directly or indirectly related with proteasome function, or had shown genetic interactions with proteasome deficiency. To limit the bias induced by the choice of tested mutants, we included 14 metabolism genes, 21 genes affecting other processes and 10 unannotated or poorly annotated genes. Gene deletion strains from the collection of haploid strains (Giaever *et al*, 2002) were modified to be suitable for GIM screens by changing the geneticin resistance cassette with the **MATα** haploid specific nurseothricin resistance cassette (Malabat & Saveanu, 2016; Decourty *et al*, 2008). The set of tested mutants (**Fig. 1 B**) included also 11 strains modified by the DAmP strategy and 15 deletion mutants affecting individual snoRNA genes, involved in targeting 2’O-methylation and pseudouridylation of rRNA (Kiss, 2002). For 4 essential genes, we decided to test the flexibility of the GIM approach, and evaluate if a *Tet-off* system (Wishart *et al*, 2006) could be used to study GIs of essential genes, as an alternative to DAmP or TS mutants. In these cases, the screen protocol, schematically depicted in **Figure 1 – figure supplement 1**, included the addition of doxycyclin in the final culture where double mutant haploid strains are selected in the presence of geneticin and nourseothricin.

**Figure 1.**
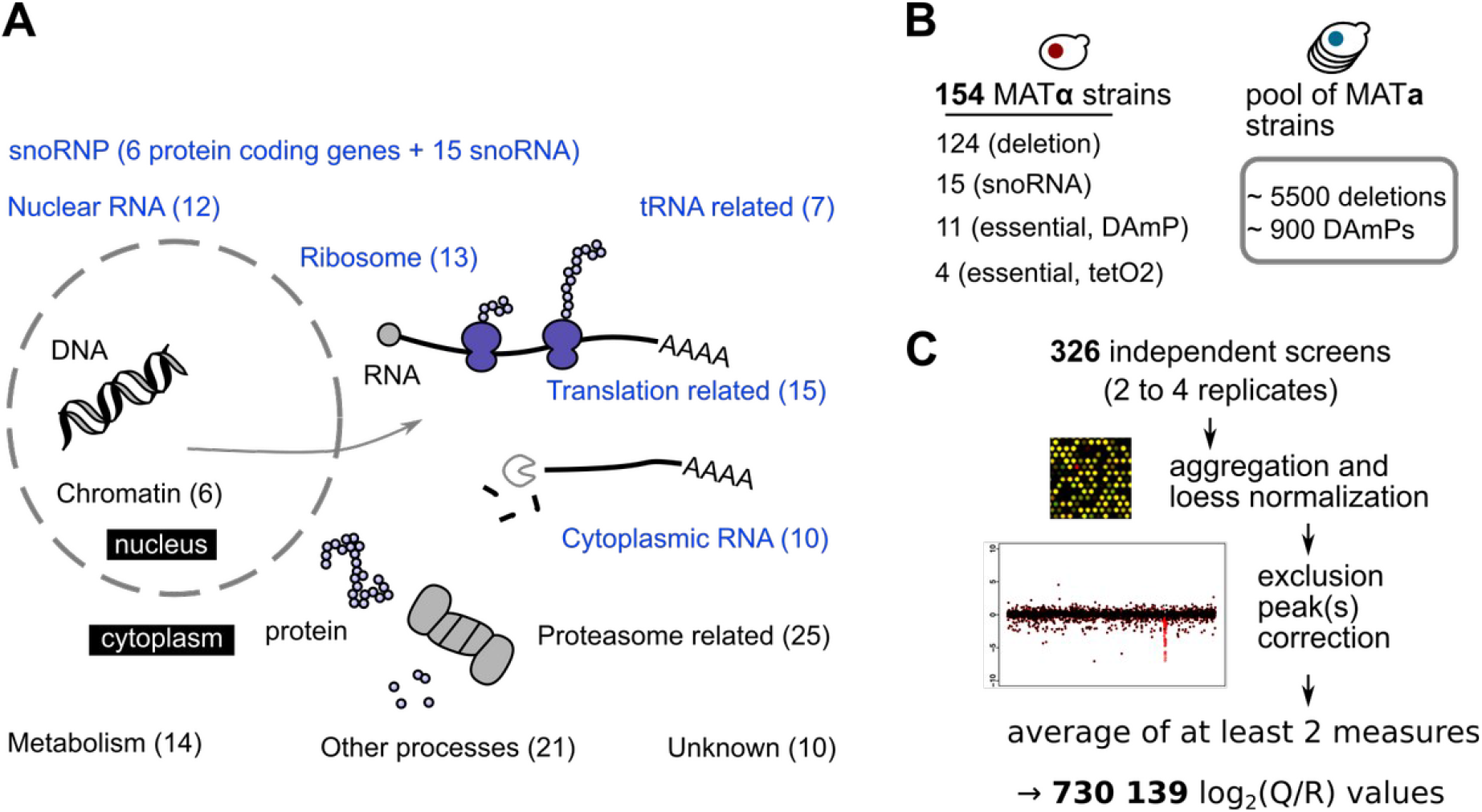
Overview of the cellular functions of query genes tested in GIM screens. **A**. Classification of the tested mutants in broad groups associated with major cellular processes, including mRNA translation, protein degradation and ribosome function and biogenesis. The number of genes for which we performed GIM screens from each class is indicated, with RNA-related processes highlighted in blue. **B**. Three types of mutants were used in screens, mostly gene deletion, but also DAmP and regulated expression strains (left). The pool of barcoded deletion strains used in each screen was supplemented with our collection of DAmP strains for essential genes (right). **C**. The workflow for analyzing the microarray results involved normalization, correction of the signal peaks that indicate the low frequency of meiotic recombination that occurs for loci situated close on the same chromosome and averaging of values from independent screens. The initial signal and corrected version for each of the screens are presented in **Supplementary Data Set 1**.

Each screen was performed at least twice, leading to results for 326 independent experiments (list of the experiments in **Supplementary Table 2**). DNA extracts from pools of double mutants were labeled and used for microarray hybridization. The obtained microarray data were normalized and the results for the two barcodes that are characteristic for each mutant were aggregated. Finally, the specific peak that corresponds to the decreased meiotic recombination frequency for genes located close to the “query” gene locus (Decourty *et al*, 2008; Baryshnikova *et al*, 2013) was corrected (**Fig. 1C and Supplementary Data Set 1**). The raw results of query versus reference ratios (Q/R) were normalized across genes and screens, to obtain a primary table of 730 139 ratios between the levels of a mutant in a given screen and its levels in a control population (log_2_ transformed values, **Supplementary Table 3**). Negative values of log_2_(Q/R) correspond to a depletion of a given mutant when combined with the specific “query” allele, null values indicate no interaction, while positive values suggest an epistatic relationship.

When looking at the distribution of the obtained values for each mutant, we observed that several strains showed large relative growth defects in screens performed with unrelated mutants. For example, the distribution of the scores observed for VPS63 was different from the average cumulative distribution of the measured growth defects, with a much larger spread (**Fig. 2 A**). For comparison, the distribution of scores for the nuclear exosome factor MPP6, known to show a very specific response to perturbations of the nuclear exosome (Milligan *et al*, 2008), showed a very steep slope (**Fig. 2 A**). To identify other mutants following the same trend like VPS63, we took the number of screens in which the log_2_(Q/R) of a given mutant was inferior to -1.25, and expressed it as the ratio of the total number of screens in which the mutant was measured. The calculated value is a “pleiotropy index” (PI) specific to our data set and has values between 0 and 1, with higher values indicating a broader shoulder of the values distribution. For example, a PI value of 0.5 would indicate that a gene deletion was seen deleterious for growth in combination with half of the query genes used in the 154 screens. The values of PI were 0.44 for VPS63, the maximum in our results, 0.39 for KEX2, 0.26 for VPS3, and 0.01 for MPP6 (**Supplementary Table 3**). Only about 11% of the tested strains had PIs higher than 0.1 (539 out of 5063 measured strains). When ranking genes in decreasing order of measured PI, we observed an enrichment of functions related to intracellular vesicular transport. Thus, 15 of the top 32 genes were annotated with the GO term for biological process “16192”, “vesicle-mediated transport”, with an adjusted p-value for functional enrichment of 5.4×10^−6^, as tested using the g:Profiler tool (Raudvere *et al*, 2019). Perturbation of the intracellular transport of macromolecules or metabolites can affect a relatively large number of different cellular processes, which probably explains this result.

**Figure 2.**
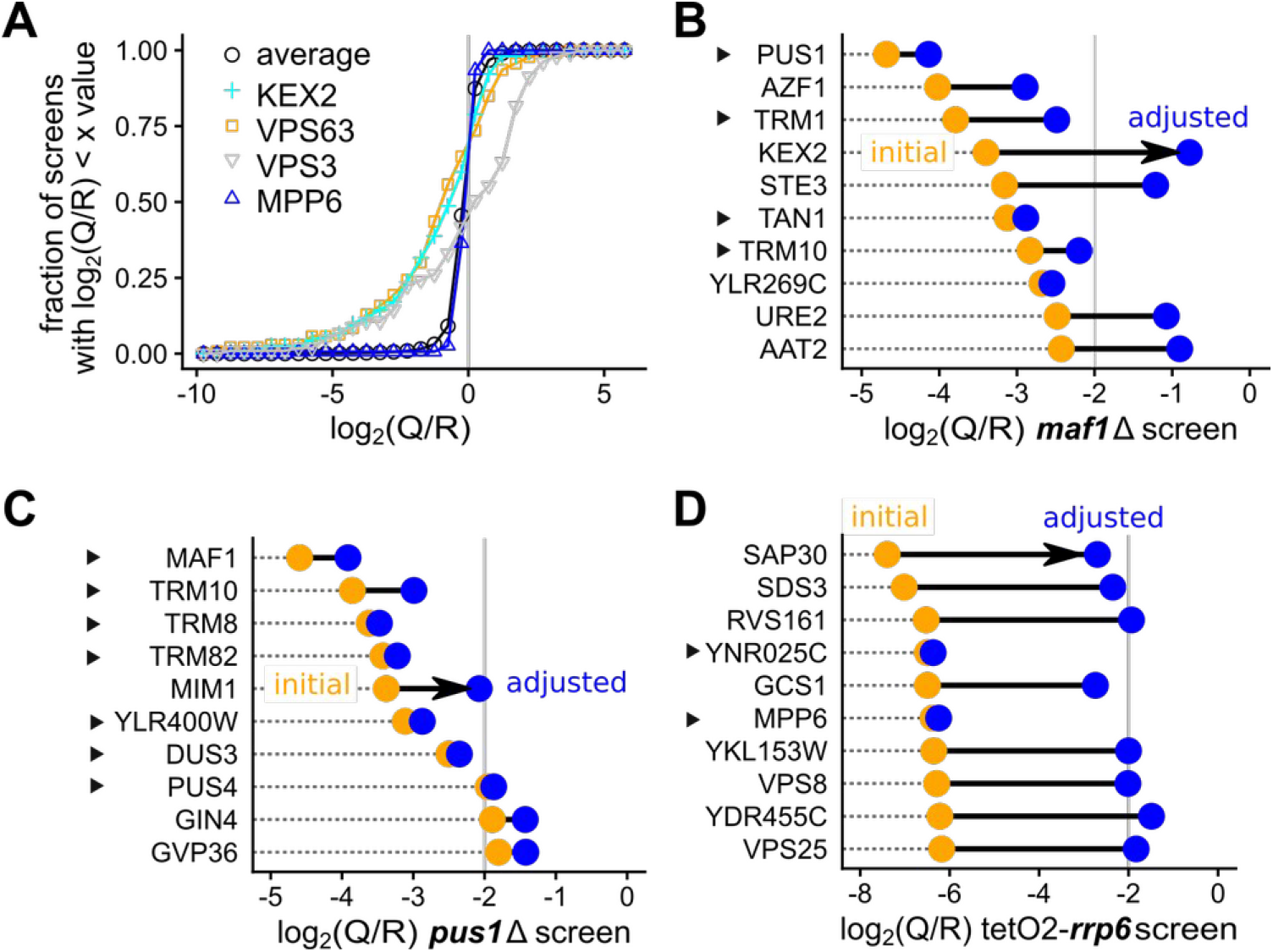
Correcting for pleiotropy improves ranks of genes functionally related with the tested mutant. **A**. For each measured effect of a mutant in the 154 screens, we evaluated the cumulative distribution of the log_2_(Q/R) values. Results for genes having an unusual behavior are displayed, including KEX2 (blue cross), VPS63 (orange square), VPS3 (downside gray triangle) compared with the average for all screens (black circle) or for a mutant showing highly specific interactions, MPP6 (upside dark blue triangle). Examples of applying a correction based on pleiotropy to the ranks of the 10 best hits for the screens performed with *maf1*Δ (**B**), *pus1*Δ (**C**) and tetO2-*rrp6* (**D**). Initial scores are indicated with orange dots and adjusted values are illustrated as blue dots. Genes marked with a triangle correspond to mutants that are known to affect the same pathway (tRNA synthesis for MAF1 and PUS1 and RNA degradation in the nucleus for RRP6).

Since mutants showing pleiotropic effects are not informative and their effects can mask more interesting functional interactions, we adjusted the screen result values by multiplication with a correction factor derived from the PI. Among several possible transformations, we applied one that improved the identification of known GIs in the screens performed with *maf1*Δ, *pus1*Δ and temporary depletion of RRP6 (tet02-*rrp6* strain). After testing several transformations, we chose to multiply the original log2(Q/R) values with (1 - PI)^3^, which had little effect on most results, but diminished the relative contribution of the highly variable mutants. More than 80% of the results were only slightly corrected, by factors between 0.8 and 1, and only 3% of the results were decreased with a factor of more than 2 (**Fig. 2 – figure supplement 1**).

When looking at the top 10 hits of the screens mentioned above, we observed the effects of the applied corrections. For example, Maf1 is a major regulator of RNA polymerase III activity (reviewed in Boguta, 2013) and is thus tightly linked with tRNA metabolism. The first and third most affected gene deletions affected by *maf1*Δ, PUS1 and TRM1, are both linked with tRNA modification (reviewed in Hopper, 2013). However, the fourth and fifth values in the *maf1*Δ screen correspond to KEX2, coding for a protease involved in the secretory pathway, and STE3, a membrane receptor. The pleiotropy correction effectively filtered out these results, while improving the ranks of TAN1 and TRM10, linked with tRNAs (**Fig. 2 B**) which were promoted in the top 5 of the adjusted results. In the screen using the *pus1*Δ strain, affecting tRNA modification, the fifth hit was most affected by the pleiotropy correction (**Fig. 2 C**). MIM1, the corresponding gene, has no known link with tRNA. However, both the first four hits and the next 3 correspond to genes affecting tRNA modification or synthesis: MAF1, TRM10, TRM8, TRM82, DUS3, YLR400W, which overlaps partially DUS3, and PUS4.

The effects of the applied corrections were occasionally impressive, as seen with the screen in which the expression of RRP6 was blocked by doxycyclin addition during the growth of the double mutant strains. RRP6 is a 3’ to 5’ exonuclease that associates with the nuclear exosome and is involved in RNA synthesis, maturation and degradation (reviewed in Fox & Mosley, 2016). However, 8 out of 10 top hits in the corresponding screen did not match known genetic interactions for this factor (**Fig. 2 D**). These 8 factors were among those highly variable in many screens and the corresponding values were strongly reduced by applying the pleiotropy correction. The remaining genes, MPP6 and deletion of overlapping YNR025C, were the top hits of the tetO2-*rrp6* screen, in agreement with previous results obtained with an *rrp6*Δ strain (Milligan *et al*, 2008). Thus, correction for pleiotropic effects can help in recovering important functional information, with variable efficiency depending on each screen particular conditions. Among the corrected results, there were 2356 gene deletions and 402 DAmP mutants with at least one adjusted log _2_(Q/ R) value lower than -1, thus showing a ratio for query screen to reference of at least 2 (**Supplementary Table 4**). Next, we wondered how good the measured GIs were, and, for this task we used several criteria, as described below.

We were confident that the identified GIs were meaningful, since the screen results were compatible with current known annotations and with previously published data sets (annotated examples in **Fig. 2 B-D**). However, we wanted to assess the quality of the measured GIs globally. We thus took advantage of the fact that GI profiles, the set of values obtained for a given mutant, provide more information than direct GIs for inferring gene function (Decourty *et al*, 2008). Thus, we used correlation of GIs to test the validity of the newly obtained data set. For a first validation of the adjusted results we looked for the similarity of GIs for the same gene mutant when tested in the query **MATα** strain (Nat^R^ marker) or in the tested pool of **MATa** strains (Kan^R^ marker). Pearson correlation values of the GI profiles for the 127 pairs of genes tested independently were clearly skewed towards positive values, as expected. In contrast, correlation for all the possible pairs of GIs in the data set showed a bell-shaped distribution centered on zero (**Fig. 3 A**). Thus, the phenotype of mutating the same gene was similar, whether the mutant was present in the query strain or in the pool of tested strains. We performed a similar analysis for cases of overlapping gene deletions to analyze the correlation between the effect of independent mutations affecting the same locus. Since two deletions affect the same gene, the two strains should behave similarly in the screens. For the available pairs of overlapping gene deletions, we observed a strong positive correlation for their GI profiles (**Fig. 3 B**, right). In conclusion, correlated GI profiles for mutants affecting the same gene (**Fig. 3 A**) and for overlapping mutants (**Fig. 3 B**) globally validated the quality of our large-scale screen results.

**Figure 3.**
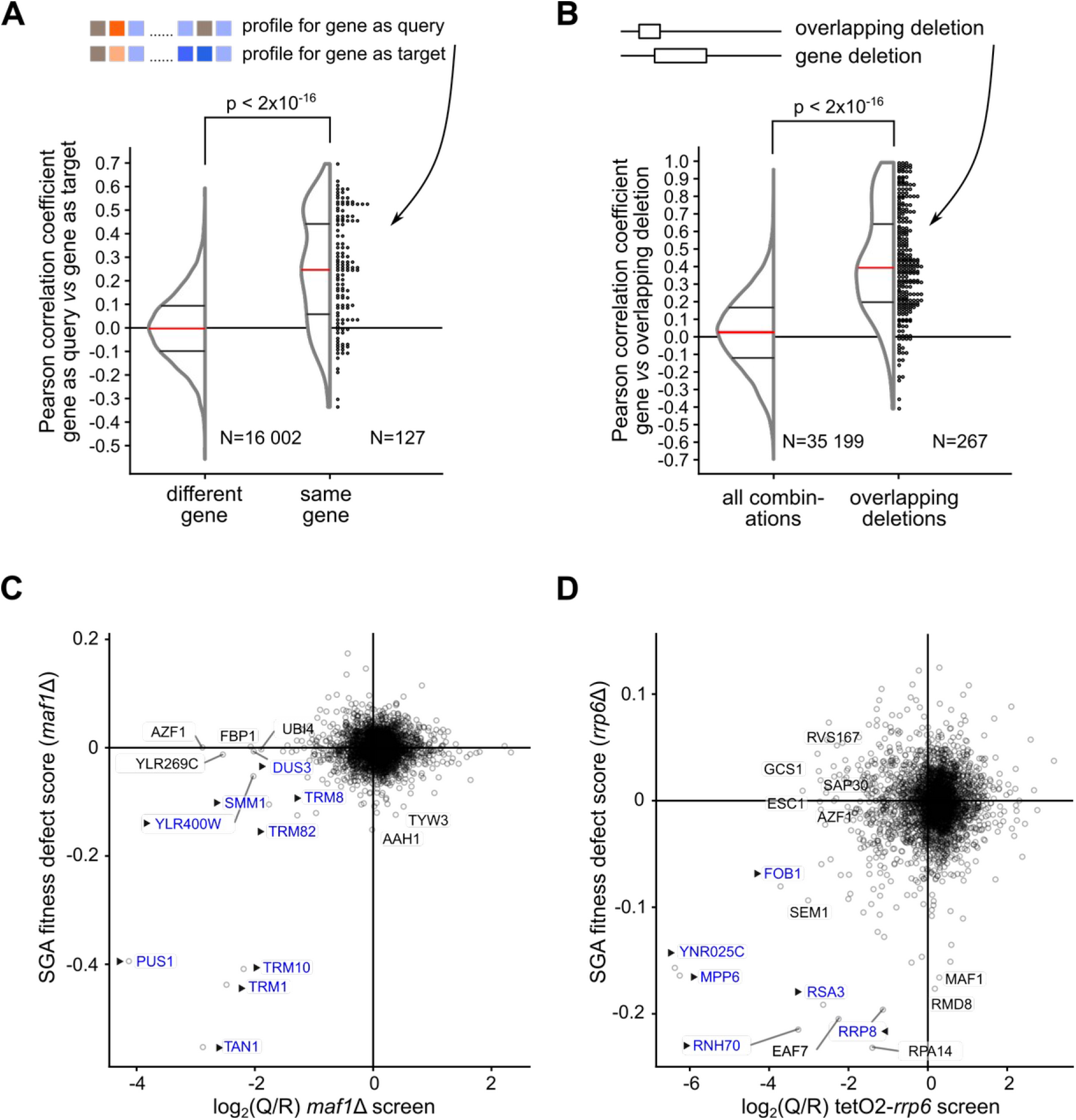
Large-scale validation of GIM data based on GI profile similarity analysis. **A**. Comparison between the GI profiles of the same gene mutant were performed on 127 query genes (out of 154 screens) that were also measured as “target” mutants. The distribution of the measured Pearson correlation coefficients are shown either for this situation, at the right of the plot, labeled “same gene”, and for all the possible other 16 002 distinct pairs of the 127 mutants, as background, labeled “different gene”, at the left. The similarity of the two distributions was evaluated using the non-parametric Wilcoxon rank sum test (p < 2*10^−16^ for the null hypothesis, no difference). Dots at the right of the distribution representation correspond to individual Pearson correlation values. **B**. We identified 267 situations where the deleted region for a gene or pseudogene had an overlap with the deleted region of another gene and extracted the Pearson correlation values for the corresponding GI profiles. The distribution of Pearson correlation coefficient values for all possible pairs involving genes for which overlapping deletions were tested (“all combinations”, left) and for overlapping deletion pairs (“overlapping deletions”, right) is shown. The two populations of values were different, as estimated with the non-parametric Wilcoxon rank sum test (p < 2*10^−16^). **C**. Example of similarity for GIM and SGA results. Scatter plot showing the top 10 genes most affected in either SGA or GIM screens performed with *maf1*Δ (GIM) compared with the same mutant in the SGA data (Costanzo *et al*, 2010). In both C and D plots, triangles and blue color indicate genes that are known to be functionally linked with the screen query gene. **D**. Example of results obtained using transcription repression for the query gene RRP6. Scatter plot to compare the results of the GIM tetO2-*rrp6* screen and SGA *rrp6*Δ screen. YNR025C partially overlaps the exosome-associated factor gene MPP6.

The GI profiles from the GIM screens were compared with results obtained with similar mutants using the SGA approach (Costanzo *et al*, 2010). There were 52 screens that were done with mutants affecting the same gene in the two sets. For many screens, a positive correlation coefficient between the SGA and GIM results indicated that part of the observed GIs were similar among these two independent assays (**Fig. 3 – figure supplement 1**). On other occasions, no correlation could be detected. This discrepancy can be explained by the fact that the GIM and SGA screens were performed in completely different culture conditions. Alternatively, for query gene mutants that have little impact on gene function, with no strong GI detected, a lack of correlation between results is to be expected.

We focused on the situations where SGA and GIM results were correlated, as these cases depend on GIs that were robustly detected across assays and laboratories. As examples of GIs responsible for the observed correlations, we present the comparison of the GIM and SGA screens performed with *maf1*Δ (**Fig 3 C**) and the comparison of the SGA *rrp6*Δ screen with RRP6 depletion GIM screen using a *Tet-off* system (**Fig. 3 D**). The correlation between RRP6 deletion and its depletion by *Tet-off* shows that the GIM protocol can be adapted to new ways to affect gene function. The mRNA depletion by transcription inhibition using the *Tet-off* system is particularly appealing, since this strategy should work for any essential gene.

While some results were specific to either GIM or SGA assays, thus being condition and assay-dependent, there are a number of direct GIs that were robustly detected in both types of screen. We thus generated a list of 479 pairs of genes having a synergistic negative impact on growth in both GIM and SGA screens (**Supplementary Table 5**). This list of GIs that were observed in very different assay condition represent a gold standard that could serve to benchmark future large-scale GI screen results.

The results presented here were thus validated by the correlation of GI profiles for the same gene mutation and by overlapping gene deletion analysis. In combination with previous SGA results, our data also validate several hundred GIs that can be considered robust.

### The amplitude of GIs for DAmP alleles correlates with specific gene features

Since the DAmP mutant strains were used in addition to gene deletion strains in our screens, we wondered if we could obtain insights into global differences between these two types of gene perturbation. DAmP perturbation of gene function depends on the effect of a long 3’ untranslated region on RNA stability. The NMD degradation pathway, responsible for destabilization of DAmP RNAs can be highly variable (Decourty *et al*, 2014; Breslow *et al*, 2008). Thus, its impact on the essential gene function, and the profile of GIs for the corresponding mutant, was likely to vary and could be correlated to various RNA features, such as abundance or coding sequence length. We thus looked for a correlation between original mRNA abundance for essential genes and the frequency at which the corresponding DAmP alleles showed a GI in the results. To this end, we arbitrarily defined *screen-responsive* gene perturbations as those in which we observed at least a variation by a factor of 2 for a given gene in at least one of the GIM screens. Mutants that showed no effect in combination with any of the 154 query gene perturbations would be included in the *non-responsive* category. We calculated which fraction of the tested mutants was in the *responsive* or *non-responsive* category in correlation with RNA abundance and coding sequence length.

Interestingly, the percent of *screen-responsive* DAmPs increased with the abundance of the corresponding mRNAs (**Fig. 4 A**). As background, and for comparison, we applied the same analysis to gene deletions, for which the effect of mRNA abundance on the frequency of response in GIM screens was less marked. However, in both cases deletion or DAmP perturbation were most correlated with an effect in GIM screens for the most abundant mRNAs. We also looked at the relation between screen responsiveness and the length of the coding sequence for the affected gene, which is linked with the destabilization of DAmP modified mRNAs (Decourty *et al*, 2014). In this case, the fraction of screen-responsive DAmP mutants decreased as the length of the initial gene coding sequence increased. This effect of coding sequence length was not found for gene deletions, where, on the contrary, the highest proportion of screen-responsive mutants was found in the group of long genes (**Fig. 4 B**). Thus, features associated with an effect visible in the GIM screen conditions were high expression level and large gene size for gene deletion and high abundance mRNA and short coding sequence size for DAmP modification.

**Figure 4.**
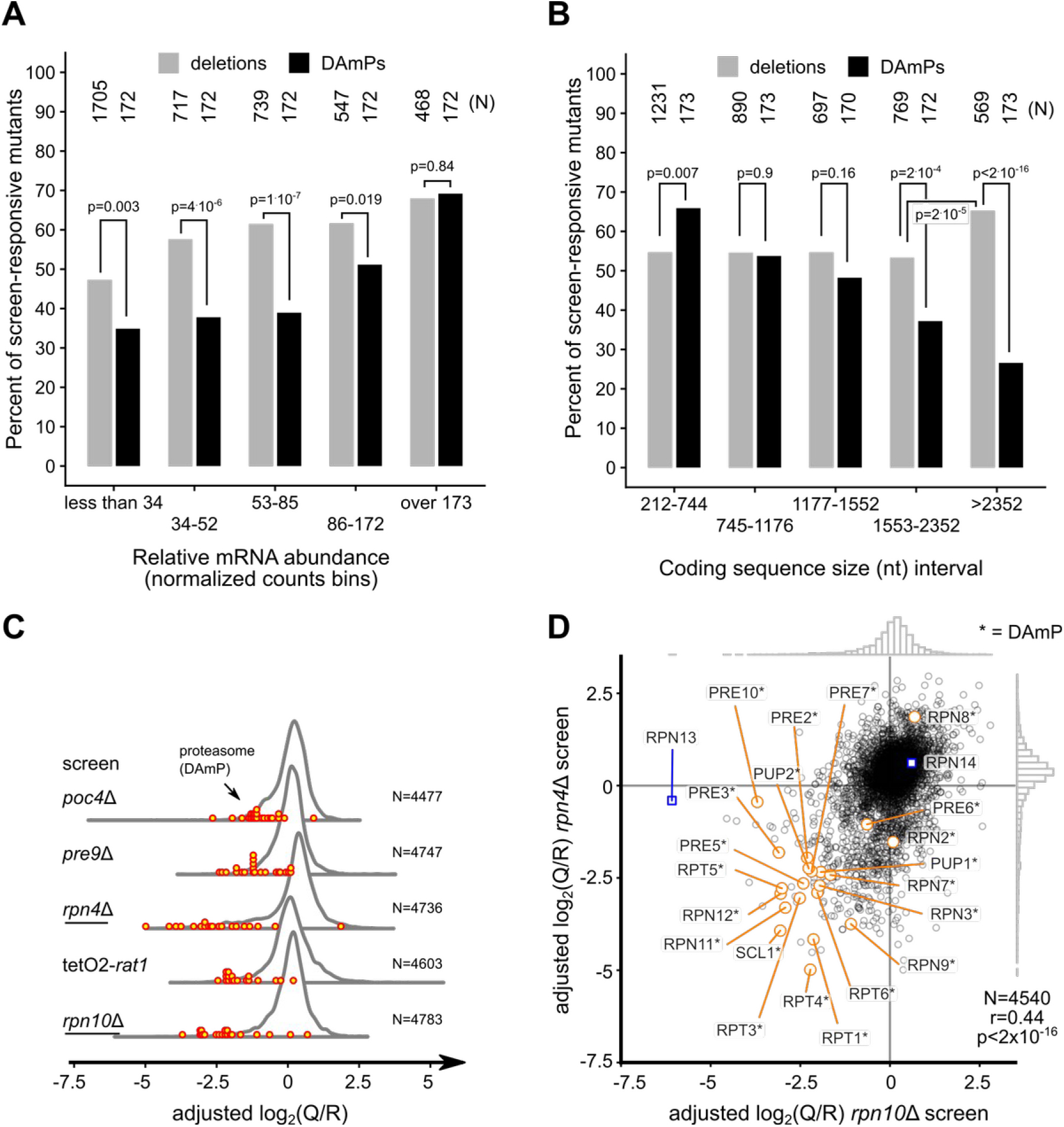
DAmP perturbation has effects correlated with mRNA abundance and coding sequence length and is valuable for the study of major cellular functions. **A**. We arbitrarily assigned the various mutants from this study in two classes: *screen-responsive*, if the corresponding mutant showed a growth defect score of at least 2 (log_2_(Q/R) < -1) in at least one of the 154 screens, and *screen-neutral* if the mutant was not affected in any of the screens. The percent of *screen-responsive* deletion (light gray) and DAmP (black) strains was plotted as a function of relative mRNA abundance (Lipson *et al*, 2009), with transcripts grouped in five bins having identical numbers of DAmP mutants. The differences between the numbers of DAmP and deletion mutants in each bin were evaluated with a chi-squared test (the p value for the null hypothesis of identical percentages is indicated). The number of genes in each bin is indicated in the upper part of the panel. **B**. Equal sized bins of DAmP mutants were created based on the coding sequence length and the percentage of *screen-responsive* strains was compared with the results for deletion mutants for genes having similar sizes of coding sequences. The number of mutants in each bin is indicated. **C**. We used the median of the relative rank for 22 DAmP mutants affecting proteasome and proteasome-related genes to identify the 5 screens in which these mutants were most affected (in increasing rank order from bottom to top). The distribution of all adjusted log_2_(Q/R) values in the five selected screens is indicated. Red dots indicate the position of the adjusted log_2_(Q/R) scores for proteasome DAmP mutants. **D**.Specific DAmP effects are illustrated by a scatter plot showing the correlation between the adjusted log_2_(Q/R) scores obtained when the screen was done with the deletion of the RPN10 proteasome component gene (horizontal axis) compared with the deletion of the RPN4 proteasome regulator (vertical axis). DAmP proteasome related mutants are indicated in orange and two non-essential proteasome gene deletions are indicated in blue.

To illustrate how useful the new results on DAmP strains can be and further validate the obtained result on a large scale, we focused on a group of 22 DAmP mutants affecting proteasome-related genes, which are highly expressed and can be relatively short. For example, 11 out of the 22 selected genes have coding sequences shorter than 1176 nucleotides, which places them in the first two bins represented in **Fig. 4 B**. We ranked the screens to find those in which the median of the adjusted log_2_(Q/R) values for this group of proteasome DAmP mutants was lowest. The top 5 screens showing GIs with proteasome-related genes were, in order, those using as query genes the deletion of RPN10, the depletion of RAT1, and deletions of RPN4, PRE9 and POC4. Four out of the 5 screens in which DAmP proteasome mutants were most affected corresponded thus to perturbation of proteasome components RPN10 and PRE9, of a regulator of proteasome formation, RPN4, and of the proteasome assembly factor POC4. Values for the DAmP proteasome mutants in those screens were clear outliers, when compared with the overall distribution of adjusted log_2_(Q/R) values in each screen (**Fig. 4 C**). In the screens performed with deletions of RPN10 and RPN4, DAmPs for proteasome-related genes represented the majority of strong negative measured GIs, as illustrated in **Fig. 4 D**.

A fifth screen showing a strong global effect in combination with proteasome DAmP mutants involved the temporary depletion of the major nuclear 5’ to 3’ exonuclease RAT1 using the *Tet-off* system. This result was surprising but compatible with the various roles of the proteasome in transcription (reviewed in Durairaj & Kaiser, 2014) and the described role of Rat1 in RNA polymerase II transcription termination (Kim *et al*, 2004). Alternatively, it could be an illustration of the deregulation of protein homeostasis following Rat1 depletion, which might require compensation by an increase in proteasome activity (Tye *et al*, 2019). Thus, the use of DAmP mutants in combination with the *Tet-off* strategy for query gene perturbation uncovered GIs that are functionally relevant and potentially important.

In addition to the proteasome analysis detailed in this section, DAmP modification led to the identification of other new links between genes involved in RNA metabolism. Thus, for example, the DAmP modification of the 3’ to 5’ RNA degradation exosome complex component RRP46, and of the rRNA modification complex component NOP56 were synthetic sick with the deletion of several RNA-related genes (**Fig. 4 – figure supplement 1**), including the poorly characterized locus YCL001W-A and the recently identified SKI complex associated protein Ska1 (Zhang *et al*, 2019). Altogether, these results illustrate the value of including mutants of essential genes in GIM screens.

### Predicting function based on GI profiles by using RECAP

Associating genes with a cellular pathway is often based on the observation of a specific phenotype when the gene function is affected by deletion, down-regulation or mutation. A different type of phenotype, tested in large-scale genetic screens, is represented by the constellation of gene perturbations that, when combined with a yeast gene mutant of interest, have an effect on the strain’s growth rate. This profile of response of a mutant to combinations with the query gene perturbation, also called GI profile, can be used to find functional relationships from screens data (Schuldiner *et al*, 2005; Decourty *et al*, 2008; Costanzo *et al*, 2010). GI profile similarity is an important type of results derived from large-scale genetic screens. We wondered whether we could use GI profile similarity from our results and integrate it with curated literature data on yeast genes to understand : a) what fraction of known functional interactions can be reached with our GI data set and b) whether we can use the similarity of GI profiles to assign new genes to known cellular pathways. To answer these questions, we developed a data analysis strategy called **RECAP** (**R**ational **E**xtension of **C**orrelated **A**nnotations and GI **P**rofiles, summarized in **Fig. 5 – figure supplement 1**), which, instead of focusing on the GI network, starts from published data curated by the Saccharomyces Genome Database maintainers (Cherry *et al*, 2012).

**Figure 5.**
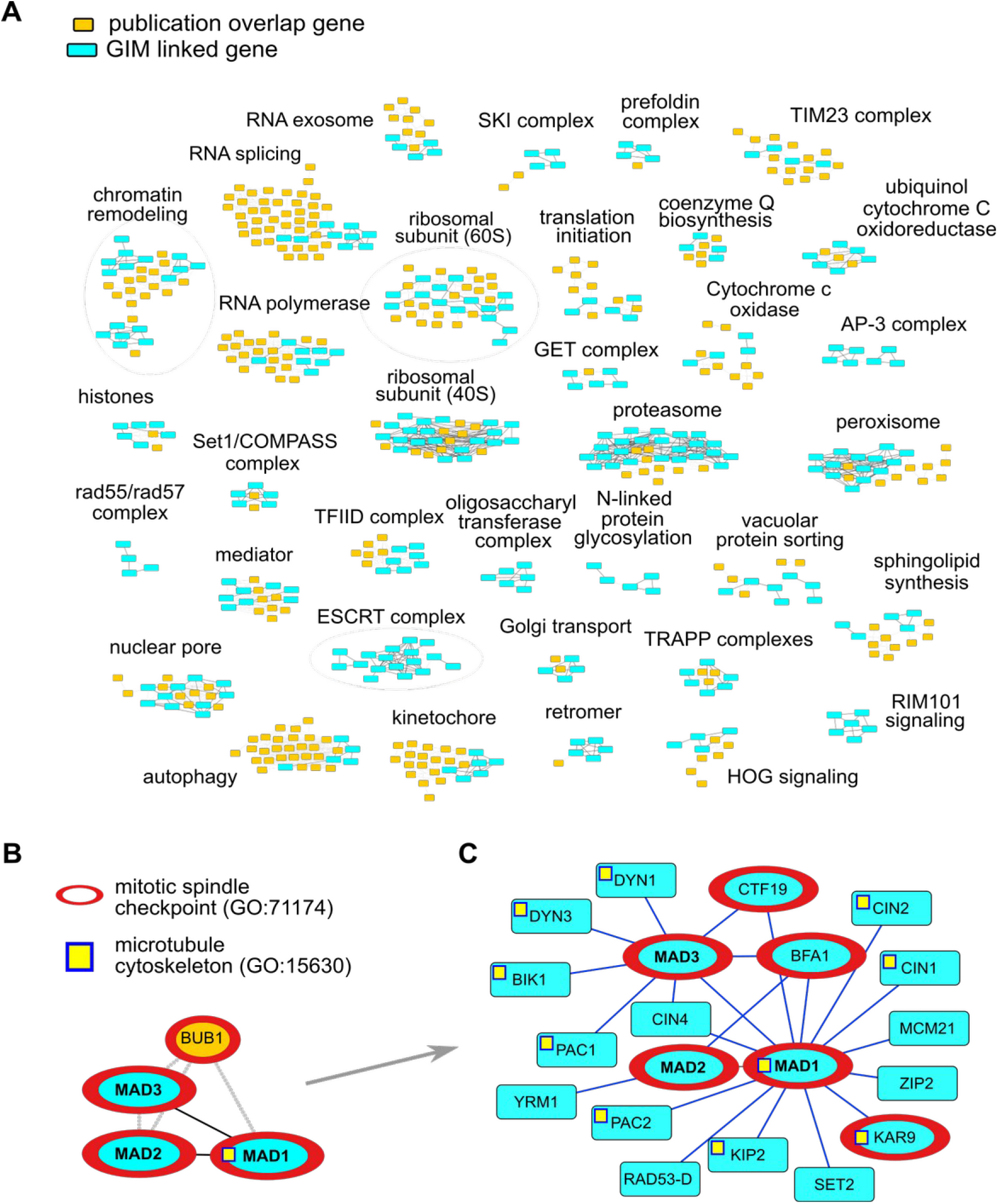
Integration of literature data and GIM profile similarity allows extension of known functional networks. **A**. Publications and linked genes were recovered from the *Saccharomyces* Genome Database and used to define highly connected factors. Gene pairs highlighted in cyan correspond to situations in which the corresponding GI profiles were correlated. Only a selection of 35 gene groups in which at least 4 genes showed correlated GI profiles is shown. Each gene group was annotated manually, either in terms of a protein complex or based on known cellular or molecular function. **B**. Example of a literature-based gene group, not shown in A, bringing together several genes involved in mitotic spindle checkpoint. Extending this network using only the similarity of GIM profiles led to the network shown in C. **C**. Starting from MAD1, MAD2 and MAD3, the GI similarity-based functional network adds supplementary genes involved in mitotic spindle checkpoint, such as KAR9, BFA1, and CTF19 (marked with a red border) and genes involved in the dynamics of the microtubule cytoskeleton (marked with a yellow square).

To establish links between genes from the table that associates genes and publications, we first removed publications associated with more than 100 genes, since we considered that such publications are too general to be informative. The remaining literature corpus consisted of 76 160 publications. We restricted our analysis to the upper half of the most cited yeast genes, leading to a selection of 3 575 genes or genomic features cited in at least 31 scientific publications. Among these well studied genes, 1 847 were present in our GIM data set of 5 063 genetic interaction profiles. For these 1 847 genes, we identified 4 072 linked gene pairs. We considered two genes, A and B, to be linked by co-citation if, for each gene A, at least 20 % of its publications mentioned gene B and reciprocally, if 20 % of citations for B also contained A. We used the Louvain algorithm (Blondel *et al*, 2008) on the set of 4 072 gene pairs to identify 439 communities of related genes corresponding either to well-studied complexes or to well-known genes involved in the same cellular pathway (**Supplementary Table 6**).

In the next step of the RECAP approach, we wanted to combine the newly defined literature-based clusters with the information available from the similarity of GI profiles in the GIM data. We calculated Pearson correlation for each pair of the 5 063 GI profiles of our data set. For each mutant, we sorted the obtained correlation coefficients in decreasing order and arbitrarily considered two mutant profiles, X and Y, to be linked if the correlation coefficient of X with Y and of Y to X were in the top 20 of correlation coefficients for both X and Y. This choice increased the specificity of the method and avoided situations in which spurious correlations would pollute the results.

Having established literature and GIM-based links between genes, we wondered how to combine these results. Among the 439 communities of related genes identified from the literature, 117 had at least two genes linked by GI profiles similarity in our GIM data set (**Supplementary Table 7**). To visualize the presence of data that matched the GI profiles we selected the 35 groups, out of 117, that had at least 4 genes from the GIM data showing correlated profiles to other genes from the same sub-network (**Fig 5A**). These groups comprised 550 genes (of the 1 847 genes of the network) and 2 393 links and involved data for 270 genetic interaction profiles (of the 5063 available), including those of 67 DAmP mutants. The cellular functions that corresponded to the 35 groups of genes covered a wide range, from DNA transcription to vesicular transport and mitochondrial function (**Fig. 5 A, see annotations**).

Our literature-based analysis of genetic interaction profiles indicated which mutant strains had phenotypes specifically correlated with the function of the corresponding gene. This knowledge allows focusing on these mutants first, since they were independently validated to provide functional information. Using this knowledge is essential to avoid conclusions about gene function that would come, for example, from perturbing an unrelated gene that is physically close on the chromosome (Atias *et al*, 2016). We thus used the validated GI profiles from each of the 117 co-citation based clusters in which at least two genes had similar GI profiles and used GI similarity to extend each of the clusters. Importantly, if genes for which we had profile data were present in co-citation clusters but were not linked by GI profile similarity to other genes in the same cluster, these profiles were ignored.

An example of the performance of this approach is shown in **Fig. 5** for the group of literature-linked genes MAD1, MAD2, MAD3 and BUB1 (a paralogue of MAD3). The MAD genes contribute to the spindle assembly checkpoint in relation with kinetochores, and are required for cell division (reviewed in Yamagishi *et al*, 2014). This group of four genes included the correlated GI profiles for the MAD1/ MAD2 and for the MAD1/MAD3 pairs (**Fig. 5 B**). The RECAP-extended network based on the MAD gene group included other genes with roles in the spindle assembly checkpoint, such as KAR9, CTF19 and BFA1, but also many genes related with microtubule cytoskeleton organization and function, a process that is linked directly with spindle assembly and function (**Fig. 5 C**). A few genes were not annotated directly to spindle assembly or microtubule function, such as RAD53 (DAmP modification), SET2, ZIP2 or MCM21. However, MCM21 is a component of the COMA kinetochore sub-complex (De Wulf *et al*, 2003), RAD53, a gene with multiple roles in DNA repair, is also linked with mitotic checkpoints (reviewed in Lanz *et al*, 2019) and ZIP2 is involved in homologous chromosome pairing in meiosis (Chua & Roeder, 1998). Thus, the large majority of the genes identified by RECAP starting from just a few components of the spindle assembly machinery had functions in relation with this process.

We used GO term enrichment analysis (Raudvere *et al*, 2019) to establish the biological process and cellular component that were predominant in each of the 117 literature-defined communities and associated the genes from the extended RECAP network to these processes (**Supplementary Table 8**). A total of 1471 genes involved in 3893 gene pairs were finally associated with defined processes, allowing new hypotheses about these genes function to be tested by future oriented experiments. The RECAP strategy is not limited to GI profile similarity, but can be adapted to other large-scale data sets in which links between related genes have been established.

### Linking inositol polyphosphate metabolism and translation initiation

In addition to GI profile similarity, the discovery of individual GIs can also be informative for gene function. We explored in detail the synthetic lethality between the deletion of LOS1, a gene involved in tRNA export from the nucleus to the cytoplasm (reviewed in Hopper, 2013), and several OCA genes. OCA1, the founding member for the OCA nomenclature, was initially identified as an Oxidant induced Cell-cycle Arrest factor (Alic *et al*, 2001). The other five members of this protein family were identified based on protein sequence similarity (Wishart & Dixon, 1998; Romá-Mateo *et al*, 2011). Only recently a biochemical role was attributed to OCA3 in the hydrolysis of specific inositol-polyphosphate species (Steidle *et al*, 2016). Modification of inositol-polyphosphate levels in OCA mutants is probably responsible for the observed phenotypes when OCA genes are deleted, ranging from changes in replication of an RNA virus (Kushner *et al*, 2003) to effects on yeast prion propagation (Wickner *et al*, 2017).

We have previously observed a strong growth defect when OCA2 and LOS1 deletions were combined (Decourty *et al*, 2008). This link was confirmed when the deletion of OCA4, another OCA gene, was tested by SGA, although LOS1 deletion was only found as the 101st most affected hit (Costanzo *et al*, 2016). GI profiles for deletions of OCA1, OCA2, OCA3 (SIW14), and OCA5 were highly similar in the SGA data (Costanzo *et al*, 2016). All the OCA deletion mutants also showed a coordinated response to a set of chemical compounds (Hoepfner *et al*, 2014), indicating that loss of these proteins leads to a similar cellular response. In view of the similarity between OCA deletion profiles, we compiled physical interaction results about OCA proteins from the BioGrid database (Oughtred *et al*, 2019). Except for Oca6, we found evidence for physical interactions between OCA proteins, potentially in a multimeric complex (**Fig. 6 – figure supplement 1**).

**Figure 6.**
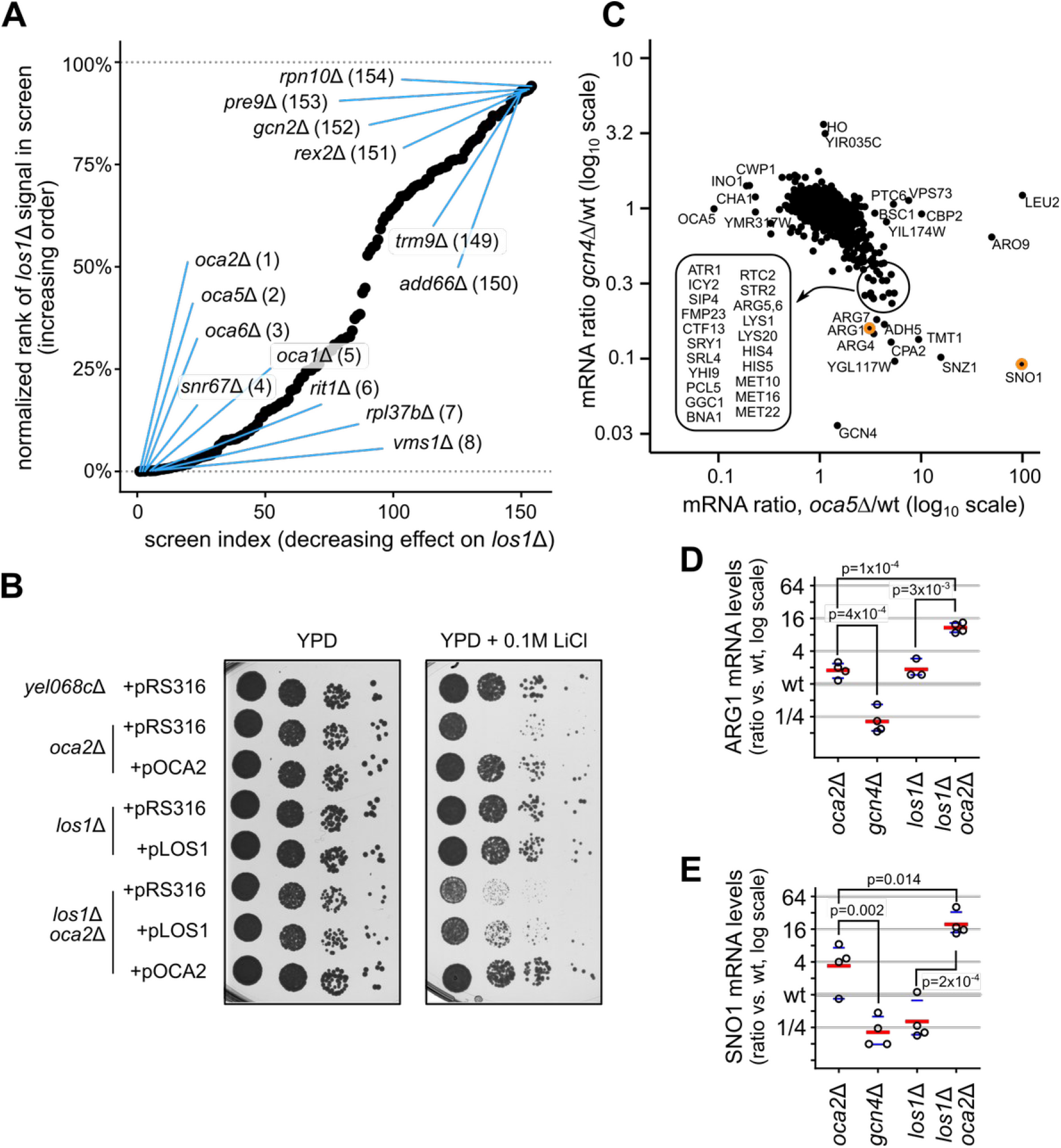
Deletion of LOS1 is functionally related to a defective OCA complex. **A**. Rank analysis of *los1*Δ results for GIM screens highlighting synthetic slow growth (lower left) and potential epistasis (upper right). **B**. The double deletion strains combining *los1*Δ and *oca2*Δ were strongly affected by the presence of 0.1M LiCl in the medium. Complementation of growth defect by empty vector (pRS316) or by centromeric plasmids expressing OCA2 and LOS1 was estimated by serial dilutions and observation of colonies after 48 hours of growth at 30°C. **C** The inverse correlation between the transcriptome changes in *oca5*Δ and *gcn4*Δ (Hughes *et al*, 2000) shows transcripts that were up-regulated in the absence of OCA5, while being targets of GCN4 activation, including many mRNAs that code for amino acid biosynthesis proteins. The position of the signal for mRNA of ARG1 and SNO1, chosen for validation of the transcriptome results, are indicated by orange dots. **D and E**. Validation by RT-qPCR of mRNA level changes in an *oca2*Δ strain, in comparison with *gcn4*Δ, *los1*Δ, and the combination *oca2*Δ/*los1*Δ for ARG1 and SNO1 mRNA, using RIM1 mRNA levels as reference. Individual measurements for 3 to 4 independent experiments are shown, with the red bar indicating the mean and the blue bars indicating limits of the 99% confidence interval (non-parametric bootstrap). The indicated p-values correspond to results of single sided t-tests.

In correlation with the previous data on OCA genes and LOS1, we found that LOS1 deletion showed, among our 154 GIM screens, the strongest negative effect on growth when combined with deletions of OCA2, OCA4, OCA6 and OCA1 (**Fig. 6 A**). The sixth screen in which deletion of LOS1 strongly affected growth involved the deletion of RIT1, which modifies initiator tRNA and renders it incompetent for translation elongation (Aström & Byström, 1994). This result was compatible with the role of LOS1 in tRNA export, including tRNA_i_^Met^ and suggested that the other observed GIs for LOS1 had high confidence. To further validate the GIs between OCA genes and LOS1 deletion, we tested the growth of single and double mutant strains in various media. We found that moderate doses of lithium chloride in rich medium sensitized the growth assay for the *los1*Δ/*oca2*Δ strain. Under these condition, expression of LOS1 and OCA2 from plasmids (MOBY collection, Ho *et al*, 2009) partially restored growth of the double mutant strain, thus confirming the screen results (**Fig. 6 B**).

To further explore OCA genes role and how they could be connected with LOS1 function, we analyzed results obtained during large-scale transcriptome profiling of deletion mutants (Hughes *et al*, 2000). Among the tested deletion mutants, the data set contained the transcriptome measures for the effects of deleting OCA5. We noticed that the transcriptome changes in this OCA5 mutant were inversely correlated with those observed when the translation-regulated transcription factor GCN4 (or its overlapping gene YEL008W) were deleted.

GCN4 is one of the best studied example of translation regulation and plays a crucial role in adaptation of yeast cells to amino acid starvation (reviewed in Hinnebusch, 2005; Hinnebusch *et al*, 2016). Activation of GCN4, whose mRNA contains four short open reading frames upstream the start codon, occurs when translation is inhibited and leads to transcription of hundreds of targets, including many genes involved in amino acid synthesis (Natarajan *et al*, 2001). Such GCN4 targets were responsible for the strong inverse correlation between the transcriptome results in the *gcn4*Δ strain compared with *oca5*Δ (Hughes *et al*, 2000, **Figure 6 C**). To validate this observation, we measured the changes in the levels of two representative transcripts, ARG1 and SNO1, in strains deleted for OCA2 and GCN4, by reverse-transcription and quantitative PCR. Correlated with the published results, OCA2 absence led to an increase, while GCN4 absence led to a decrease in their levels (**Fig. 6 D and E**). The increase in the levels of these transcripts in the absence of OCA2 was further enhanced in double mutant strains combining the deletion of OCA2 with the deletion of LOS1 (**Fig. 6 D and E**). Altogether, these results suggested a link between OCA genes and translation regulation that was potentiated in the absence of the tRNA export factor LOS1.

A possible explanation for the observed results was that the loss of OCA genes leads to GCN4 activation. To test this hypothesis, we used a reporter system in which beta-galactosidase is expressed in a GCN4-like configuration, with its coding sequence fused with the 5’ untranslated region of GCN4 (Mueller & Hinnebusch, 1986). In this system, we observed a clear increase in the beta-galactosidase activity when LOS1 deletion was present in an *oca2*Δ strain (**Fig. 7 A**). Thus, the perturbation of OCA function coupled with a tRNA export deficiency led to GCN4 activation, most likely through inhibition of translation initiation.

**Figure 7.**
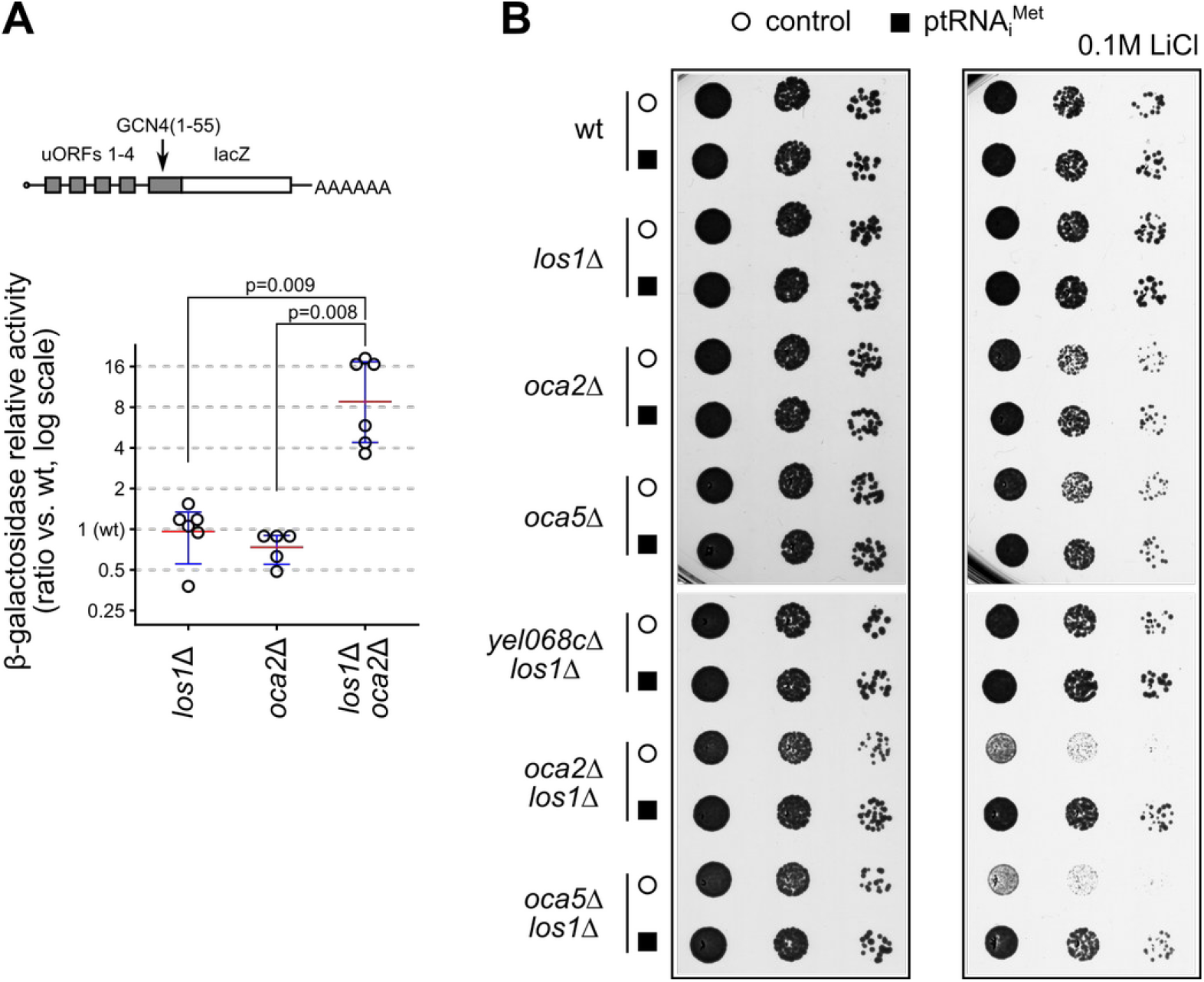
Initiator tRNA limits cell growth when the OCA complex is defective. **A**. Double mutant strains show translation initiation defects, as measured using a GCN4 uORF-lacZ reporter, schematically represented in the upper part of the panel (p180 plasmid, Mueller & Hinnebusch, 1986). Wild type strain was used as reference and the variation in the amounts of produced beta-galactosidase were measured in at least 5 independent experiments. The p-values of single-sided t-tests for differences between the different conditions are indicated. **B**. Over-expression of the tRNA_i_^Met^ (p1775 plasmid, Dever *et al*, 1995) allows better growth of *oca2*/*los1* and *oca5*/*los1* double deletion strains under stress conditions (0.1M LiCl). Serial dilutions of fresh cells were grown on plates for 48 hours. The presence of an empty vector (empty circle) or of the plasmid over-expressing the tRNA_i_^Met^ (black square) are indicated.

A central factor in translation initiation is the tRNA_i_^Met^ and, since LOS1 is involved in its nuclear export, we wondered if it could be involved in the synthetic sick effect observed when combining LOS1 and OCA gene deletions (**Fig. 6 A** and **B**). Over-expression of tRNA_i_^Met^ (Dever *et al*, 1995) led to a reversal of the slow growth phenotype of both *los1*Δ/*oca2*Δ and *los1*Δ/*oca*5Δ strains (**Fig. 7 B**), indicating that initiator tRNA shortage becomes limiting in the absence of OCA genes. How the inositol-polyphosphate imbalance generated in such strains affects translation initiation remains an interesting question for future research.

## Discussion

The set of about 700 000 GIs described in our study, together with multiple validations of the obtained results (**Fig. 2, 3**), establish a new resource for functional genomics in yeast, along and complementary to previous large-scale GI results (for example, Costanzo *et al*, 2010, 2016). Individual screens performed with temporary transcription repression of query genes also demonstrated the value and flexibility of GIM screens for the study of essential gene function (see, for example the case of RRP6, **Fig. 2D** and RAT1, **Fig. 4A**). One of the advantages of GIM screens is that they do not require any robotic devices. The detection of barcodes, originally done with DNA microarrays can be switched to DNA sequencing, as shown for chemogenomic screens (Smith *et al*, 2009). Thus, GIM screens are a powerful alternative to SGA for large scale GI tests. The results presented here identify novel GIs for essential and non-essential genes involved in RNA metabolism in yeast and bring an independent validation for hundreds of previously observed GIs. As demonstrated through the analysis of the correlated GI profiles, this new data set explores a large variety of cellular processes and macromolecular complexes, well beyond the function of the 154 screen query genes (**Fig. 5**).

One of the goals of performing large-scale genetic screens is to establish new functional links between genes and cellular processes. To this end, gene set enrichment analysis (Subramanian *et al*, 2005) can be applied to groups of genes that share similar GI profiles. A refinement of this approach, as implemented in the spatial analysis of functional enrichment (SAFE) method (Baryshnikova, 2016), includes in the enrichment analysis the topology of the gene network, built, most frequently, from the similarity of GI profiles. This method has the advantage of providing a map for how various enriched GO terms distribute across network and allows a visually rich inspection of the results. We demonstrate here a complementary approach, called RECAP, that combines gene co-citation links with GI profile similarity and identifies pairs of genes that are both related by the literature data and by the experimental results (overview of the method in **Fig. 5 – figure supplement 1**). The inclusion of literature information in the analysis of GI profiles highlights mutants that behave as expected in the GI data set. This selection process validates hundreds of GI profiles and allows the identification of linked genes and their association with well described biological processes or macromolecular complexes.

The originality of RECAP consists in the use of published results to find GI profiles of high confidence, serving as anchoring points to extend the network of functional links. The reason we used co-citation to build the initial functional network is that it suggests links between genes and groups of genes in a manner that is quite natural and independent from the hierarchical gene ontology terms annotations (Ashburner *et al*, 2000; The Gene Ontology Consortium, 2019). Co-citation is used by major gene annotation and protein interaction databases such as STRING (Szklarczyk *et al*, 2017). However, it has not been used until now for the analysis of GI networks as a yardstick in the initial validation of experimental results.

A potential problem of co-citation is that its quality depends on the availability of a high-quality curated database that associates genes and publications, such as the one maintained by the Saccharomyces Genome Database (Cherry *et al*, 2012). Since the version of RECAP presented here depends on such manually curated database, extending it to other organisms depends on the presence of equivalent resources. It is likely that full text mining, such as the one implemented by Textpresso (Müller *et al*, 2018) would allow automatic building of such databases for other organisms. Alternatively, association by GO term similarity and by other ways to link genes, such as the network extracted ontology (Dutkowski *et al*, 2013), could be also effective as the first step of a RECAP analysis.

RECAP is useful to confidently extend functional interaction networks with GI results (for an example see **Fig. 5 B** and **C**). Among the mutants validated by this analysis we were particularly interested in those affecting essential genes through the DAmP modification, since functional interactions, as detected by large-scale genetic screens, are scarcer for essential than for non-essential genes. Many DAmP strains display normal growth rates under standard culture conditions (Yan *et al*, 2008; Breslow *et al*, 2008). However, testing a few conditions might miss specific phenotypes associated with DAmP perturbation of genes. The GIM screens performed in this study are equivalent to testing 154 different stress conditions for each of the tested DAmP strain. Similar to the observation that deletion of most non-essential genes does not affect growth under standard culture conditions, but can be limiting in the presence of a chemical (Hillenmeyer *et al*, 2008), we view the set of GIM screens we performed as a series of highly diverse stress conditions. Thus, the screens probed a panorama of conditions for the 900 DAmP mutant strains and allowed the identification of global trends, such as the correlations of DAmP modification effect with short coding sequence length and high gene expression (**Fig. 4**). Together with the previously published results on DAmP mutants (Costanzo *et al*, 2010, 2016), the large-scale characterization of these collections of strains is a useful resource for anyone interested in the study of specific essential genes.

In addition to the 154 different tests for each of the approximately 5 000 tested mutants, the presence of translation inhibitors in the GIM screens introduced an additional stress common to all our screens. This stress allowed the identification of GIs that would have been otherwise of much lower amplitude or undetectable. An example is the link between the OCA complex genes and the tRNA export factor LOS1, which was weak in SGA results (Costanzo *et al*, 2016), but among the highest-ranking ones in the GIM data (**Fig. 6 A**). We validated this link on individual double mutant strains and showed that it is dependent on the availability of the initiator tRNA_i_^Met^ (**Fig. 6 and 7**). Since the OCA complex affects inositol polyphosphate metabolis, this result adds a new element in the complex puzzle of the influence of inositol poly-phosphates on cellular processes and highlights the usefulness of measuring genetic interactions under a variety of conditions, as previously suggested (Martin *et al*, 2015; Jaffe *et al*, 2019).

The new GI resource together with RECAP and the associated validation experiments will be useful for further exploration of gene function in yeast and other organisms.

## Materials and Methods

**GIM screens** were performed as described originally (Decourty *et al*, 2008), and following the protocol described in detail in (Malabat & Saveanu, 2016), using custom-made turbidostat devices that allowed performing 16 cultures in parallel. Briefly, **MATα** query strains were obtained by replacing the KanMX resistance cassette in strains from the gene deletion collection (Giaever *et al*, 2002) with a Prα-Nat cassette that expresses the nourseothricin resistance gene only in the context of a haploid **MATα** strain. Hygromycin B resistance was also added to the query strain using a centromeric plasmid to allow selection of diploid strains after mating. Pools of deletion (Giaever *et al*, 2002) and DaMP strains (Decourty *et al*, 2014) were recovered from stock maintained at -80°C and left to recover in rich medium for 30 minutes by incubation at 30°C, then mixed with fresh query strain culture for mating on a plate. Recovered diploids were incubated overnight at 30°C in the presence of 0.2 mg/ml hygromycin B and 0.2 mg/ml G418. Sporulation was induced after culture in GNA medium (5% glucose, 3% Difco nutrient broth, 1% Difco yeast extract) by switching to potassium acetate medium (1% potassium acetate, 0.005% zinc acetate, supplemented with 2 mg uracil, 2 mg histidine and 6 mg leucine for 100 ml). After sporulation, cells were recovered in YPD medium (2% glucose, 1% Difco yeast extract, 1% Difco Bactopeptone), incubated for 6 hours without antibiotics and grown for 45-60 hours in the presence of 0.2 mg/ml G418 and 20 μg/ml nourseothricin. For each batch of 16 screens, the reference against which the screens were compared was a mix of cells from all the final cultures. For *Tet-off* query strains and screens, doxycyclin at 10 μg/ml was included in the dual antibiotic haploid selection step for 16 to 24 hours in liquid culture. DNA was extracted from the final cell pellets and used to amplify upstream and downstream barcodes. Barcode DNA relative levels were measured using custom microarrays (Agilent Technologies, California, USA) and the collected images were processed with GenePix Pro 6 (Molecular Devices, California, USA) and analyzed using R (R Core Team, 2019). Data analysis consisted of normalization of the Cy3/Cy5 using the loess algorithm, aggregation of results for upstream and downstream barcodes and normalization of the aggregated results. Each screen result was examined for the presence of the expected signal around the query gene locus that corresponds to the decrease in recombination frequency during meiosis due to physical proximity on the same chromosome. Situations with secondary peaks or lacking exclusion peaks were eliminated from further analysis. The exclusion peaks were corrected using estimates of recombination frequency based on the observed signal (Decourty *et al*, 2008). Finally, results from at least two independent screens were expressed as the log_2_ of the ratio between the screen of interest and the reference (Q/R) and combined to obtain GI estimates. Results were corrected for pleiotropic effects by counting the fraction of screens in which a given mutant showed a log_2_(Q/R) value inferior to the arbitrary threshold of -1.25, named pleiotropic index (PI). Each initial log_2_(Q/R) value was multiplied with (1-PI)^3^ to decrease the weight of mutants showing a response in most screens. Adjusted values of log_2_(Q/R) were used to compute Pearson correlation coefficients for all the GI profiles pairs. To assess the reciprocity of the observed GI profiles, we ranked for each mutant the similarity of profiles for all the other mutants in decreasing order of the corresponding Pearson correlation coefficient. If the GI profile for mutant A was among the top 20 profiles for mutant B and, conversely, the profile for mutant B was in the top 20 profiles for mutant A, we considered that A and B were linked.

### RECAP data analysis

To annotate the observed GI profile links we used the curated database of yeast literature from the *Saccharomyces* Genome Database (Cherry *et al*, 2012). The table associates genes with publications. Only articles dealing with less than 100 genes were selected, and only the half most cited yeast genes were used to build a network of co-citations. Links in this network were based on the presence of the two genes in the same publications. At least 20 % of articles citing a gene had also to cite the other one to establish a connection. The obtained co-citation network for 1 847 genes showed strong connections among 439 isolated groups of genes. Within each group, we analyzed the presence of genes linked by GI profile similarity and selected 117 cases in which at least one such connection was present. Genes connected by GI profile similarity and by co-citation were considered valid in terms of GIM screens and served to extend the network using the current GIM data set of adjusted log_2_(Q/R) and the computed links based on reciprocal GI profile similarity. The links based on GI profile similarity were then used to associate genes with biological processes or cellular components. To this end we used the list of genes of each group to interogate the g:Profiler web server https://biit.cs.ut.ee/gprofiler/gost for gene ontology term enrichment, using the *gprofiler2* R package (Raudvere *et al*, 2019) and selected the top entry for biological process and cellular component in each case. To test various configurations for RECAP, we used the R *igraph* (Csardi & Nepusz, 2006) and *RCy3* (Gustavsen *et al*, 2019) packages, together with visualization and network analysis in Cytoscape (Shannon *et al*, 2003).

### Strains and plasmids

The generation and details of the DAmP strains collection were previously published (Decourty *et al*, 2014). Briefly, DNA from diploid strains from the deletion collection (Giaever *et al*, 2002) for the genes of interest was used to amplify the Kan^R^ cassette flanked by upstream and downstream barcode sequences. The cassette was amplified using oligonucleotides that targeted the insertion of the cassette in the genome of a BY4741 yeast strain downstream the stop codon of the same gene. Individual clones from each transformation were tested by PCR amplification with specific oligonucleotides. For the situations when the DAmP strain was used to perform GIM screens, the Kan^R^ cassette was transferred by amplification and transformation in the BY4742 **MATα** background. In these strains, the G418 resistance cassette was next replaced by the Prα-Nat cassette. For the *Tet-off* strains, we used the pCM224 vector (Bellí *et al*, 1998) to amplify the G418 resistance cassette and place the tetO2 sequence upstream the start position for the coding sequence of selected genes. Individual clones were tested by PCR on genomic DNA to test the correct insertion of the cassette. G418 resistance was next changed to **MATα**-specific nourseothricin resistance using the pGID3 vector (Decourty *et al*, 2008; Malabat & Saveanu, 2016). SnoRNA deletion strains for box H/ACA snoRNPs were derived from yeast strains available in our laboratory (Torchet *et al*, 2005). For box C/D snoRNAs deletion strains, we used the KanMX6 cassette from the pFA6 vector (Longtine *et al*, 1998) with the oligonucleotides listed in **Supplementary Table 9**. Double deletion mutant strains were built by mating of G418-resistant and nourseothricin resistant strains. After sporulation and selection of haploid clones, we ensured that the obtained strains had the same panel of auxotrophy markers as the parental strain BY4742. The strains for individual validation of screen results are listed in **Supplementary Table 10**.

### Beta-galactosidase activity

was measured on total cell extracts obtained by lysis by vortexing with glass beads. The assay buffer contained 0.5 mM chlorophenol red-β-D-galactopyranoside (CPRG), 100mM sodium phosphate, 10 mM KCl, 1 mM MgSO_4_, 5 mM dithiotrhreitol (DTT), pH 7. After incubation at 37°C, the absorbance change at 574 nm was normalized to protein concentration measured using the Bradford assay. The reported values are relative to the beta-galactosidase activity of a wild-type strain processed in parallel.

### RNA and RT-QPCR

Cells were grown in synthetic complete medium to log phase and collected. Total RNA was obtained using hot phenol extraction and DNA was removed with DNase I (Ambion TURBO DNA-free kit) before reverse-transcription (RT) and PCR amplification. For each experiment, 500 ng of total RNA were used in a RT reaction with Superscript III (Invitrogen) using a mix of the following oligonucleotides: SNO1rv AAC TCC TGA GGA TCT AGC CCA GTG, ARG1rv ACC ATG AGA GAC CGC GAA ACA G, and RIM1rv ACC CTT AGA ACC GTC GTC TCT C. Quantitative PCR reactions used the same oligonucleotides coupled with the following forward primers (one pair for each target): SNO1fw AAC TCC TGA GGA TCT AGC CCA GTG, ARG1fw GCA AGA CCT GTT ATT GCC AAA GCC and RIM1fw GCG CTT TGG TAT ATG TTG AAG CAG. For each experiment, the ARG1 and SNO1 signal was normalized to the RIM1 signal and all the results were compared with the wild type strain.

## Data availability

Raw and normalized microarray data were deposited in GEO (GSE119174, 312 samples), and ArrayExpress (E-MTAB-7191, 16 samples). Aggregated, normalized and pleiotropy adjusted results, including correlations of GI profiles can be explored at http://hub05.hosting.pasteur.fr/GIM_interactions/.

## Supporting information

Exclusion peaks correction for all screens

## Acknowledgments

We thank Lucia Oreus and Antonia Doyen for their contribution to performing the GIM screens, Thomas Moncion for early work on data analysis, Gwenaël Badis-Breard, Micheline Fromont-Racine and Frank Feuerbach for critically reading of the manuscript and our other colleagues from the GIM laboratory for help, advice, discussions and criticism. We thank Jean-Yves Coppée, Institut Pasteur, for his assistance in microarray hybridization and scanning. We thank Alan Hinnebusch and Thomas Dever (National Institutes of Health, Bethesda, MD, USA) for sharing plasmids and antibodies. This study was supported by the ANR GENO-GIM and CLEANMD grants (ANR-08-JCJC-0019, ANR-14-CE10-0014) from the French “Agence Nationale de la Recherche” and by continuous financial support from the Institut Pasteur and Centre National de Recherche Scientifique, France.

## Author contributions

A.J. and C.S. designed the experiments, L.D. and C.S. performed experimental work, C.M. and C.S. analyzed the data, E.F., L.D., A.J. and C.S. developed the multi-turbidostat device used for the GIM screens. C.S. prepared the figures, including the web visualization of results, and wrote the manuscript.

## Supplementary data synopsis

**Supplementary Table 1**: Query genes classification and features (for Fig. 1).

**Supplementary Table 2**: List of the 326 performed screens.

**Supplementary Data Set 1**: Images representing meiosis-dependent exclusion peaks for log_2_(Q/R) before and after correction for all the screens.

**Supplementary Table 3**: List of pleiotropy index (PI) values for 5063 tested gene mutants.

**Supplementary Table 4**: Pleiotropy corrected log_2_(Q/R) values for 5063 mutants in 154 screens.

**Supplementary Table 5**: A list of 479 pairs of synthetic slow growth interaction present in SGA and GIM screens.

**Supplementary Table 6**: Communities for 1847 genes grouped by their co-occurrence in publications.

**Supplementary Table 7**: Annotation of 117 gene communities containing GI profile similarity results.

**Supplementary Table 8**: Association of genes to biological processes and cellular components based on GI profile similarity and clusters of related genes.

**Supplementary Table 9**: Oligonucleotides used for snoRNA deletion strains construction.

**Supplementary Table 10**: *S. cerevisiae* strains used in this study.

---

**Figure 1– figure supplement 1.**
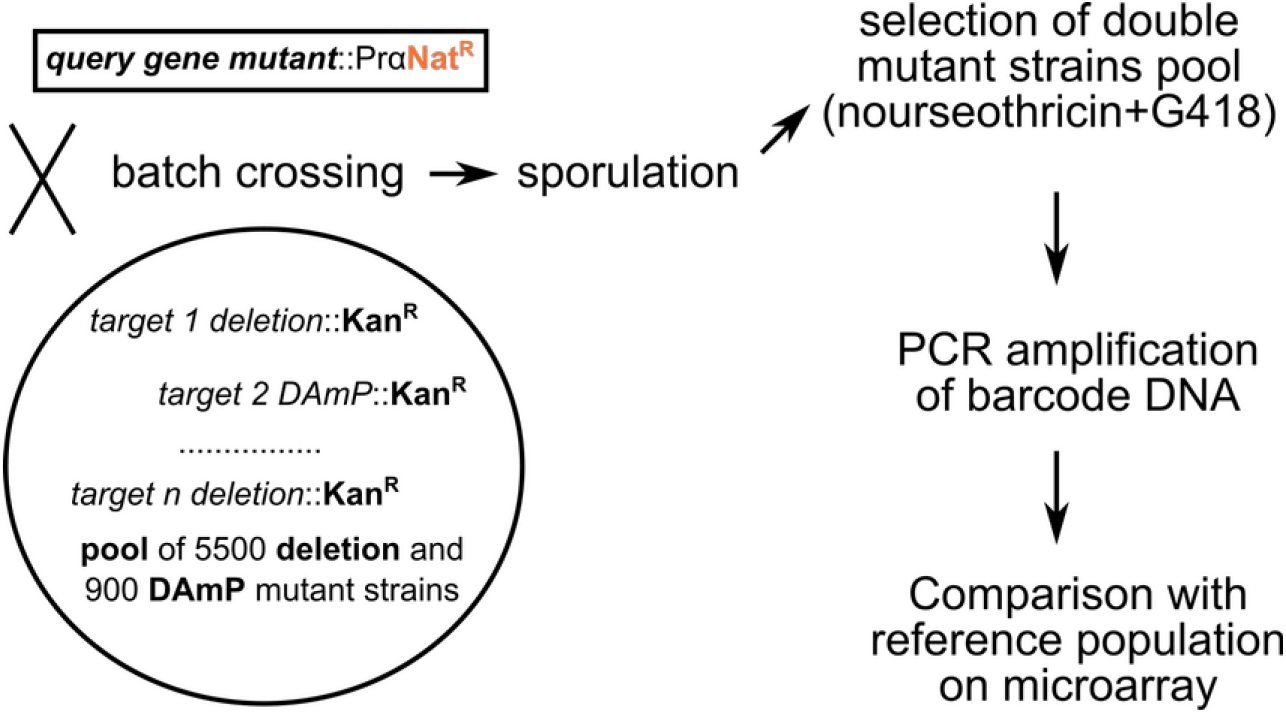
Overview of the GIM method (Decourty *et al*, 2008), in which a pool of double mutant diploid strains is obtained by crossing a MAT**α** query strain with a pool of MAT**a** mutants. Sporulation of the obtained diploid yeast strains is followed by the selection of haploid double mutants by using two antibiotics, one that selects for the haploid MAT**α** cells and the initial query mutation and another that selects for the mutation present in the initial pool of target genes. Amplification of barcode DNA from the population of double mutant strains is followed by comparison of barcode signal to a reference population on barcode-specific microarrays.

**Figure 2– figure supplement 1.**
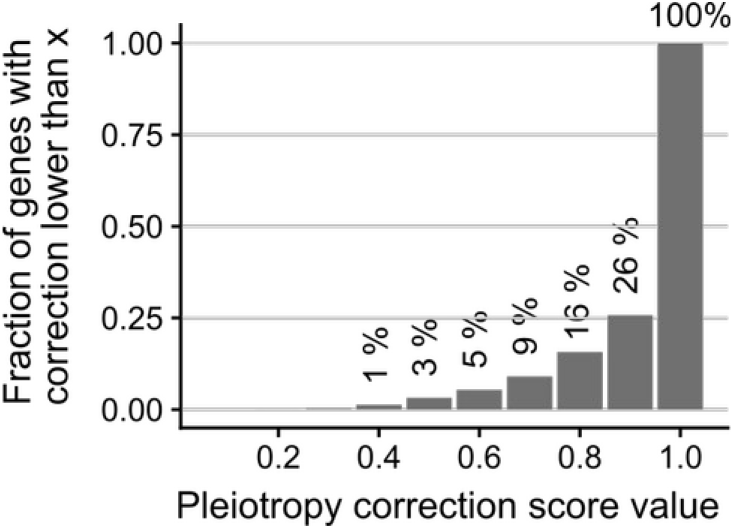
A small fraction of results were strongly affected by the pleiotropy correction. Distribution of the correction scores among the 5063 mutant results showing that, for 74% of the cases, the raw results were multiplied by a correction factor between 0.9 and 1. Only for 65 genes (1.3%) the correction factor was lower than 0.4.

**Figure 3– figure supplement 1.**
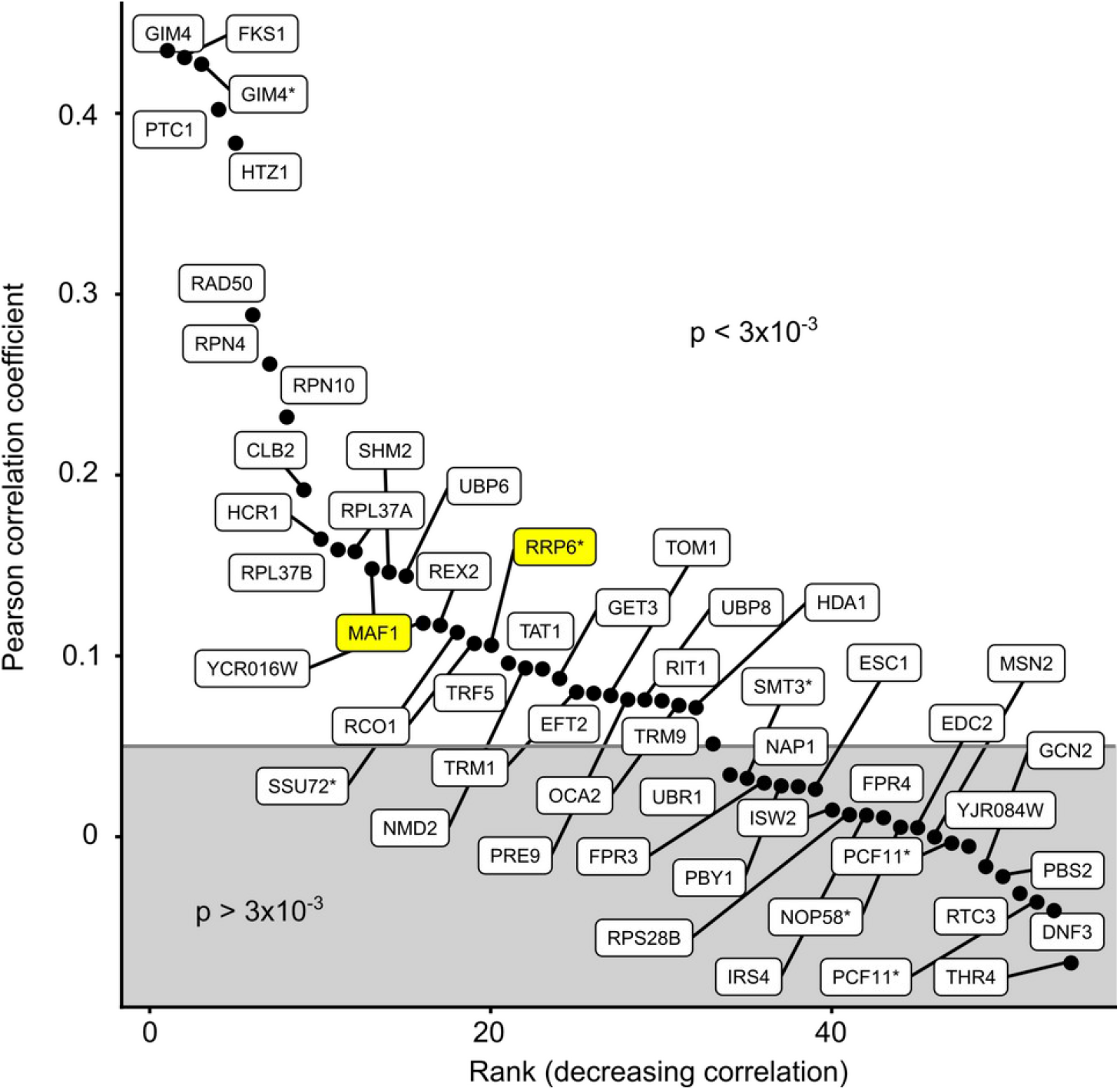
Pearson correlation coefficients for 52 GIM screens for which equivalent SGA results were available (Costanzo *et al*, 2010) were drawn in decreasing order of correlation coefficient, with a boundary situated at an arbitrary threshold for the Pearson correlation p-value of 3×10^−3^. Labels with a star indicate screens for which the mutants were DAmP or *Tet-off* variants. Screens for which correlations are shown in **Fig. 3 C** and **3 D** are highlighted in yellow.

**Figure 4– figure supplement 1.**
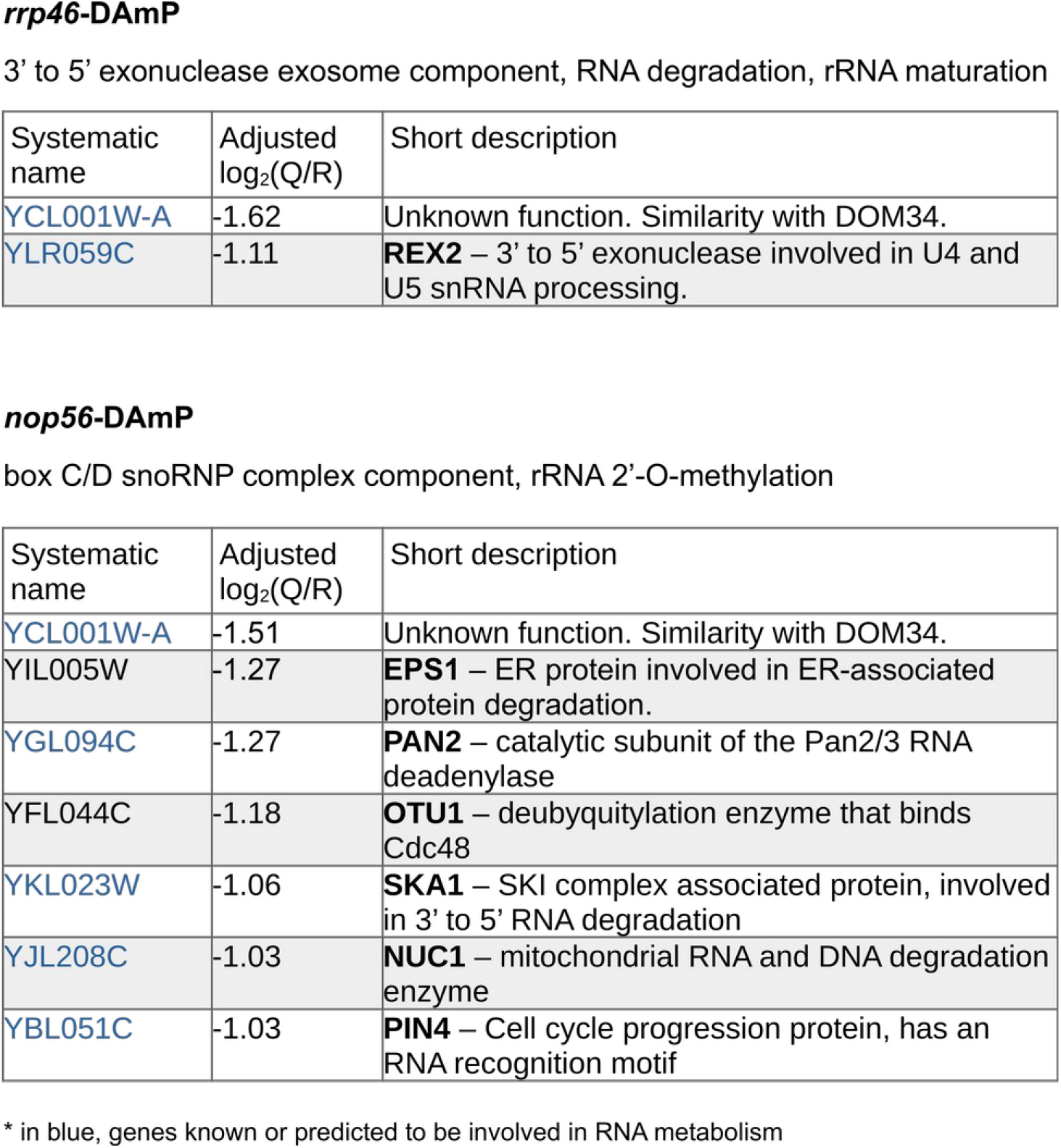
Examples of identification of novel RNA metabolism genes through GIs with DAmP mutants. The DAmP modification of RRP46, a component of the RNA exosome, and of NOP56, part of the box C/D snoRNPs, showed a synthetic growth defect with the deletion of YCL001W-A, an uncharacterized gene. The corresponding protein has similarity with a region of DOM34, a protein involved in ribosome dissociation and RNA degradation during no-go decay. SKA1, whose deletion was synthetic sick with *nop56*-DAmP was recently described to have a role in the 3’ to 5’ degradation of RNAs (Zhang *et al*, 2019).

**Figure 5– figure supplement 1.**
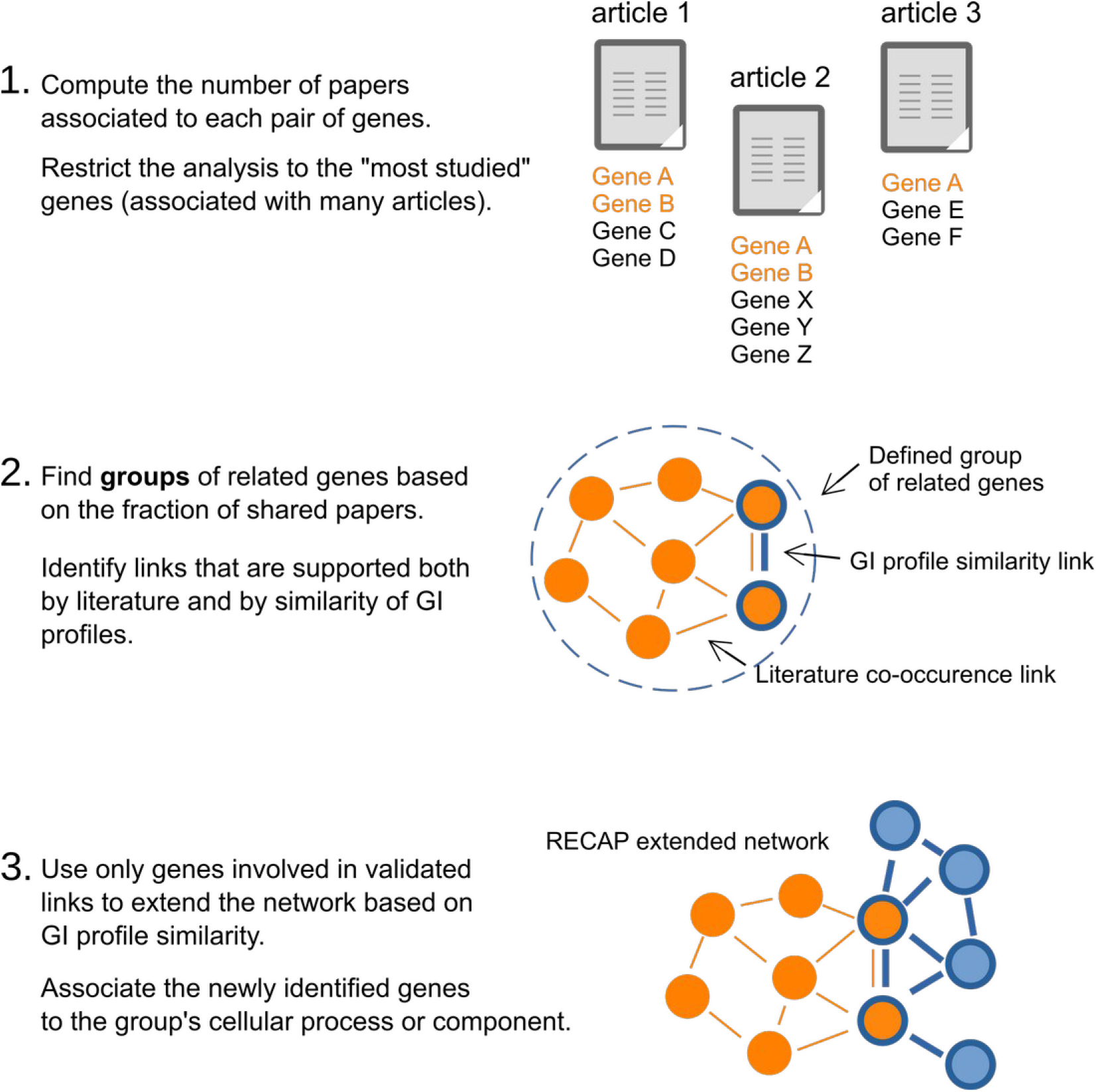
The RECAP data analysis workflow. RECAP uses an initial network of functional links between genes to validate GI profile results. Only validated gene mutants are then used to extend the network and associate new genes with known biological processes or multiprotein complexes.

**Figure 6– figure supplement 1.**
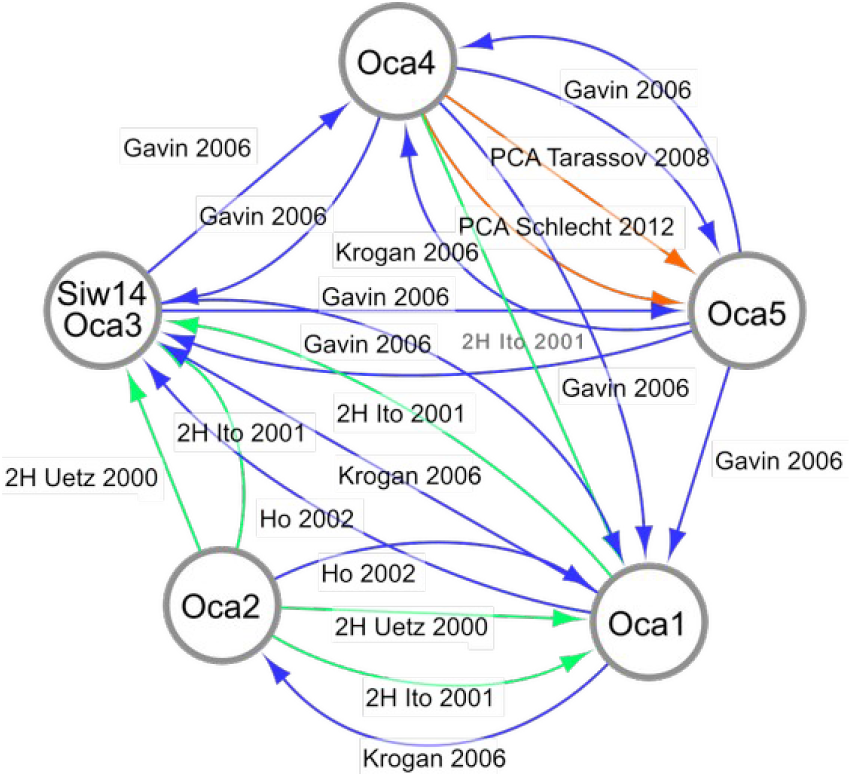
Protein-protein interactions for 5 of the 6 similar OCA proteins. The results were obtained from BioGrid (Oughtred *et al*, 2019), drawn using Cytoscape (Shannon *et al*, 2003) with the source of the interactions mentioned on the arrows. “2H” indicates two-hybrid screens (Uetz *et al*, 2000; Ito *et al*, 2001), “PCA” is for protein complementation assay (Tarassov *et al*, 2008), while the other studies used affinity purification and mass-spectrometry identification of partners (Ho *et al*, 2002; Krogan *et al*, 2006; Gavin *et al*, 2006).

## Notes

http://hub05.hosting.pasteur.fr/GIM_interactions/

### Competing Interest Statement

The authors have declared no competing interest.

