## Supplementary material for "Investigation of RNA metabolism through large-scale genetic interaction profiling in yeast": Exclusion peaks correction for all screens

Supplementary figure set (exclusion peak correction)

**Exclusion peak correction for GIM screens**. For each microarray hybridation file, the horizontal axis represents the relative genomic position of the affected gene, while the vertical axis represents, in $\log_2$ transformed values, the measured growth defect. The title indicates the identifier of the screen replicate. For example « YDR283C GCN2 DEL 2 » corresponds to the screen done with the deletion (« DEL ») of the GCN2 gene, systematic name « YDR283C », for replicate number 2.

In red, original values, in black, values after correction using an estimation of recombination frequencies.

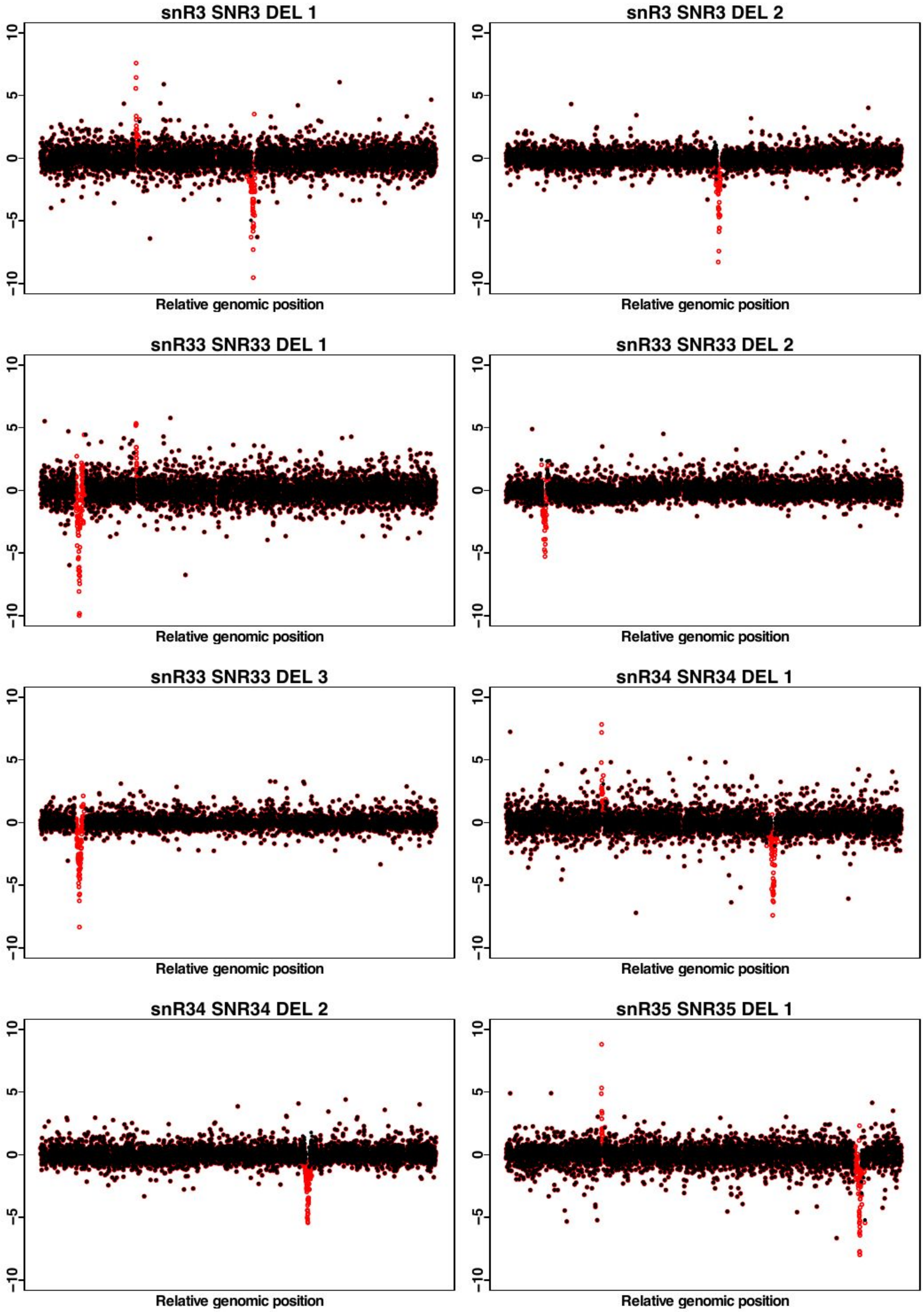
snR3 SNR3 DEL 1
snR3 SNR3 DEL 2
snR33 SNR33 DEL 1
snR33 SNR33 DEL 2
snR33 SNR33 DEL 3
snR34 SNR34 DEL 1
snR34 SNR34 DEL 2
snR35 SNR35 DEL 1
10
5
0
-5
-10
Relative genomic position

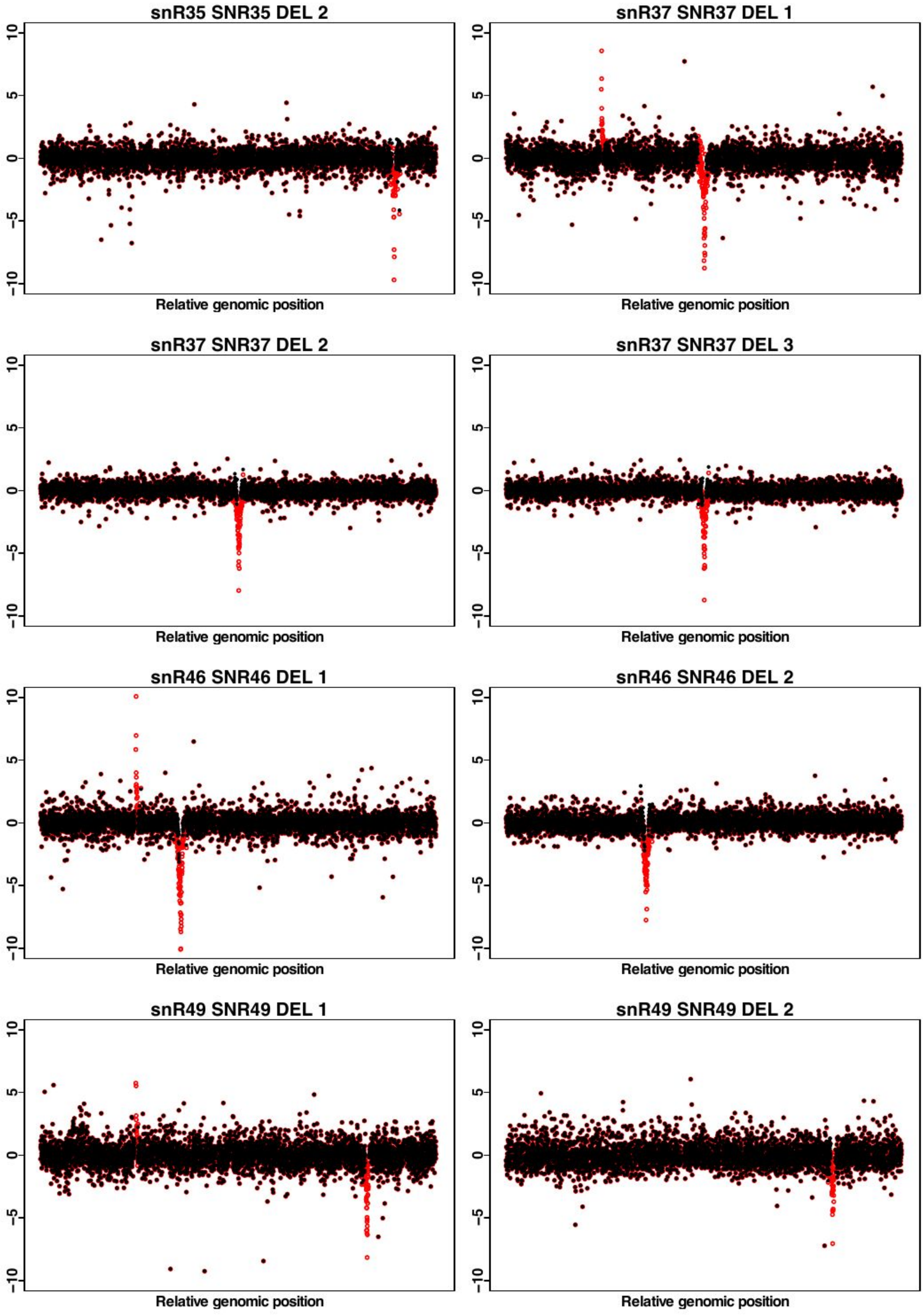
snR35 SNR35 DEL 2
snR37 SNR37 DEL 1
snR37 SNR37 DEL 2
snR37 SNR37 DEL 3
snR46 SNR46 DEL 1
snR46 SNR46 DEL 2
snR49 SNR49 DEL 1
snR49 SNR49 DEL 2
10
5
0
−5
−10
Relative genomic position

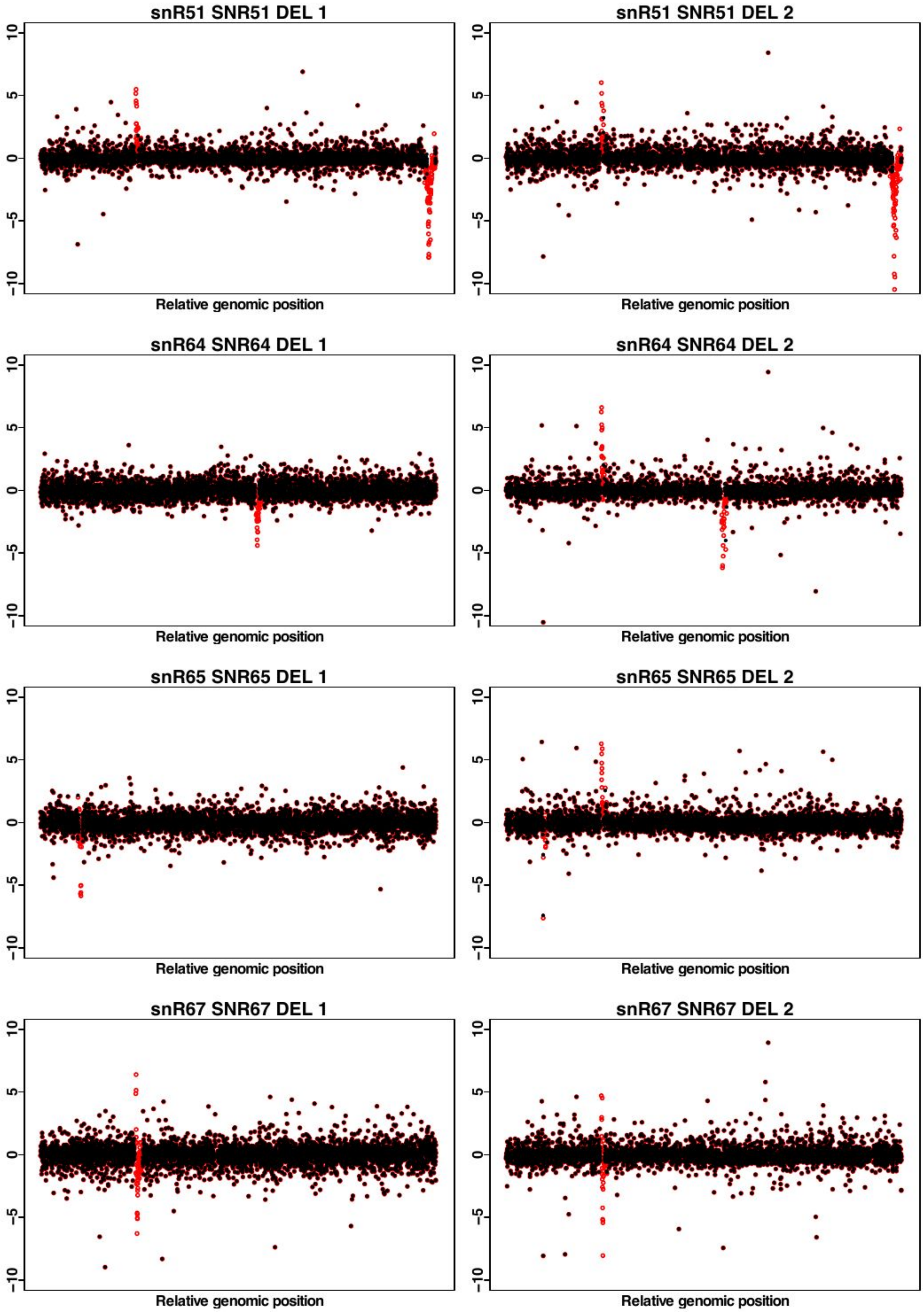
snR51 SNR51 DEL 1
snR51 SNR51 DEL 2
snR64 SNR64 DEL 1
snR64 SNR64 DEL 2
snR65 SNR65 DEL 1
snR65 SNR65 DEL 2
snR67 SNR67 DEL 1
snR67 SNR67 DEL 2
10
5
0
-5
-10
Relative genomic position

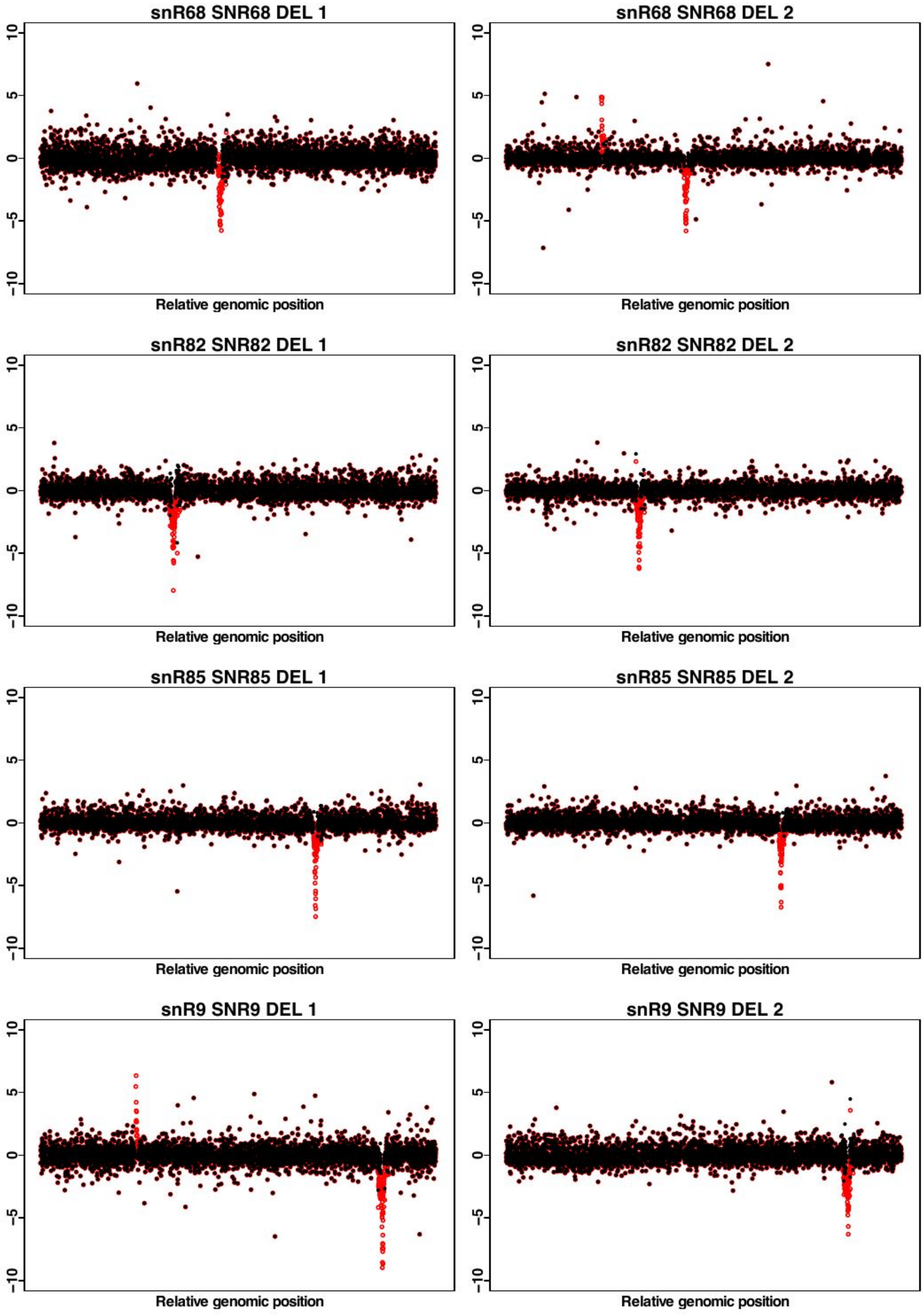
snR68 SNR68 DEL 1
snR68 SNR68 DEL 2
snR82 SNR82 DEL 1
snR82 SNR82 DEL 2
snR85 SNR85 DEL 1
snR85 SNR85 DEL 2
snR9 SNR9 DEL 1
snR9 SNR9 DEL 2
10
5
0
−5
−10
Relative genomic position

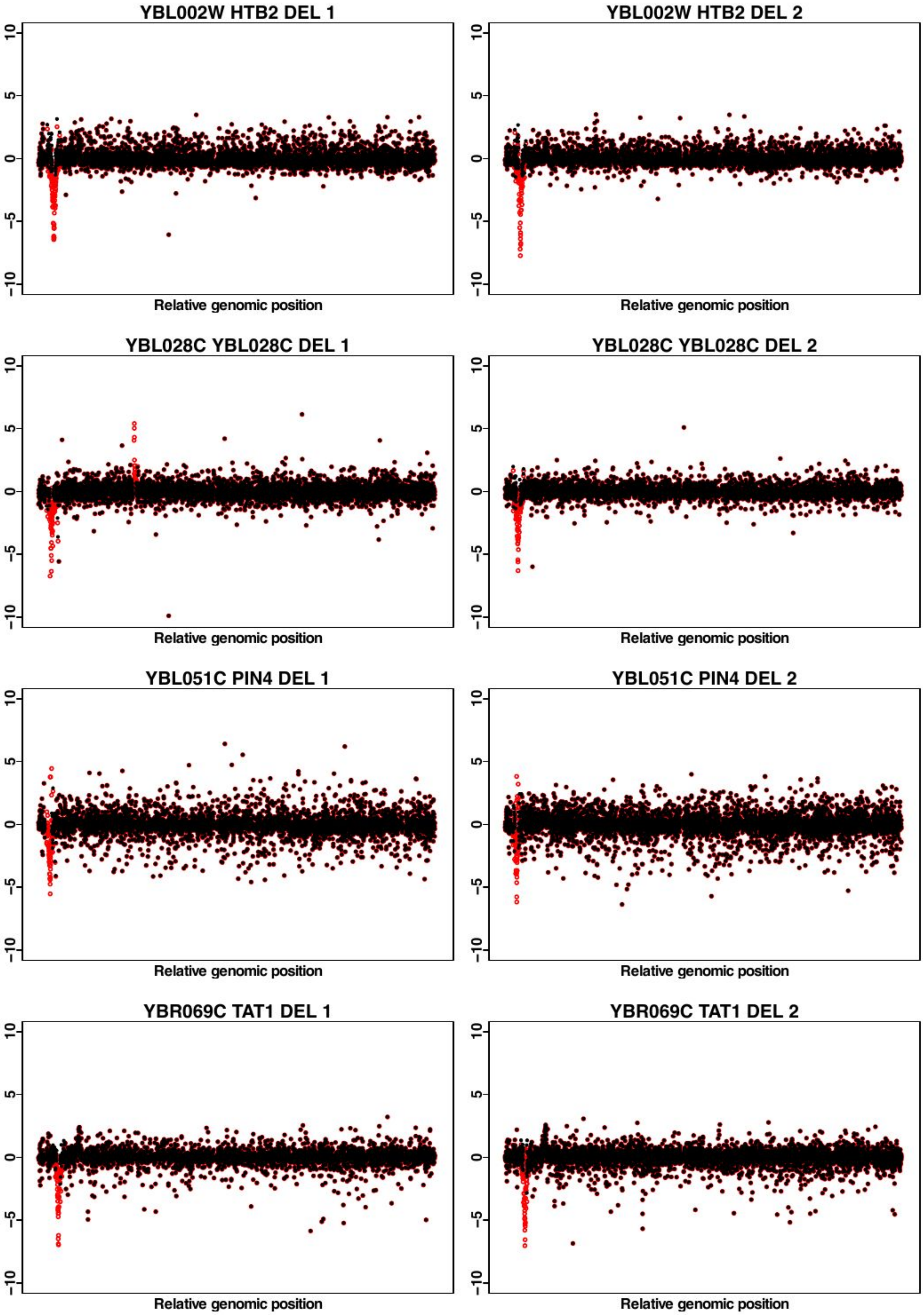

YBL002W HTB2 DEL 1
YBL002W HTB2 DEL 2
YBL028C YBL028C DEL 1
YBL028C YBL028C DEL 2
YBL051C PIN4 DEL 1
YBL051C PIN4 DEL 2
YBR069C TAT1 DEL 1
YBR069C TAT1 DEL 2
10
5
0
−5
−10
Relative genomic position

YBR094W PBY1 DEL 1

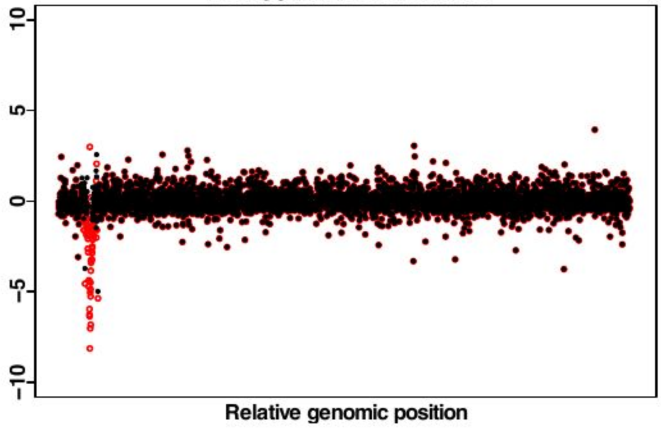


YBR094W PBY1 DEL 2

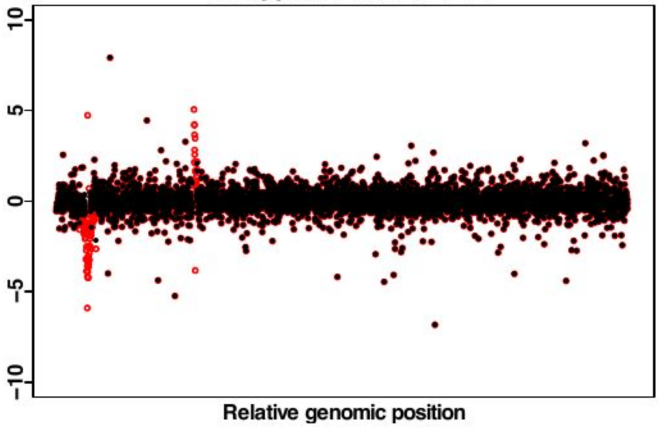


YBR255W MTC4 DEL 1

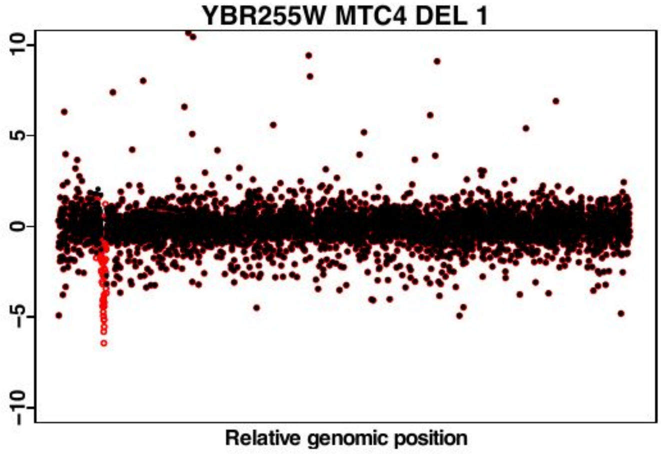


YBR255W MTC4 DEL 2

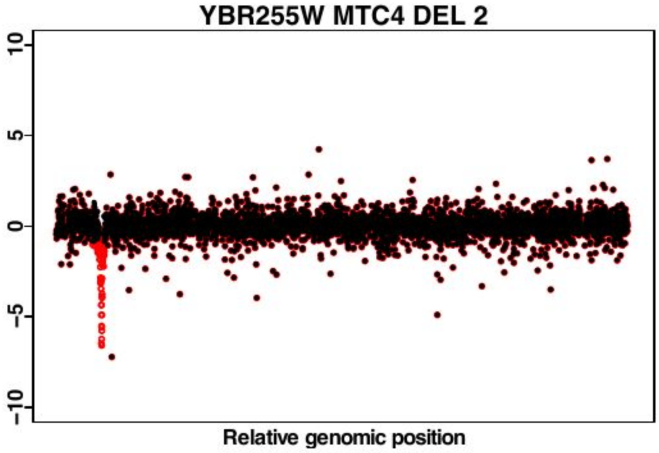


YBR255W MTC4 DEL 3

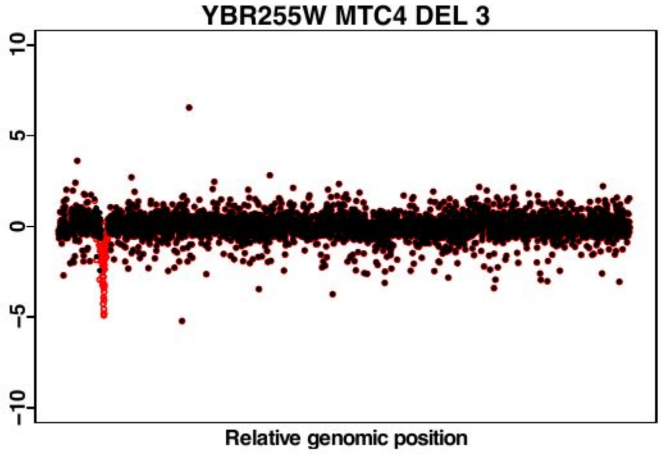


YBR281C DUG2 DEL 1

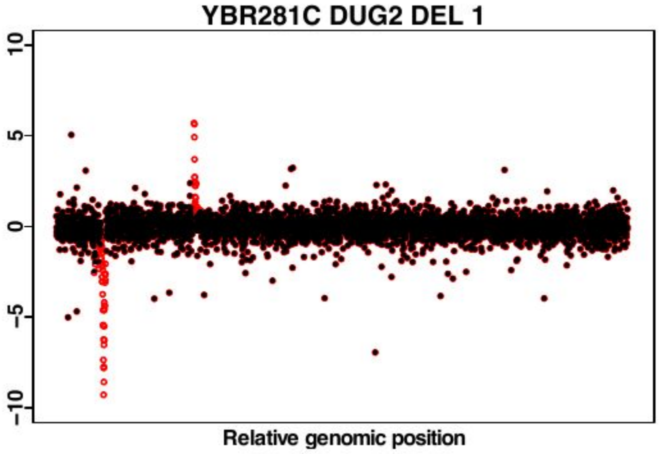


YBR281C DUG2 DEL 2

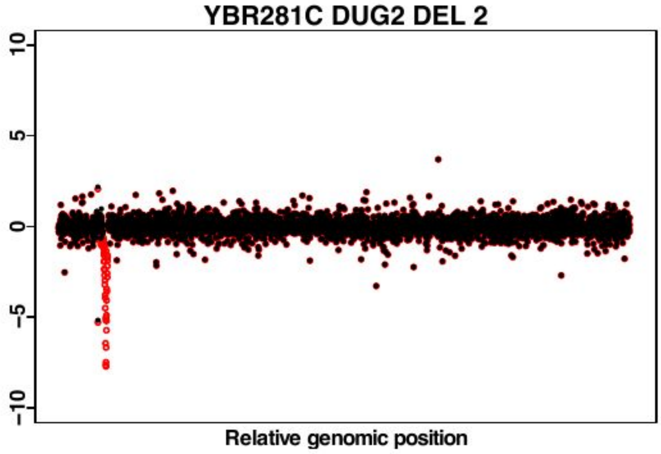


YCL001W.A YCL001W.A DEL 1

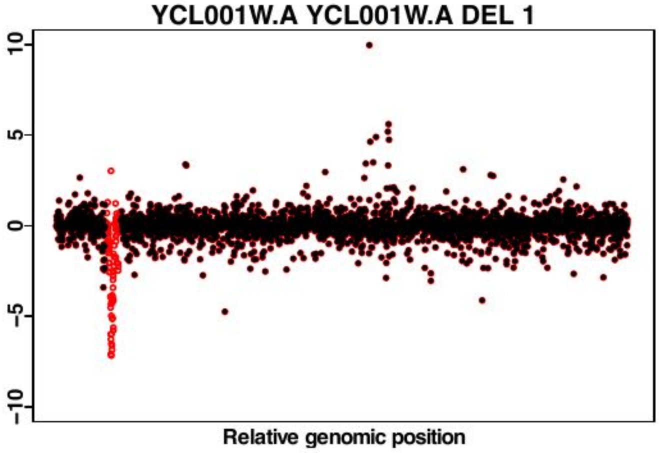

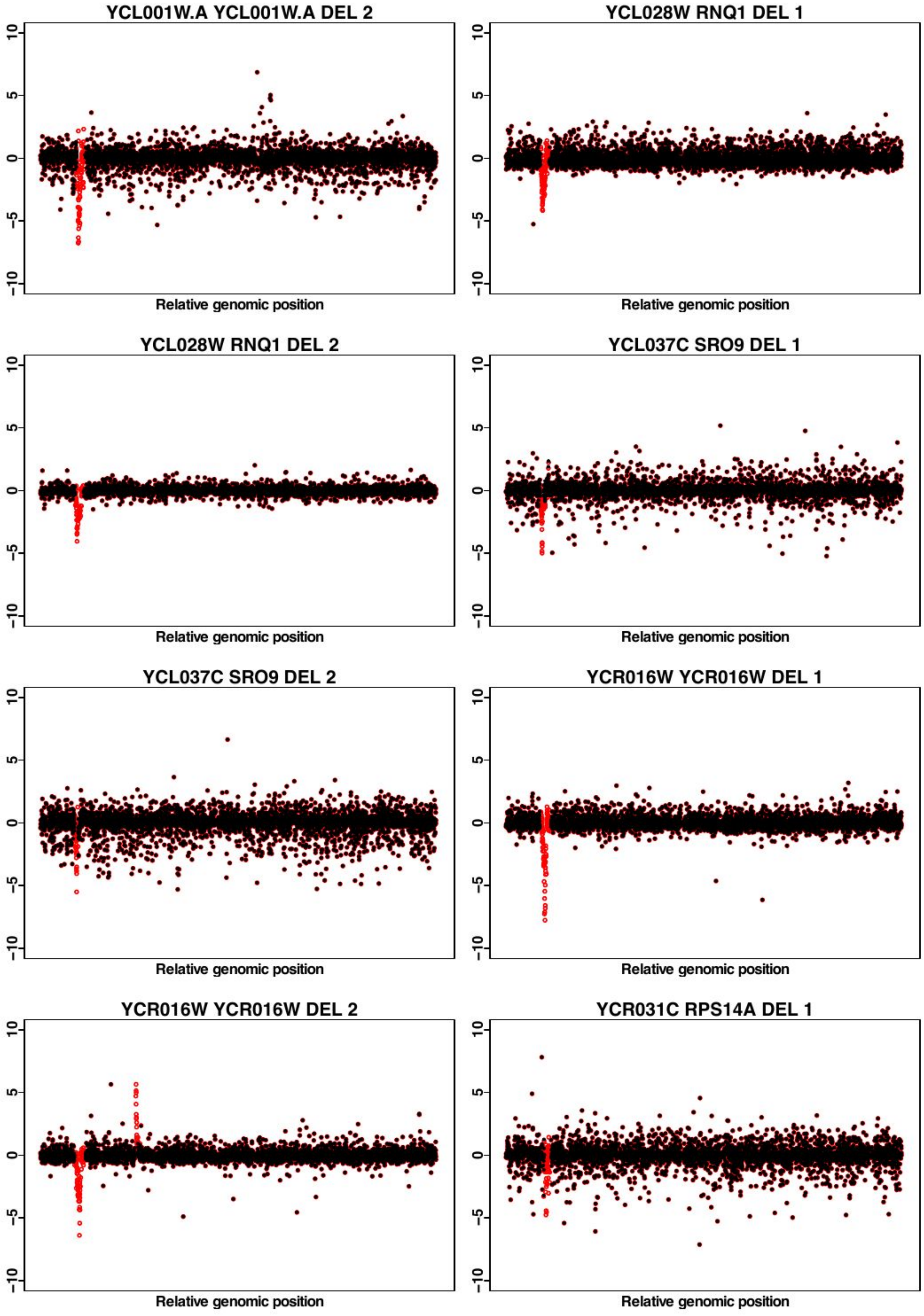
YCL001W.A YCL001W.A DEL 2
YCL028W RNQ1 DEL 1
YCL028W RNQ1 DEL 2
YCL037C SRO9 DEL 1
YCL037C SRO9 DEL 2
YCR016W YCR016W DEL 1
YCR016W YCR016W DEL 2
YCR031C RPS14A DEL 1
10
5
0
-5
-10
Relative genomic position

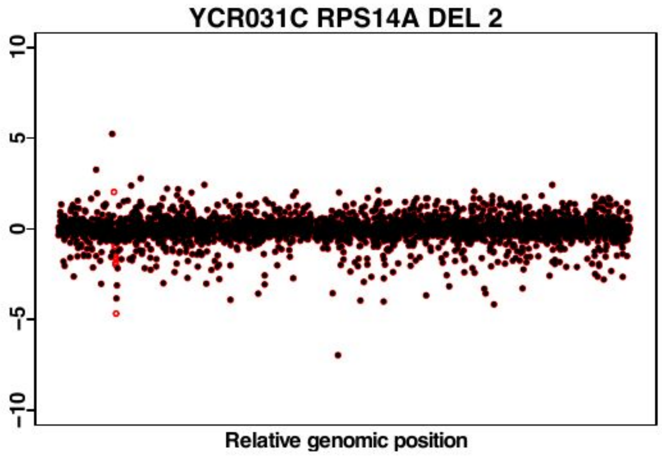
YCR031C RPS14A DEL 2
10
5
0
-5
-10
Relative genomic position

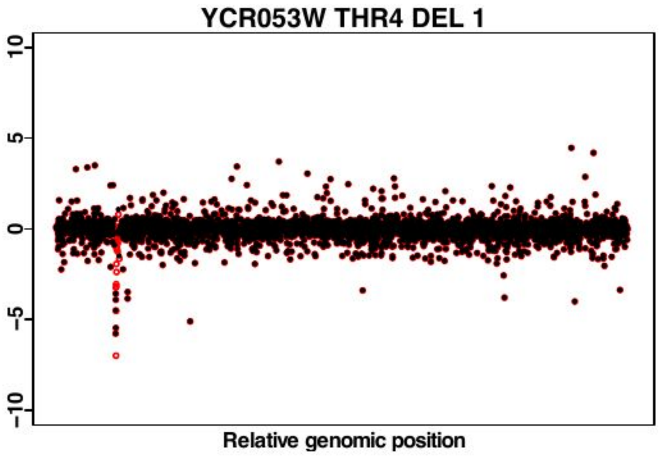
YCR053W THR4 DEL 1
10
5
0
-5
-10
Relative genomic position

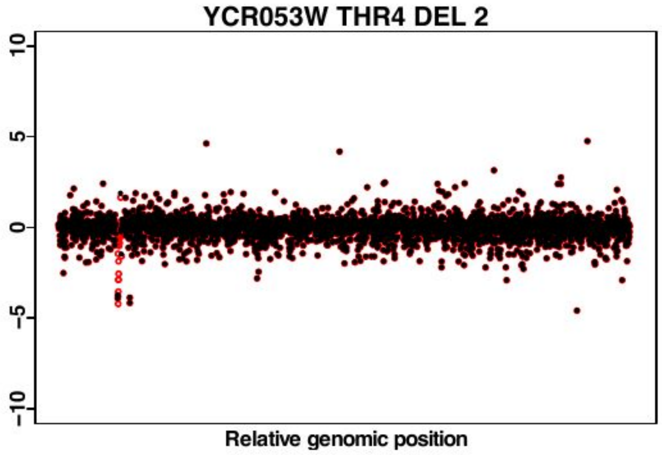
YCR053W THR4 DEL 2
10
5
0
-5
-10
Relative genomic position

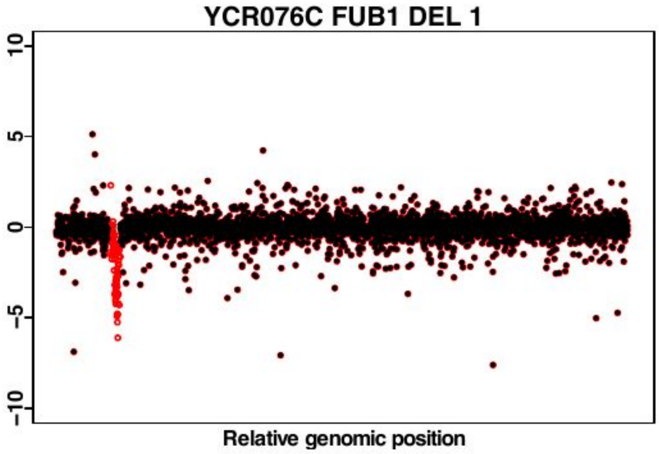
YCR076C FUB1 DEL 1
10
5
0
-5
-10
Relative genomic position

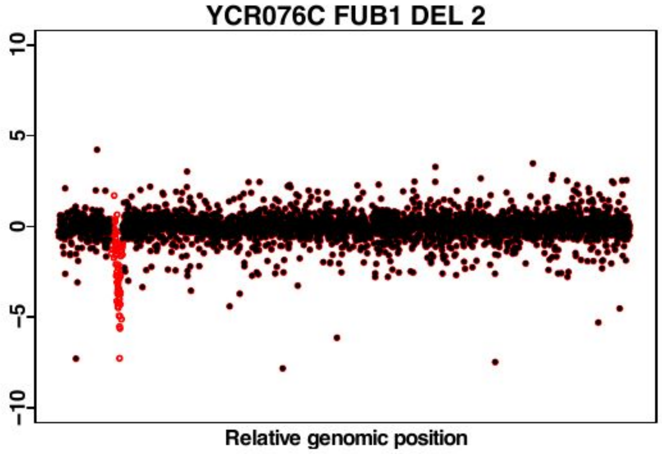
YCR076C FUB1 DEL 2
10
5
0
-5
-10
Relative genomic position

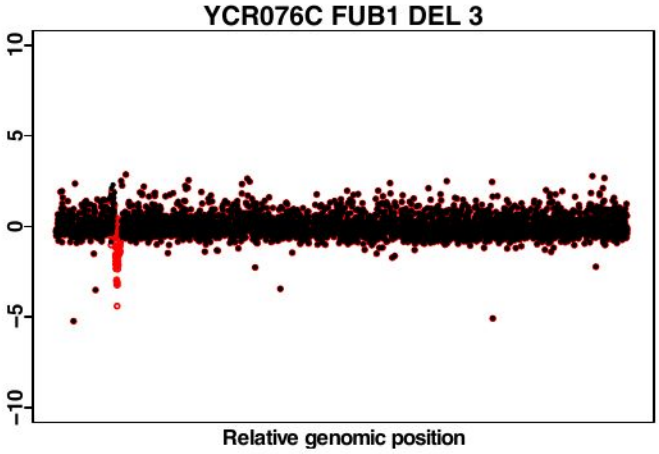
YCR076C FUB1 DEL 3
10
5
0
-5
-10
Relative genomic position

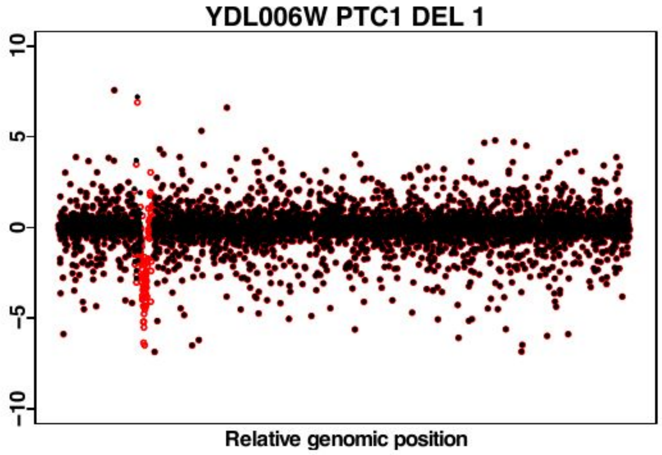
YDL006W PTC1 DEL 1
10
5
0
-5
-10
Relative genomic position

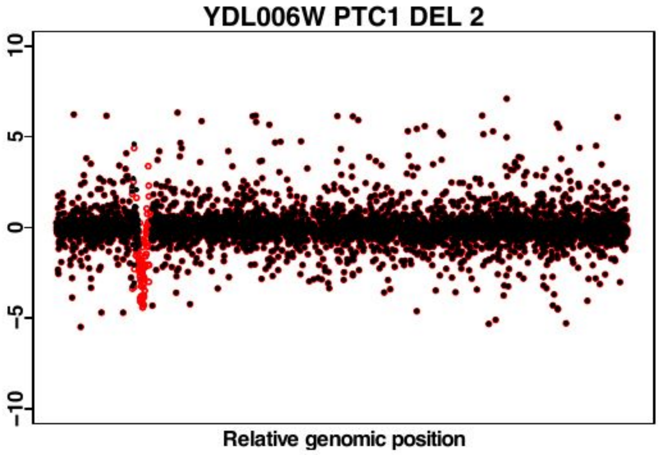
YDL006W PTC1 DEL 2
10
5
0
-5
-10
Relative genomic position

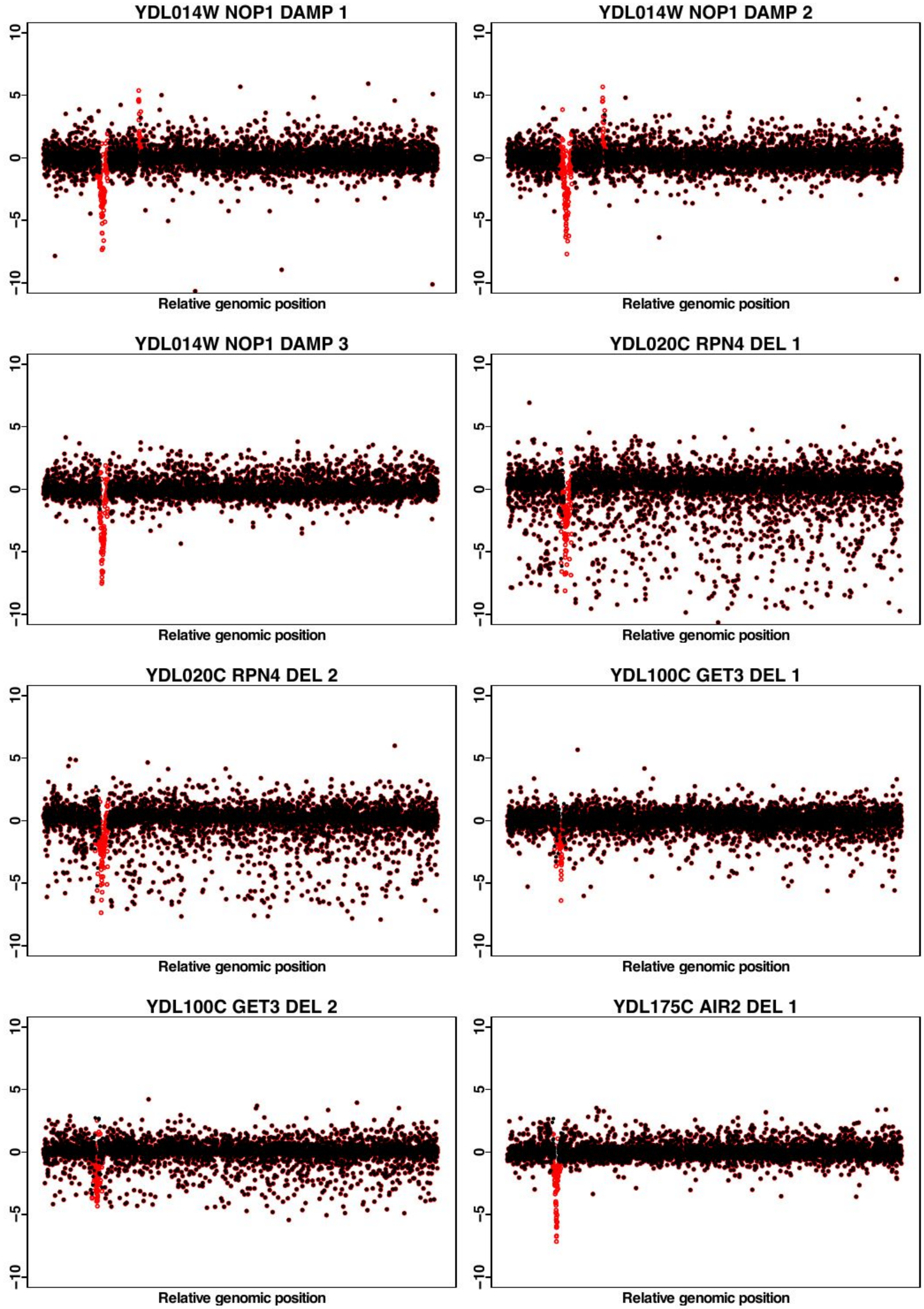
YDL014W NOP1 DAMP 1
YDL014W NOP1 DAMP 2
YDL014W NOP1 DAMP 3
YDL020C RPN4 DEL 1
YDL020C RPN4 DEL 2
YDL100C GET3 DEL 1
YDL100C GET3 DEL 2
YDL175C AIR2 DEL 1
Relative genomic position
-10
-5
0
5
10

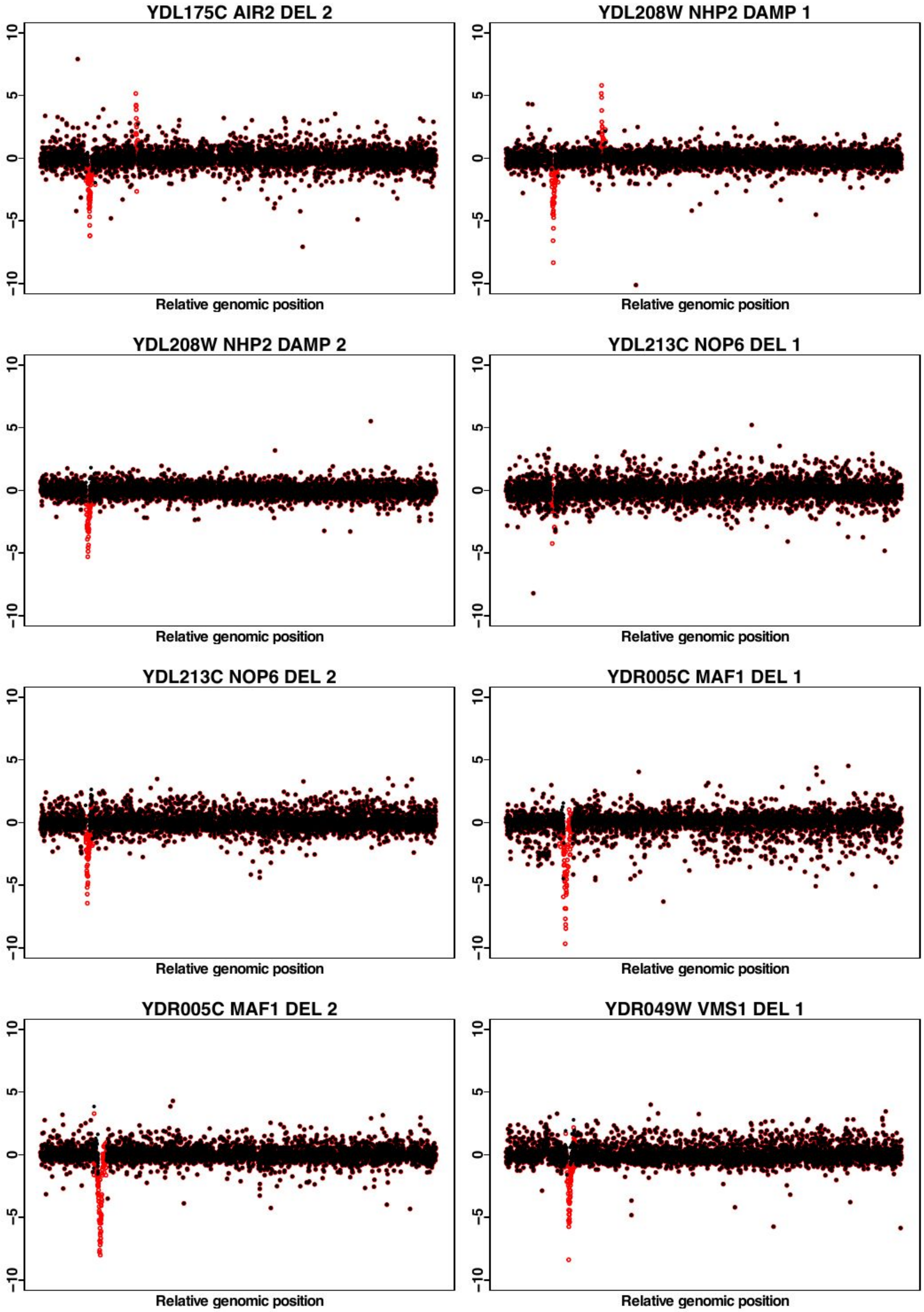
YDL175C AIR2 DEL 2
YDL208W NHP2 DAMP 1
YDL208W NHP2 DAMP 2
YDL213C NOP6 DEL 1
YDL213C NOP6 DEL 2
YDR005C MAF1 DEL 1
YDR005C MAF1 DEL 2
YDR049W VMS1 DEL 1
Relative genomic position
10
5
0
-5
-10

YDR049W VMS1 DEL 2

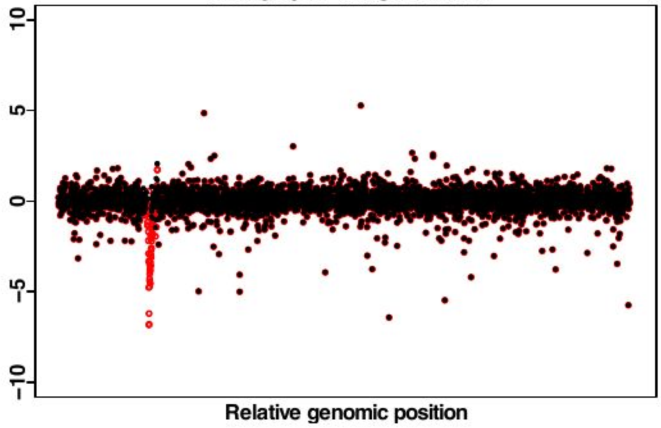


YDR049W VMS1 DEL 3

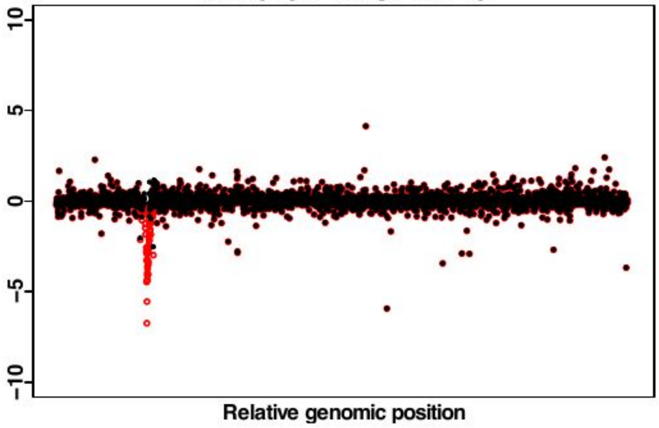


YDR049W VMS1 DEL 4

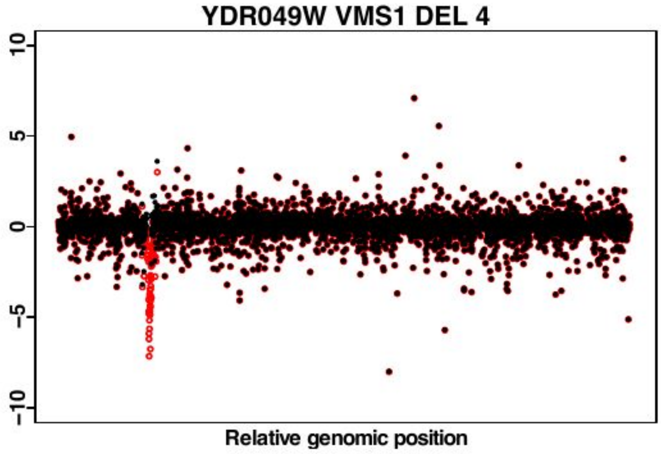


YDR059C UBC5 DEL 1

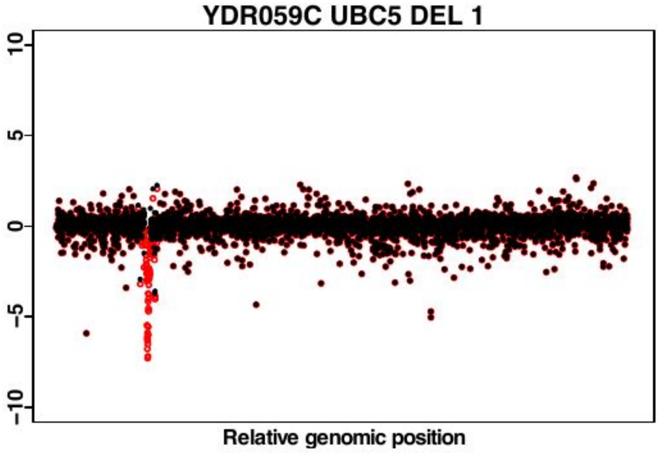


YDR059C UBC5 DEL 2

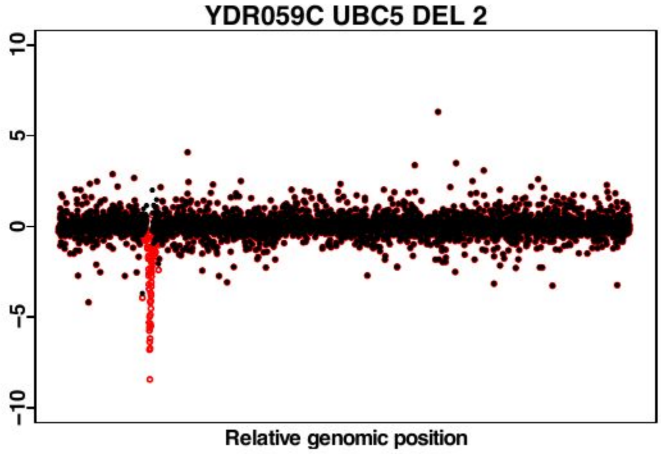


YDR067C OCA6 DEL 1

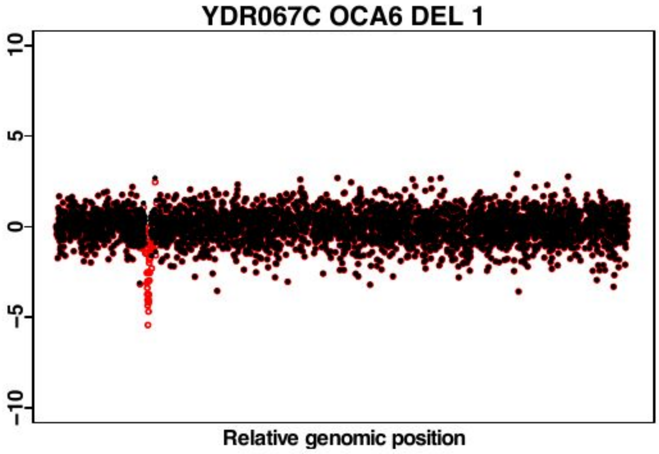


YDR067C OCA6 DEL 2

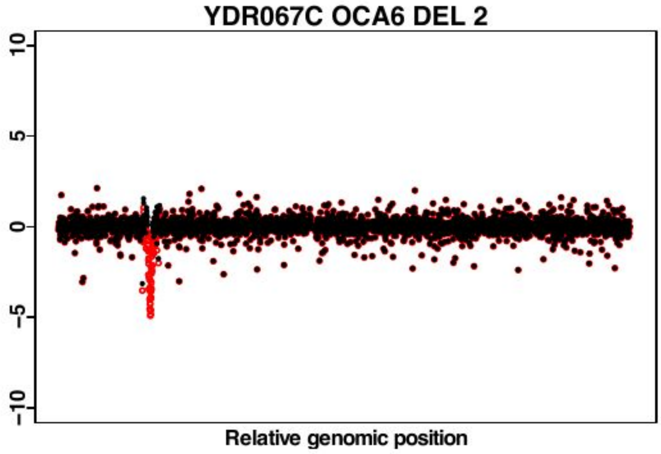


YDR120C TRM1 DEL 1

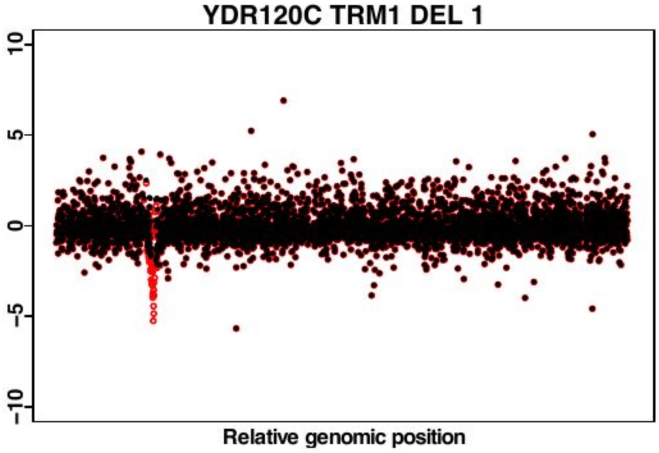

YDR120C TRM1 DEL 2

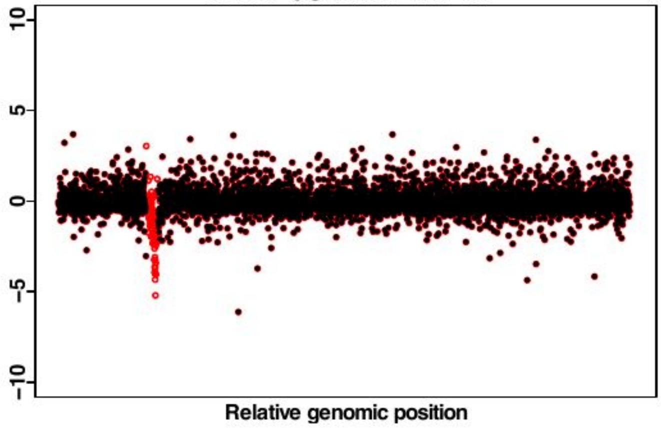


YDR140W MTQ2 DAMP 1

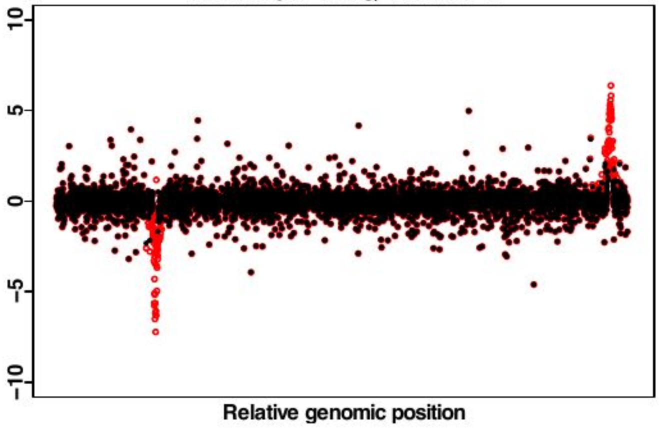


YDR140W MTQ2 DAMP 2

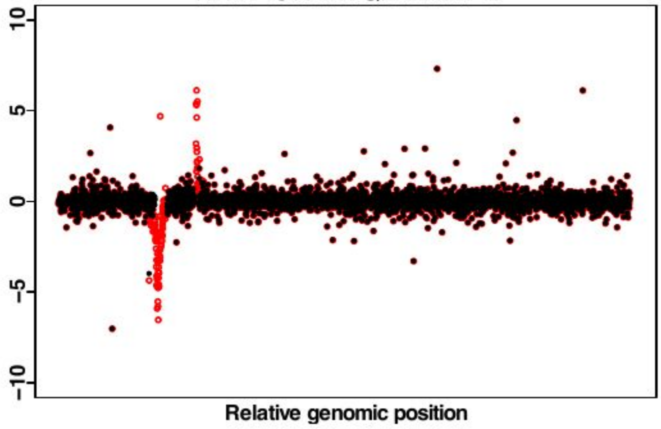


YDR143C SAN1 DEL 1

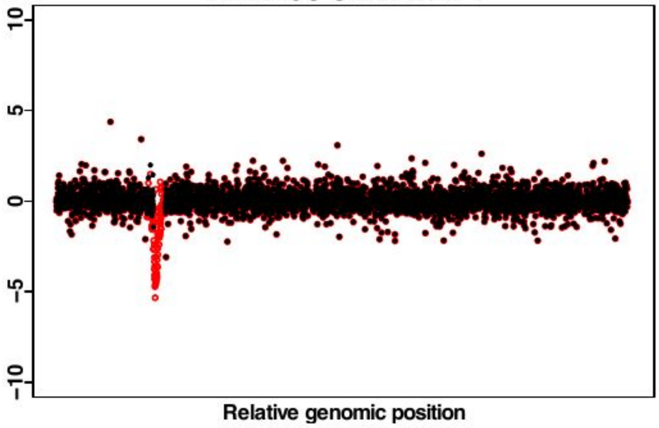


YDR143C SAN1 DEL 2

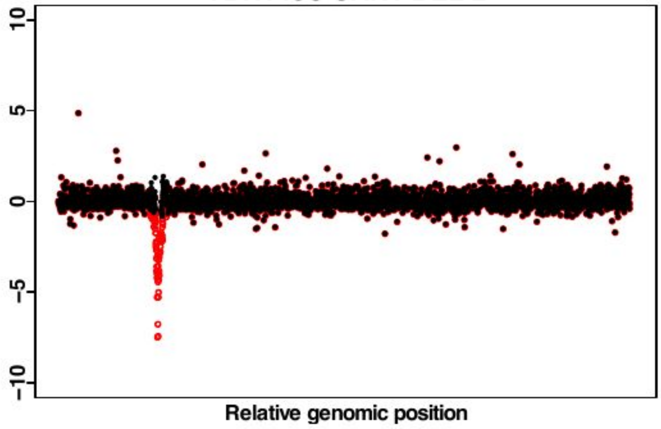


YDR228C PCF11 DAMP 1

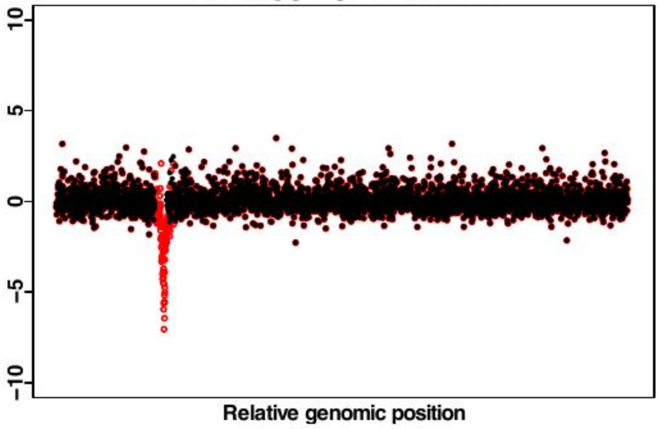


YDR228C PCF11 DAMP 2

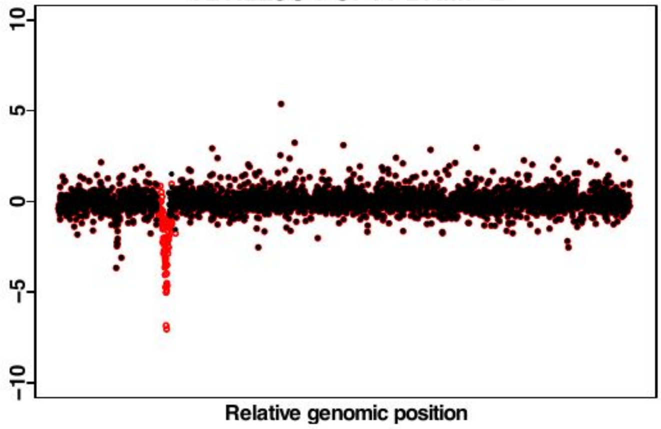


YDR283C GCN2 DEL 1

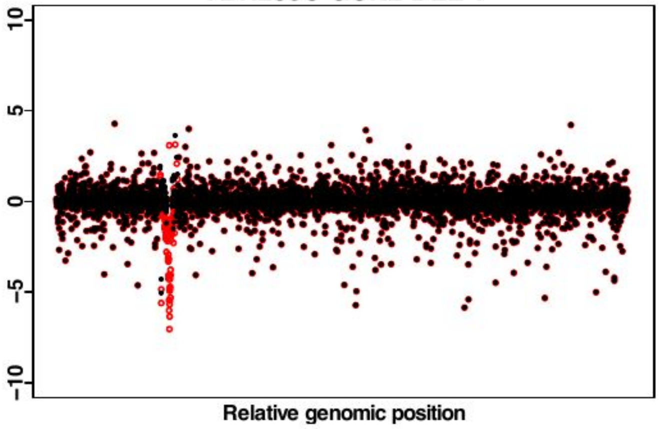

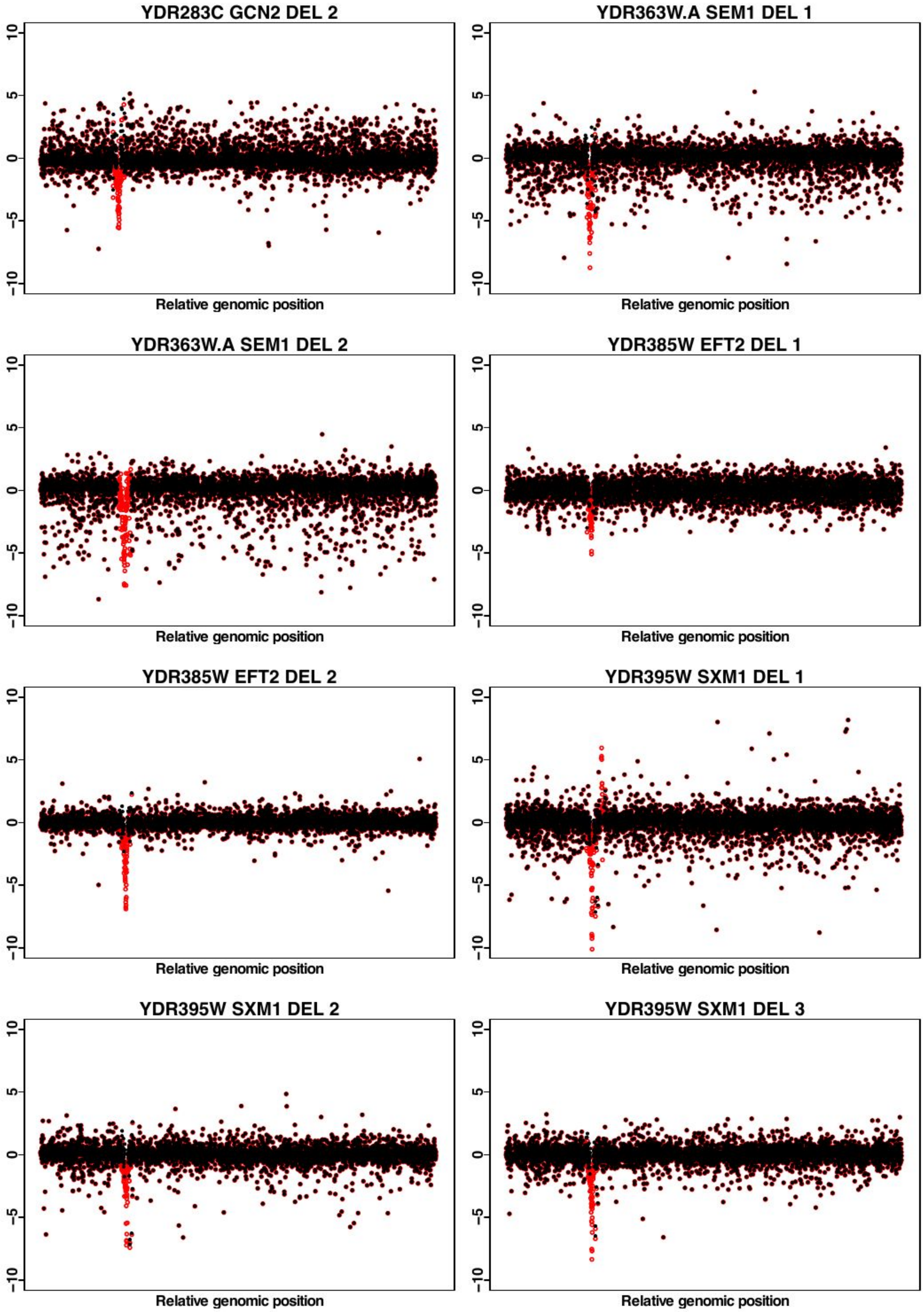

YDR283C GCN2 DEL 2
YDR363W.A SEM1 DEL 1
YDR363W.A SEM1 DEL 2
YDR385W EFT2 DEL 1
YDR385W EFT2 DEL 2
YDR395W SXM1 DEL 1
YDR395W SXM1 DEL 2
YDR395W SXM1 DEL 3
10
5
0
−5
−10
Relative genomic position

YDR395W SXM1 DEL 4

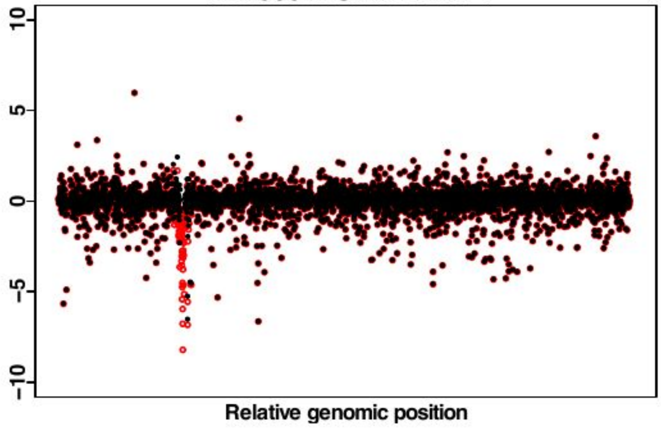


YDR418W RPL12B DEL 1

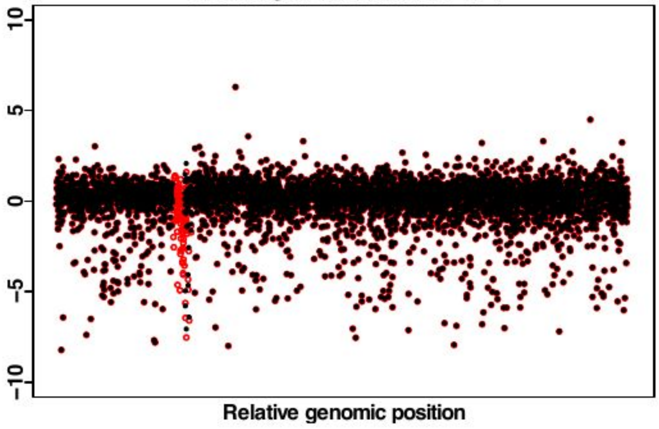


YDR418W RPL12B DEL 2

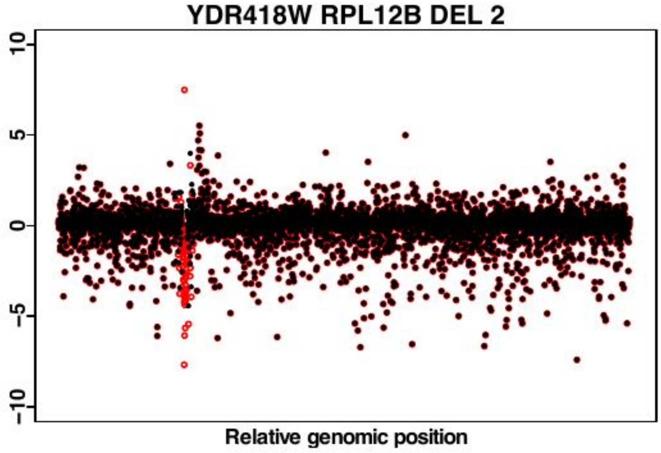


YDR457W TOM1 DEL 1

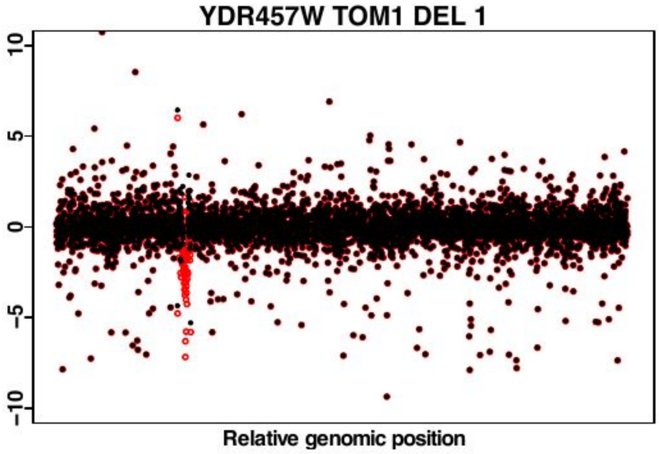


YDR457W TOM1 DEL 2

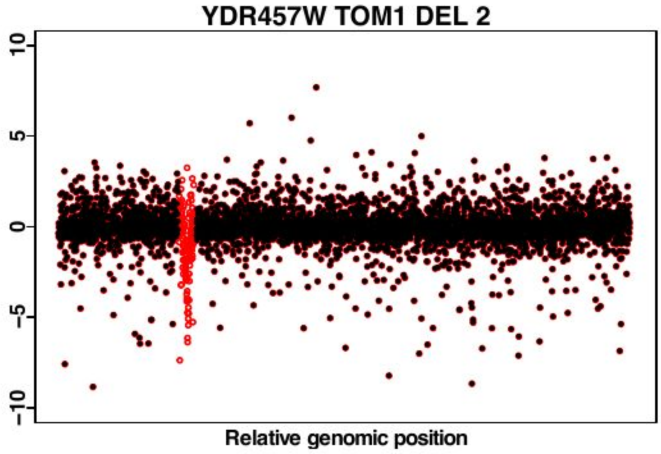


YDR500C RPL37B DEL 1

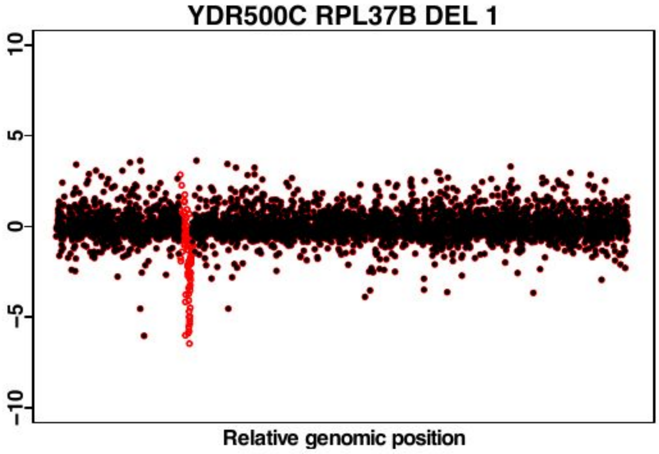


YDR500C RPL37B DEL 2

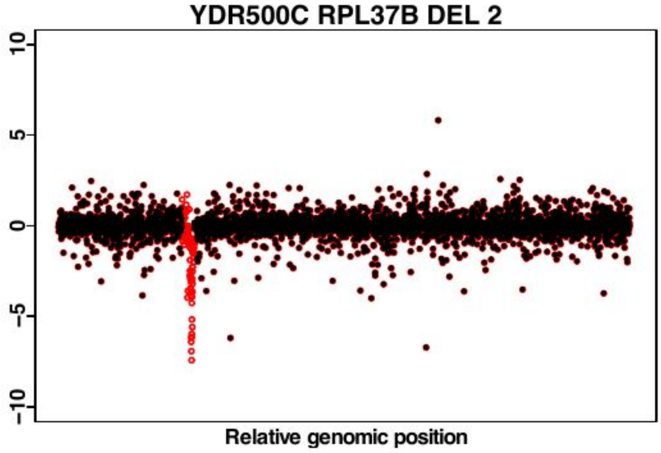


YDR510W SMT3 DAMP 1

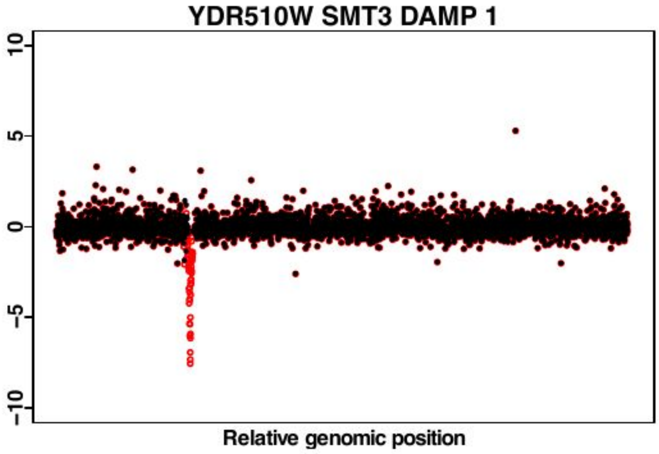

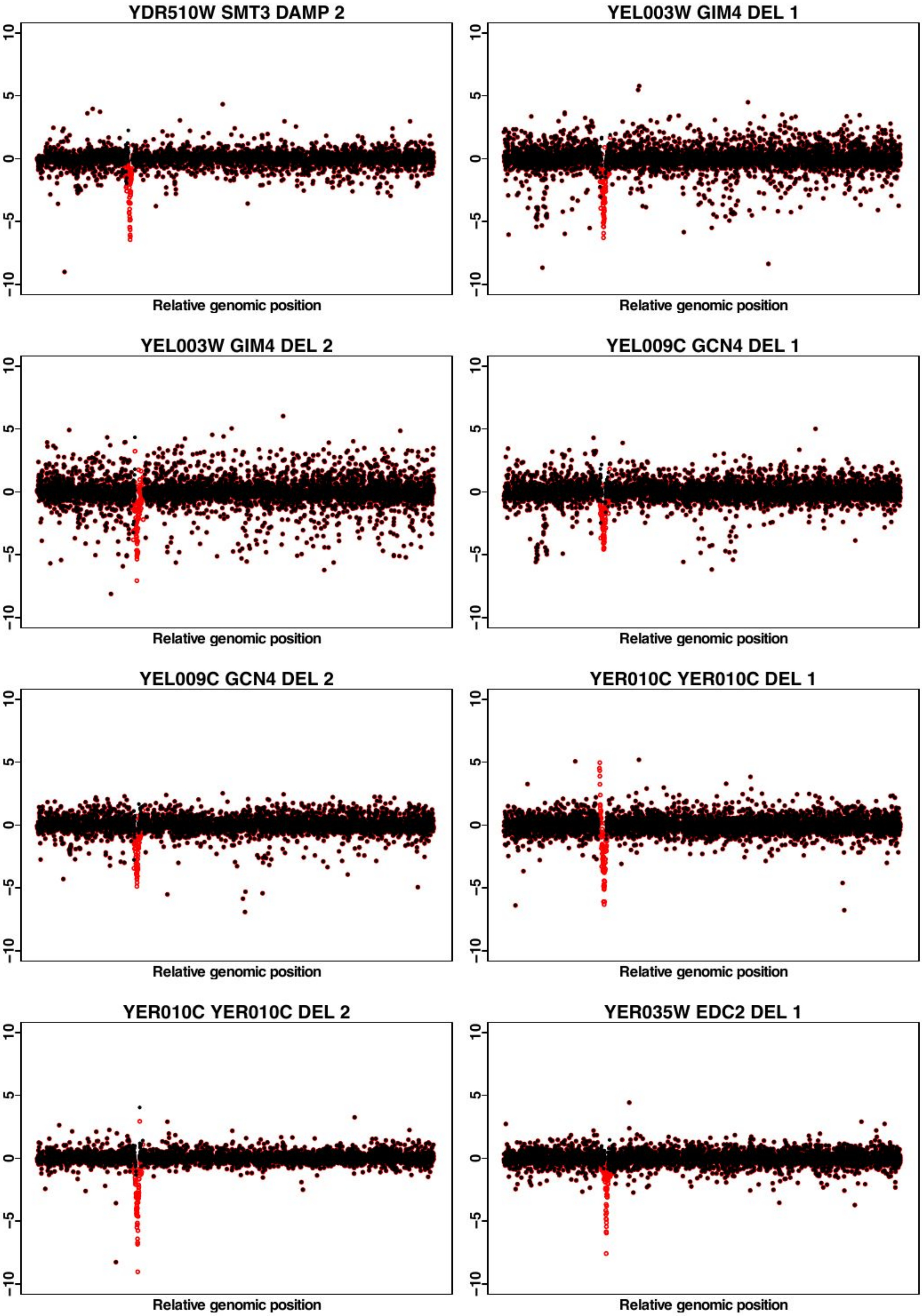
YDR510W SMT3 DAMP 2
YEL003W GIM4 DEL 1
YEL003W GIM4 DEL 2
YEL009C GCN4 DEL 1
YEL009C GCN4 DEL 2
YER010C YER010C DEL 1
YER010C YER010C DEL 2
YER035W EDC2 DEL 1
10
5
0
−5
−10
Relative genomic position

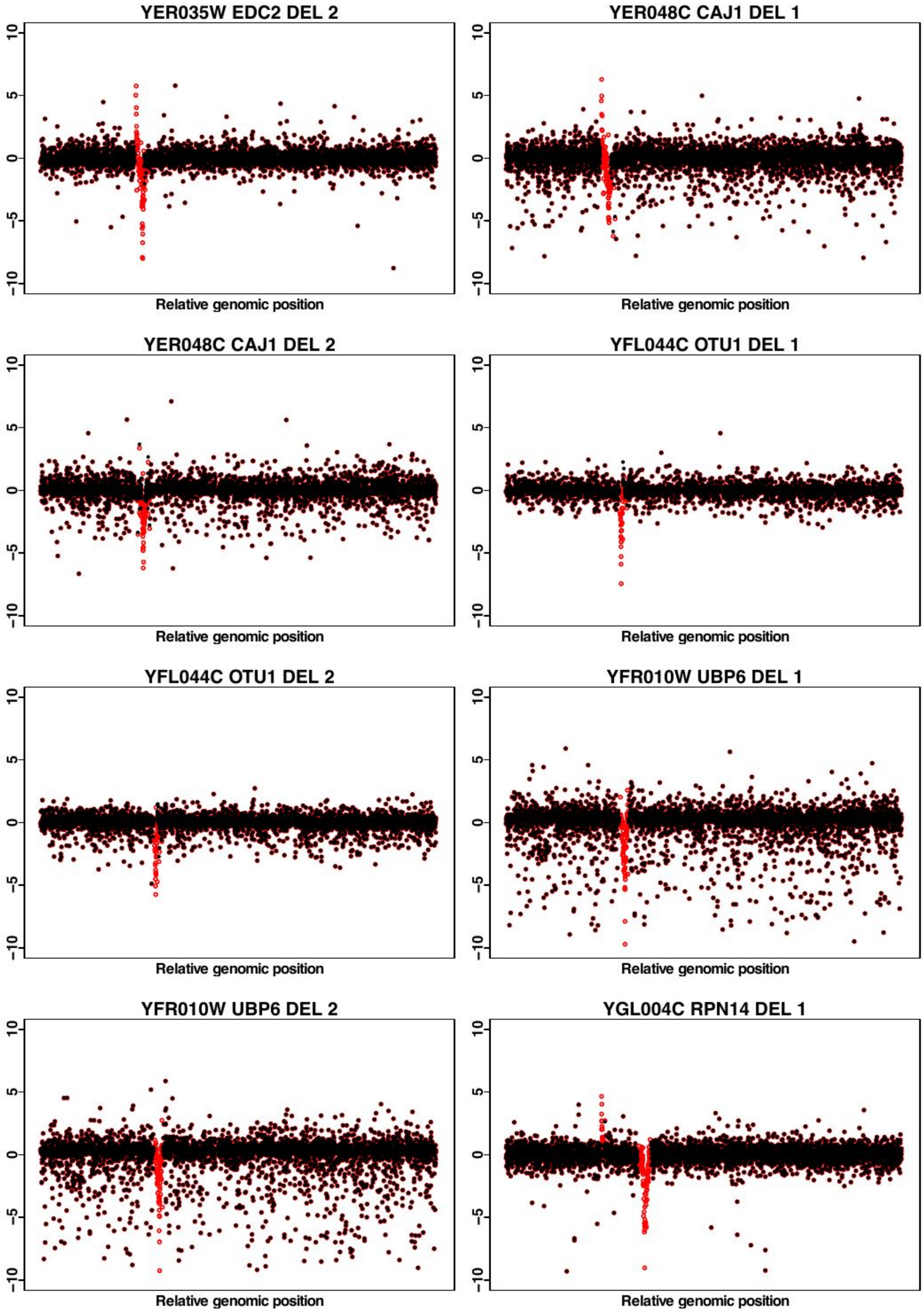
YER035W EDC2 DEL 2
YER048C CAJ1 DEL 1
YER048C CAJ1 DEL 2
YFL044C OTU1 DEL 1
YFL044C OTU1 DEL 2
YFR010W UBP6 DEL 1
YFR010W UBP6 DEL 2
YGL004C RPN14 DEL 1
10
5
0
-5
-10
Relative genomic position

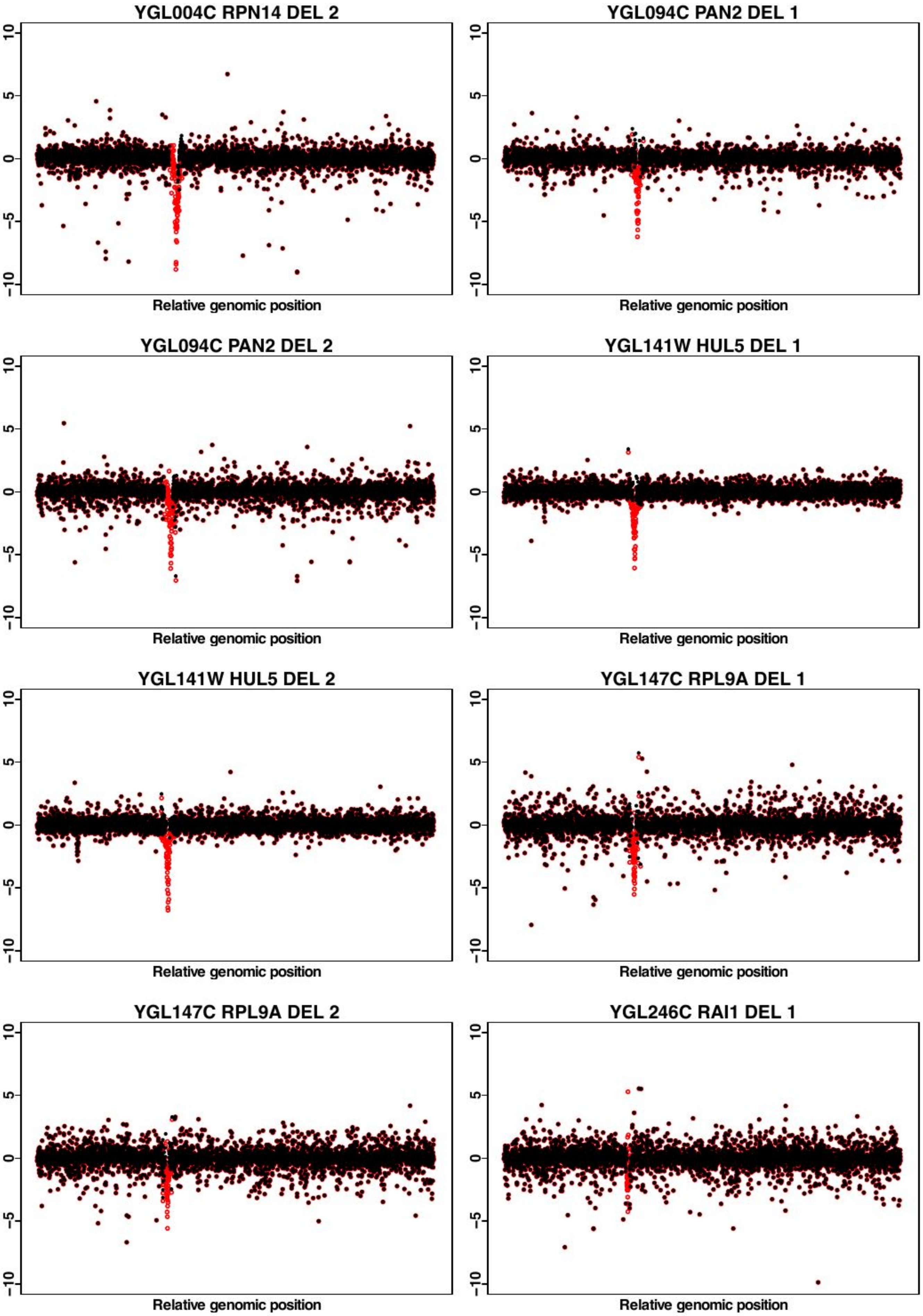
YGL004C RPN14 DEL 2
YGL094C PAN2 DEL 1
YGL094C PAN2 DEL 2
YGL141W HUL5 DEL 1
YGL141W HUL5 DEL 2
YGL147C RPL9A DEL 1
YGL147C RPL9A DEL 2
YGL246C RAI1 DEL 1
10
5
0
−5
−10
Relative genomic position

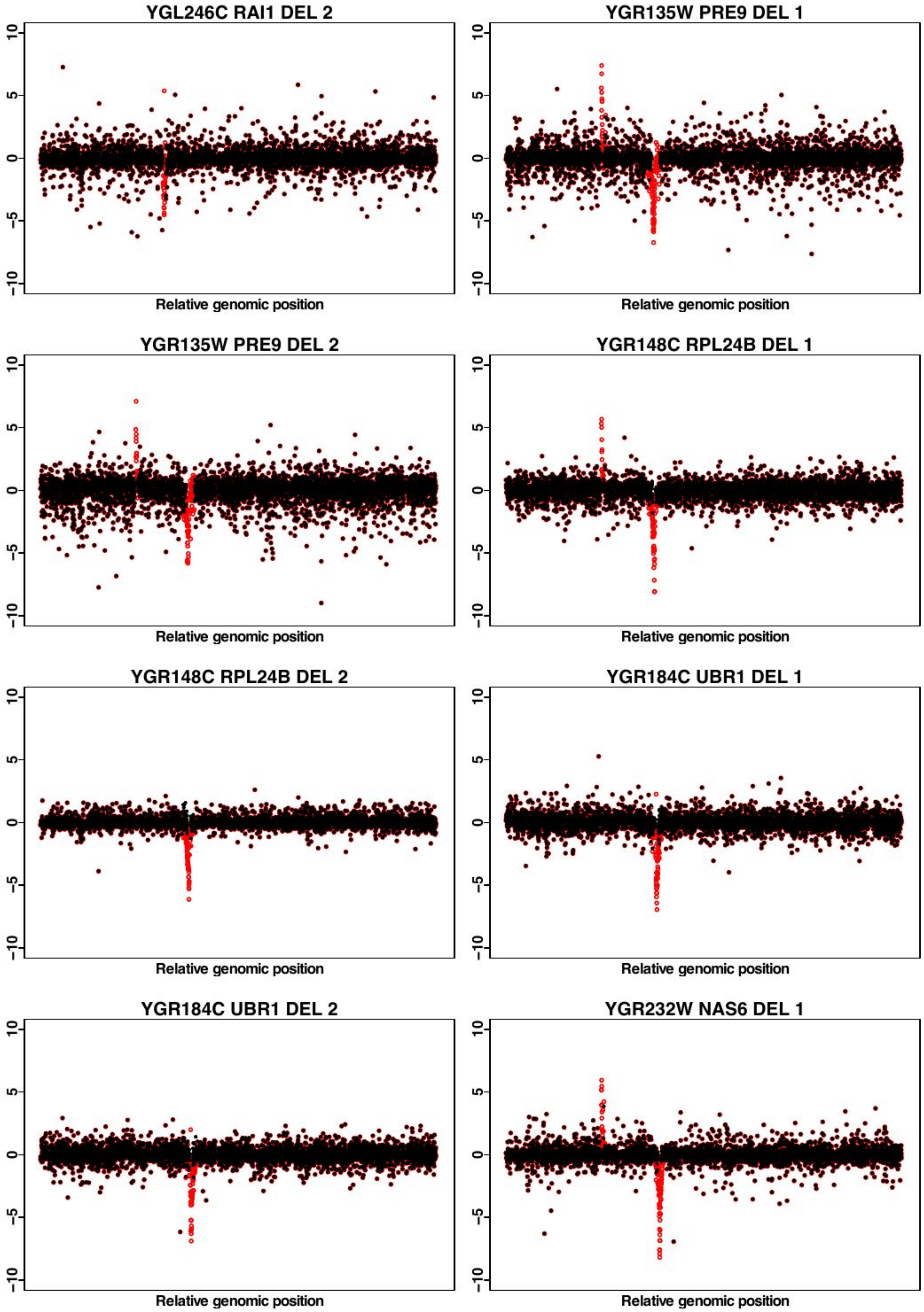
YGL246C RAI1 DEL 2
YGR135W PRE9 DEL 1
YGR135W PRE9 DEL 2
YGR148C RPL24B DEL 1
YGR148C RPL24B DEL 2
YGR184C UBR1 DEL 1
YGR184C UBR1 DEL 2
YGR232W NAS6 DEL 1
-10
-5
0
5
10
Relative genomic position

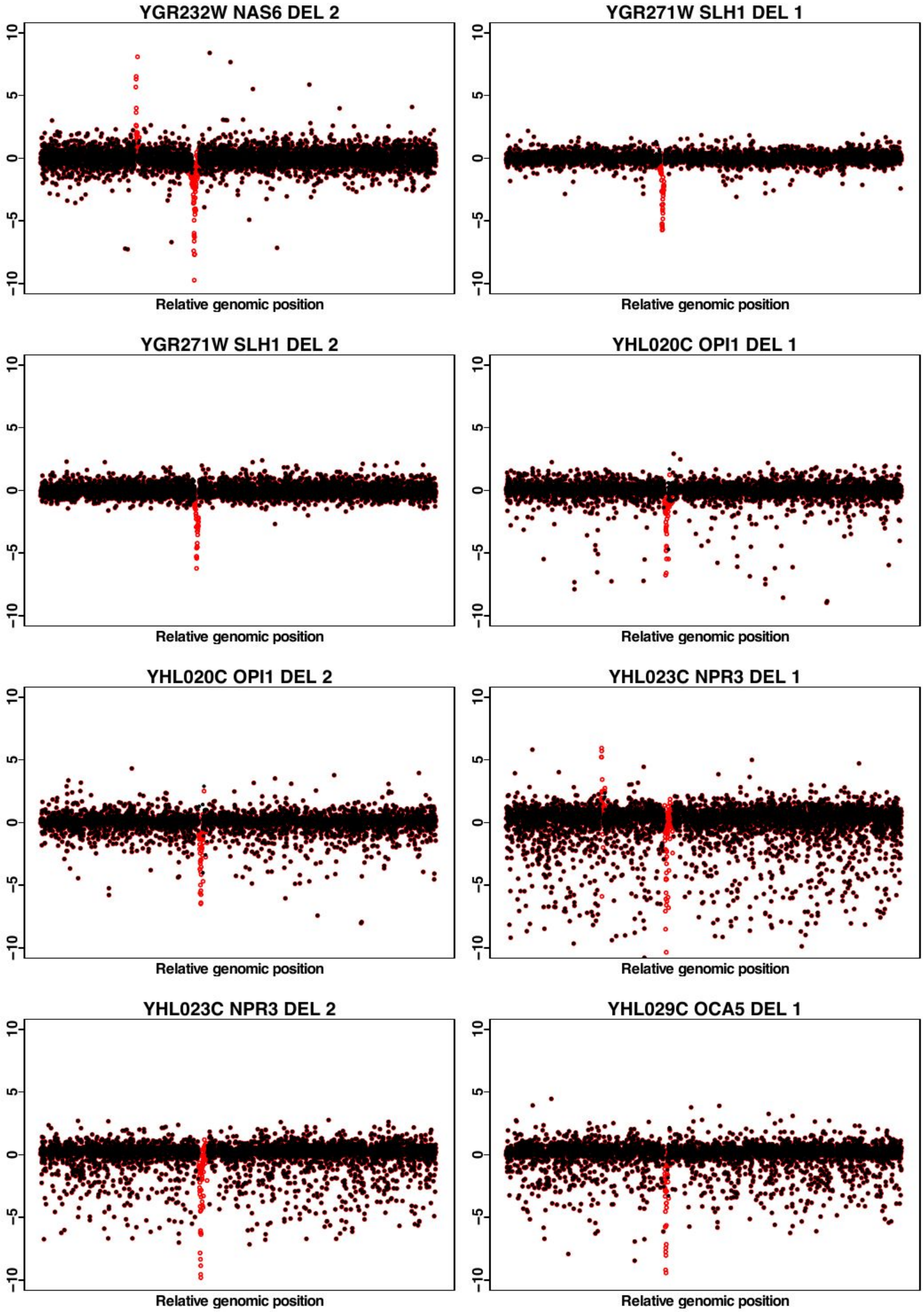

YGR232W NAS6 DEL 2
YGR271W SLH1 DEL 1
YGR271W SLH1 DEL 2
YHL020C OPI1 DEL 1
YHL020C OPI1 DEL 2
YHL023C NPR3 DEL 1
YHL023C NPR3 DEL 2
YHL029C OCA5 DEL 1
10
5
0
-5
-10
Relative genomic position

**YHL029C OCA5 DEL 2**

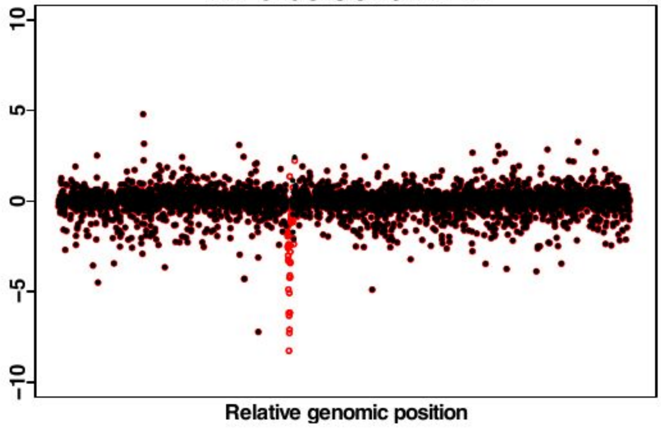


**YHR029C YHI9 DEL 1**

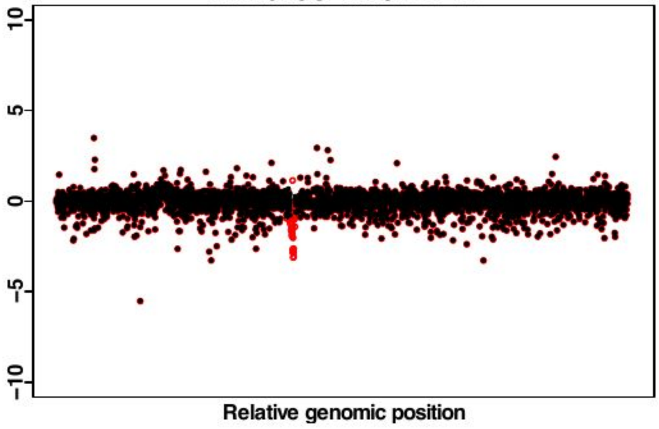


**YHR029C YHI9 DEL 2**

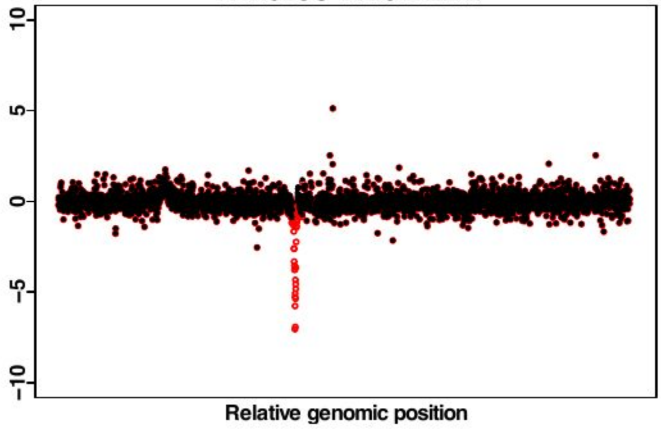


**YHR077C NMD2 DEL 1**

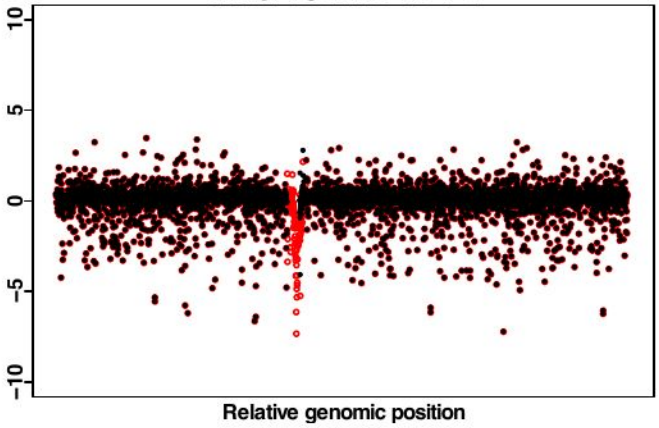


**YHR077C NMD2 DEL 2**

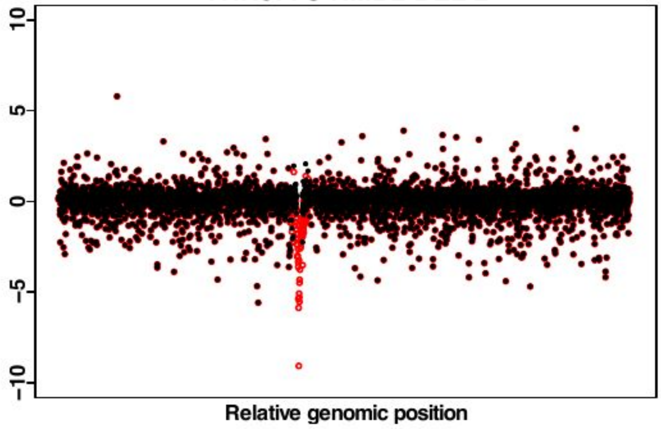


**YHR086W NAM8 DEL 1**

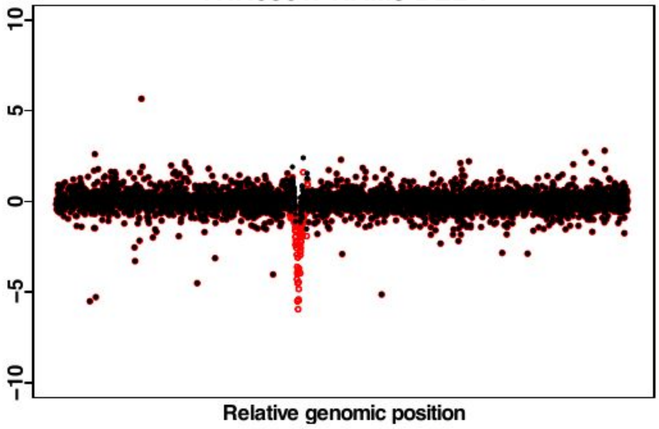


**YHR086W NAM8 DEL 2**

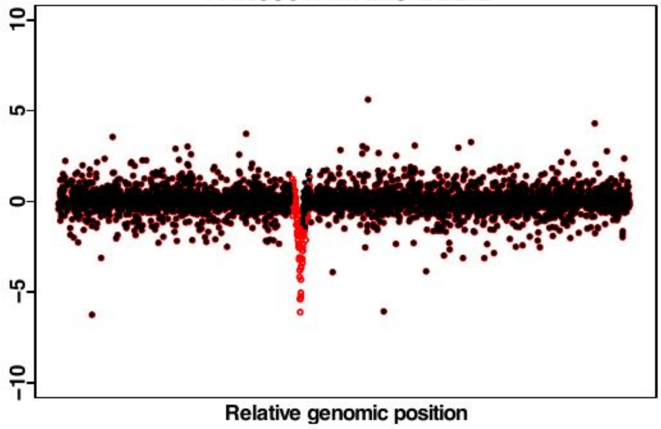


**YHR087W RTC3 DEL 1**

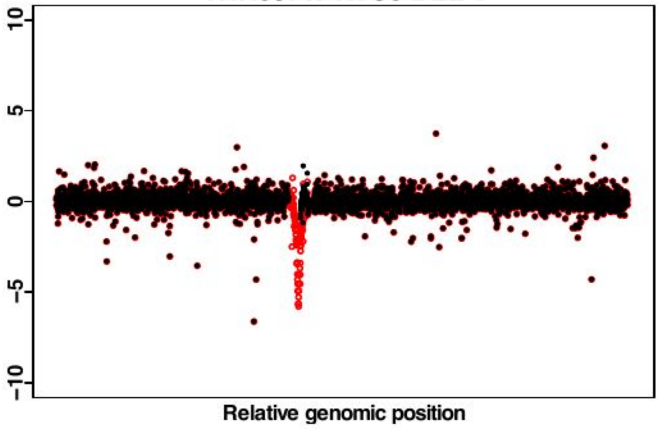

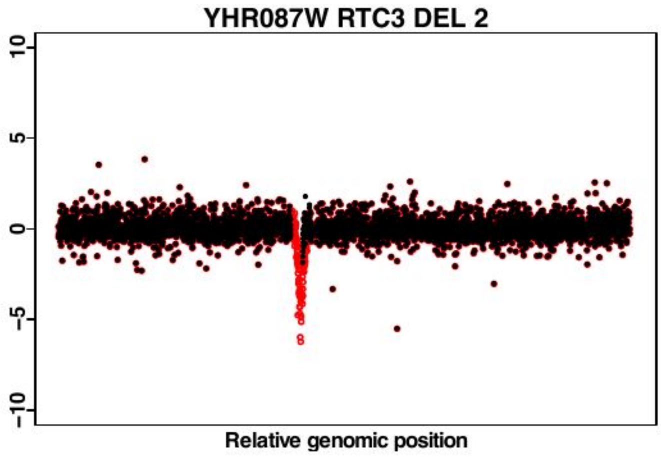
YHR087W RTC3 DEL 2
10
5
0
-5
-10
Relative genomic position

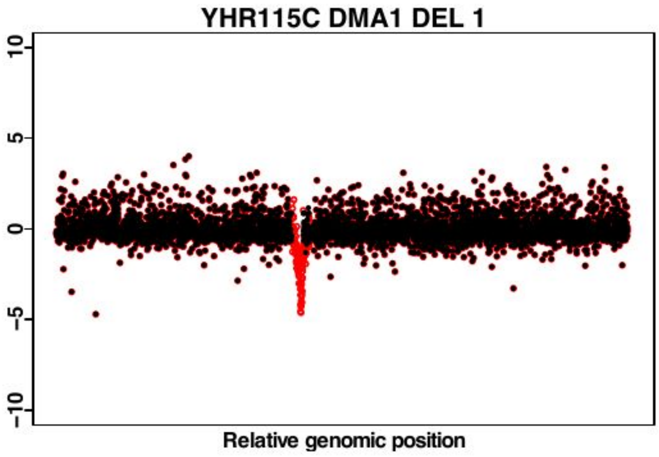
YHR115C DMA1 DEL 1
10
5
0
-5
-10
Relative genomic position

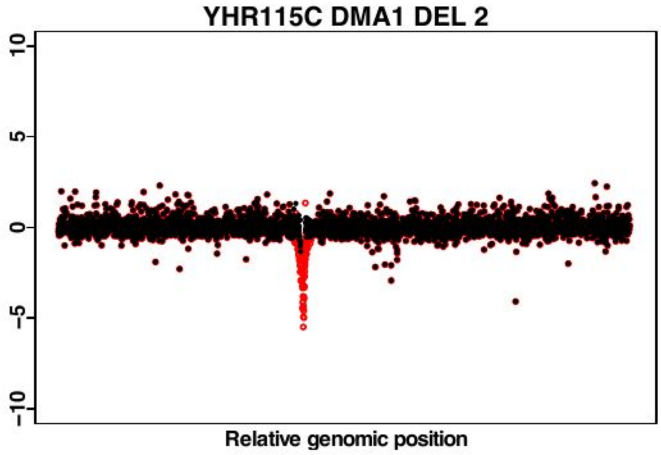
YHR115C DMA1 DEL 2
10
5
0
-5
-10
Relative genomic position

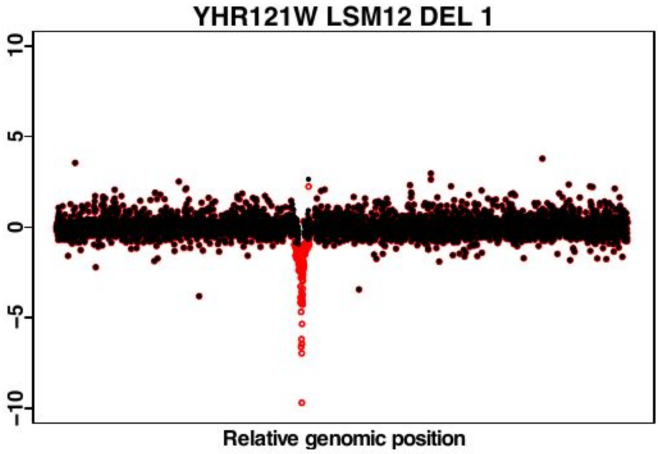
YHR121W LSM12 DEL 1
10
5
0
-5
-10
Relative genomic position

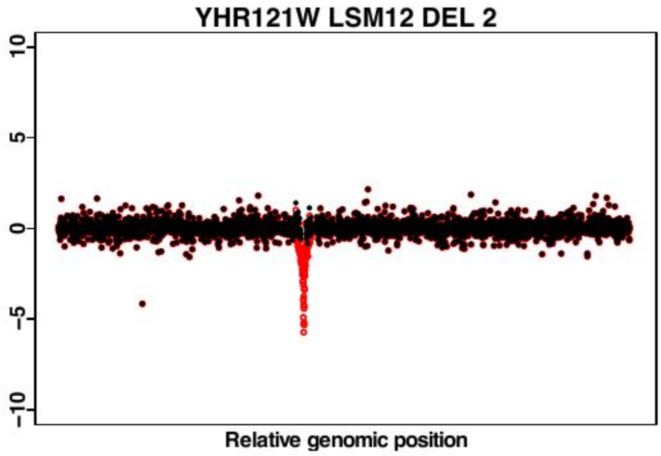
YHR121W LSM12 DEL 2
10
5
0
-5
-10
Relative genomic position

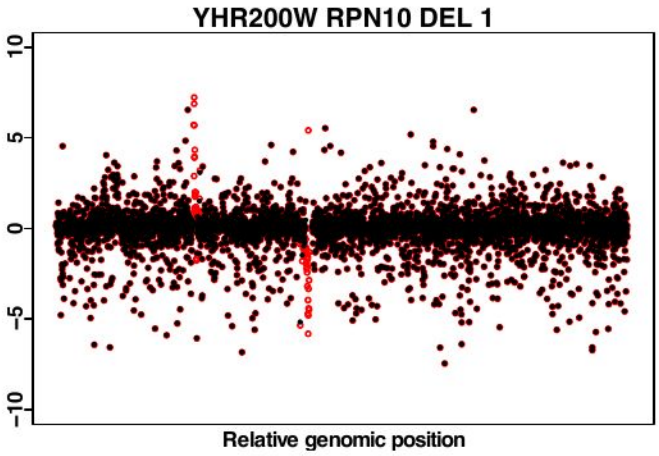
YHR200W RPN10 DEL 1
10
5
0
-5
-10
Relative genomic position

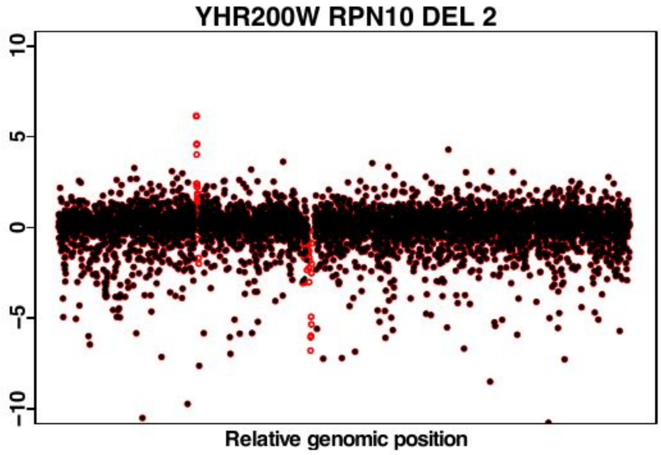
YHR200W RPN10 DEL 2
10
5
0
-5
-10
Relative genomic position

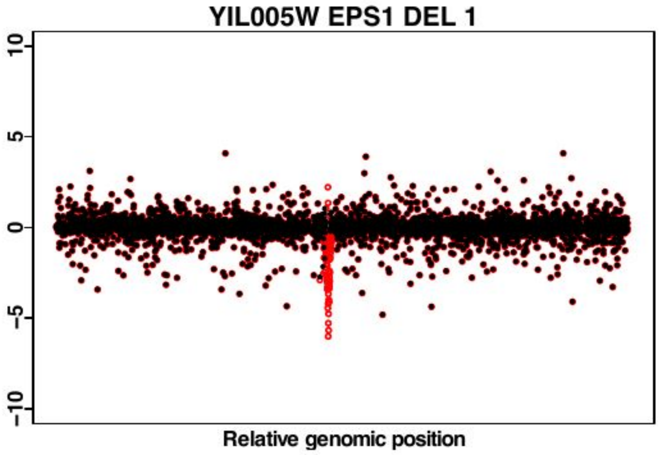
YIL005W EPS1 DEL 1
10
5
0
-5
-10
Relative genomic position

YIL005W EPS1 DEL 2

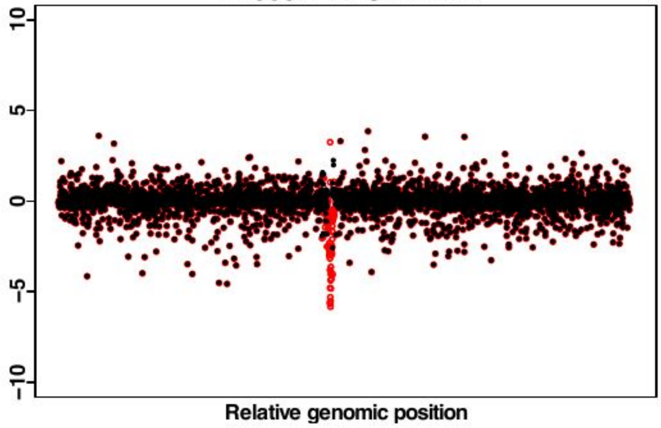


YIL007C NAS2 DEL 1

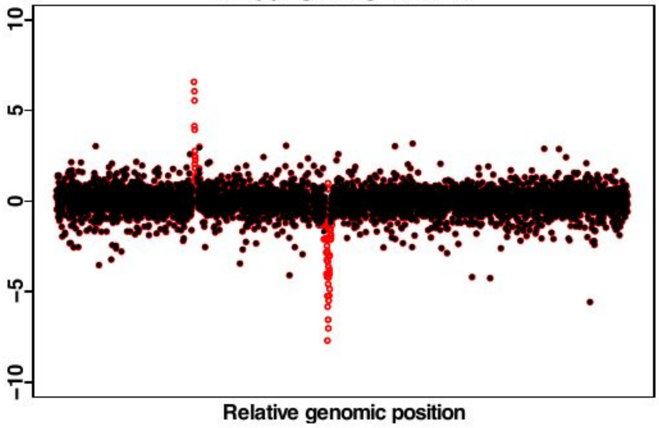


YIL007C NAS2 DEL 2

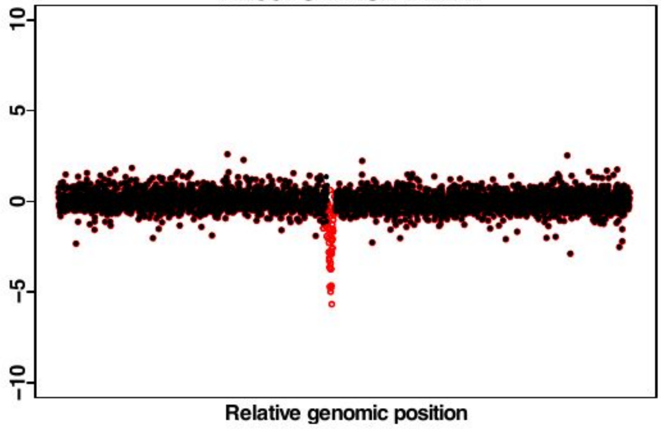


YIL096C YIL096C DEL 1

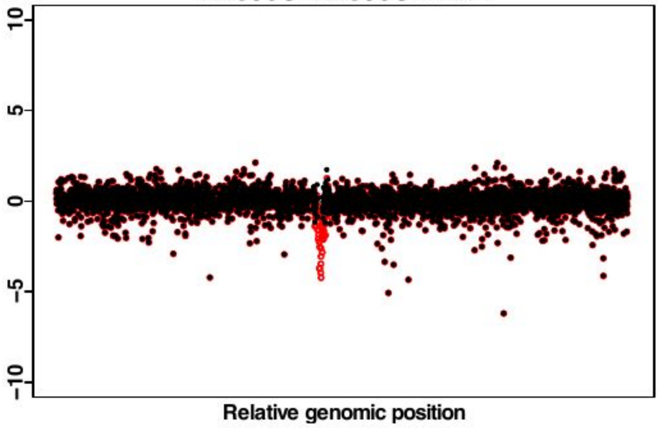


YIL096C YIL096C DEL 2

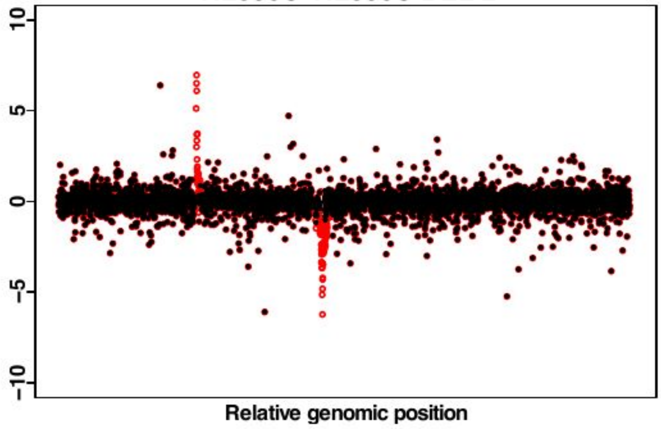


YIL104C SHQ1 DAMP 1

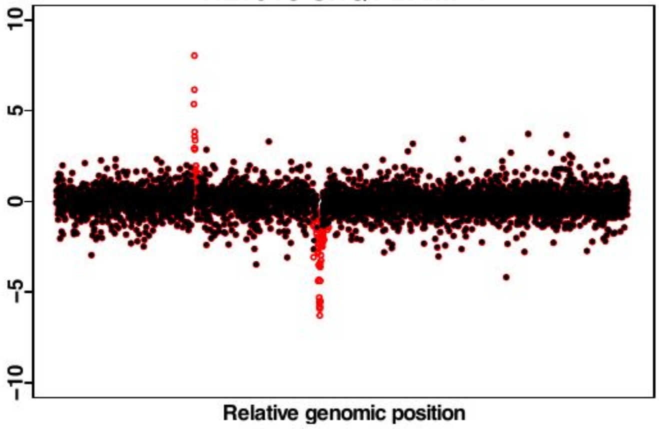


YIL104C SHQ1 DAMP 2

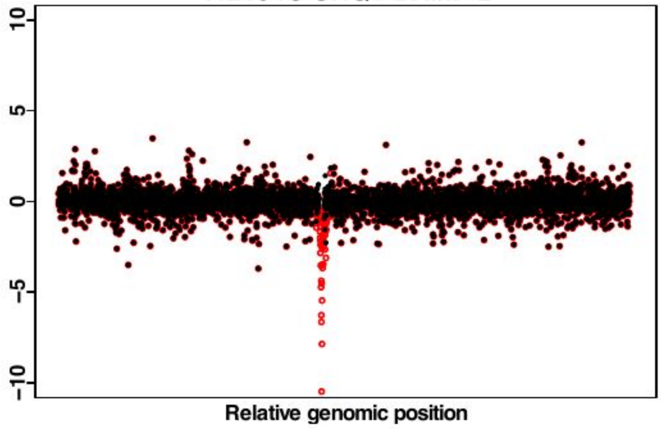


YJL128C PBS2 DEL 1

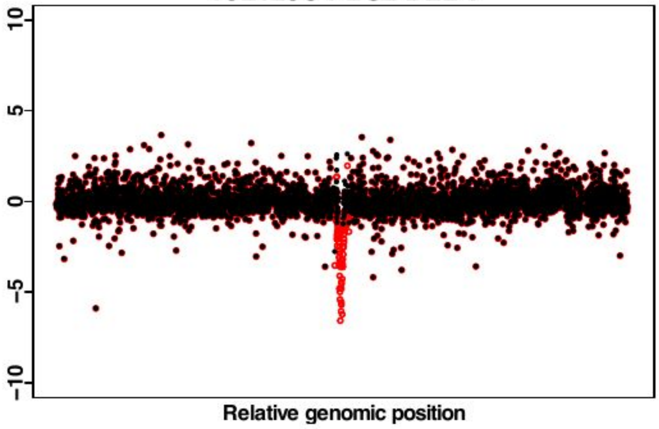

YJL128C PBS2 DEL 2

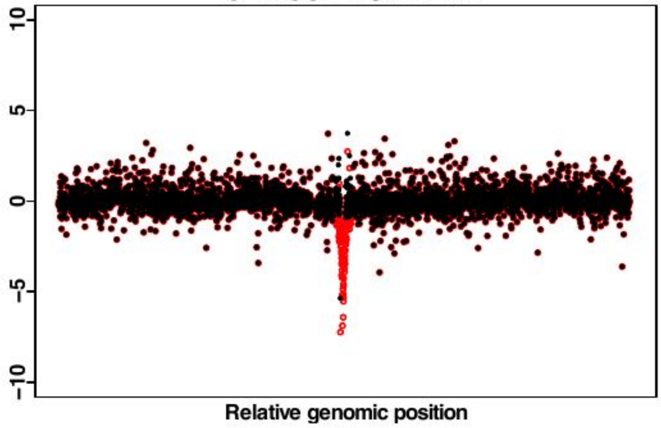


YJL141C YAK1 DEL 1

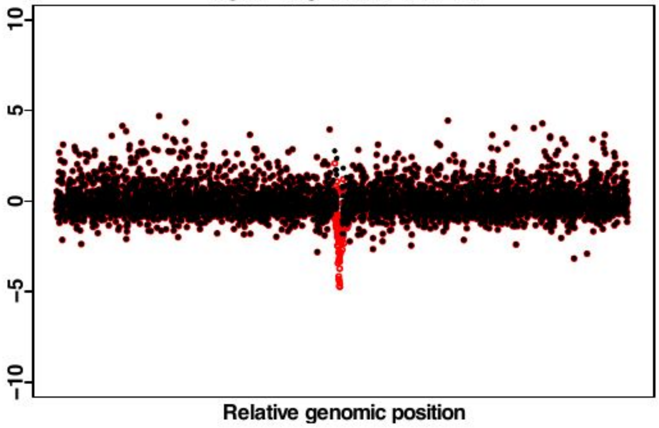


YJL141C YAK1 DEL 2

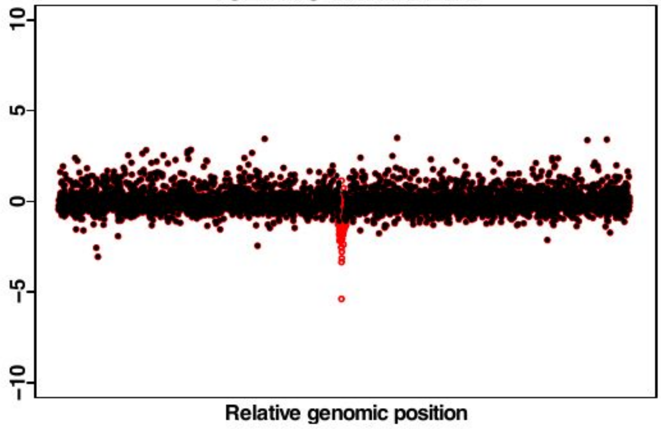


YJL191W RPS14B DEL 1

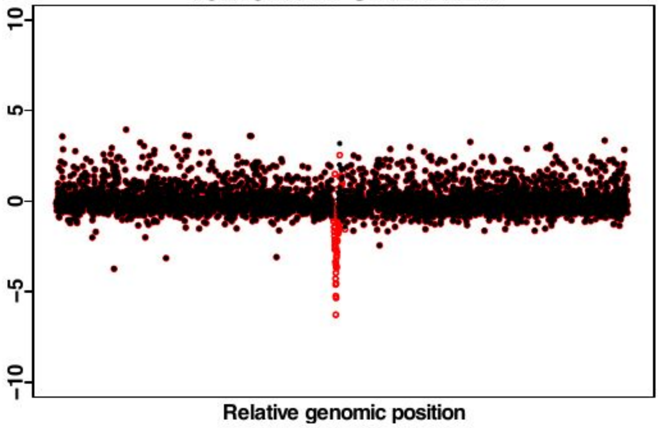


YJL191W RPS14B DEL 2

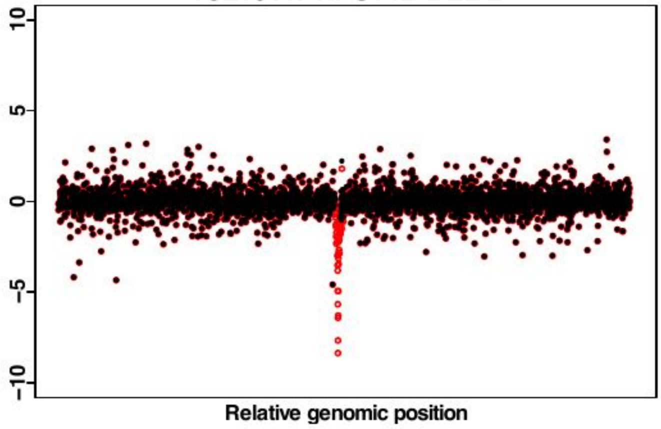


YJL208C NUC1 DEL 1

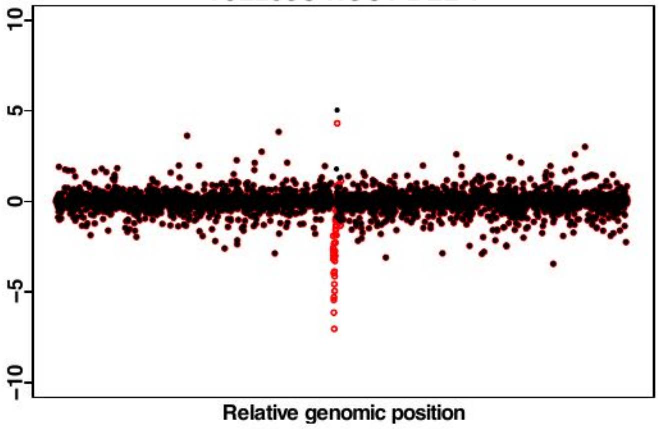


YJL208C NUC1 DEL 2

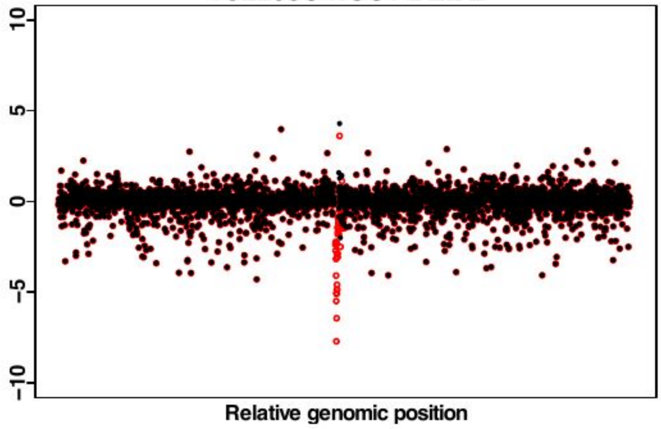


YJR070C LIA1 DEL 1

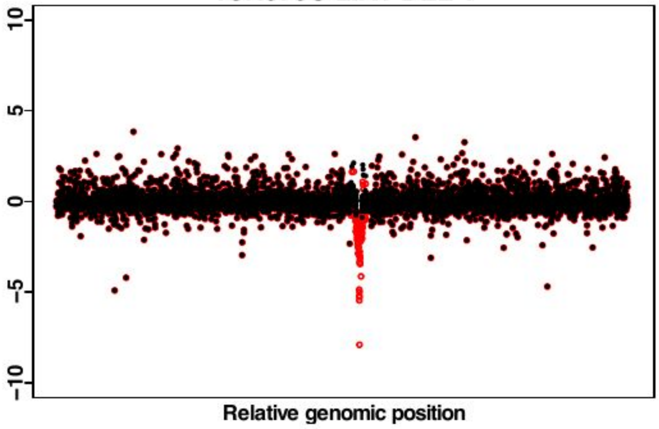

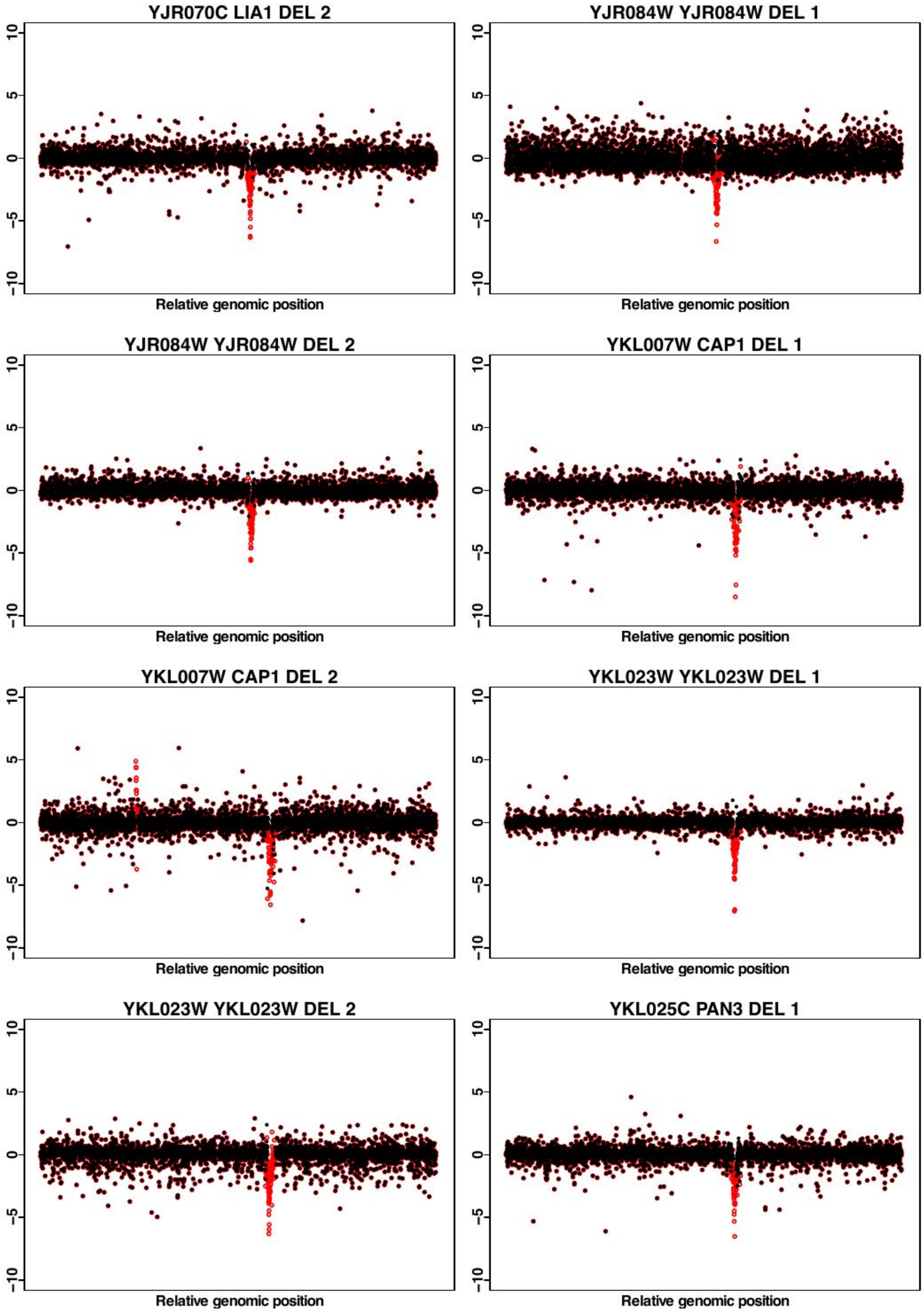
YJR070C LIA1 DEL 2
YJR084W YJR084W DEL 1
YJR084W YJR084W DEL 2
YKL007W CAP1 DEL 1
YKL007W CAP1 DEL 2
YKL023W YKL023W DEL 1
YKL023W YKL023W DEL 2
YKL025C PAN3 DEL 1
10
5
0
−5
−10
Relative genomic position

YKL025C PAN3 DEL 2

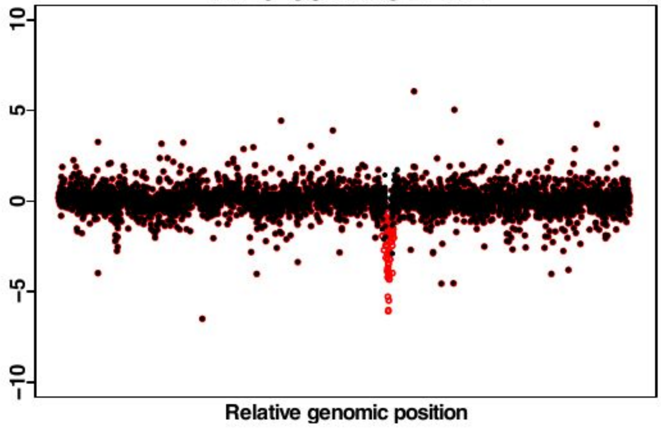


YKL081W TEF4 DEL 1

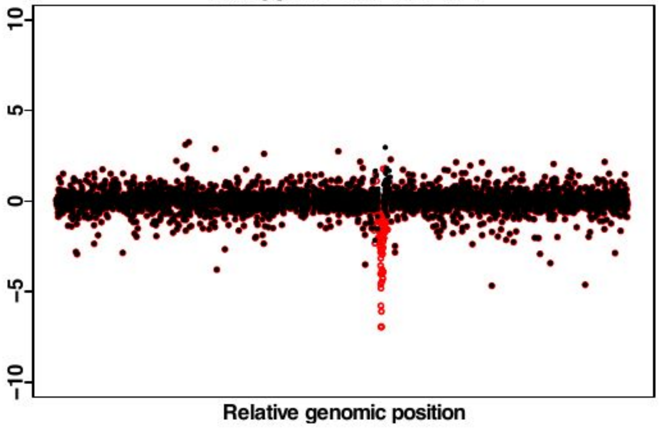


YKL081W TEF4 DEL 2

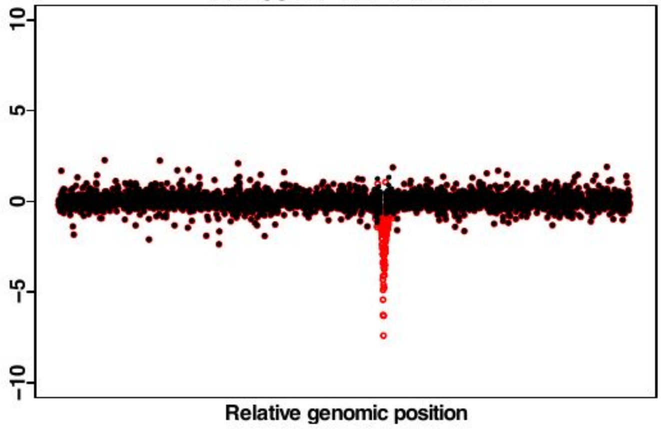


YKL206C ADD66 DEL 1

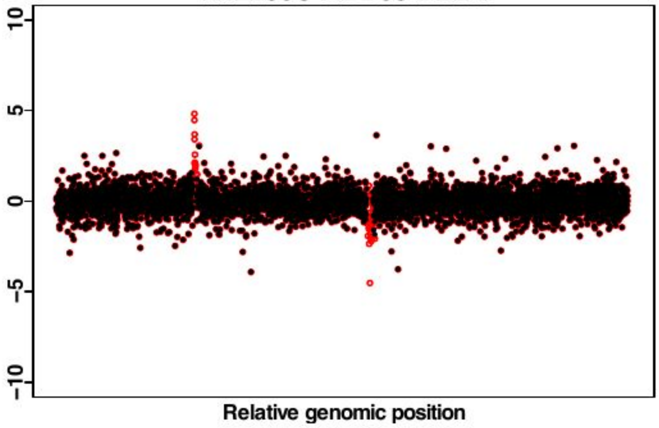


YKL206C ADD66 DEL 2

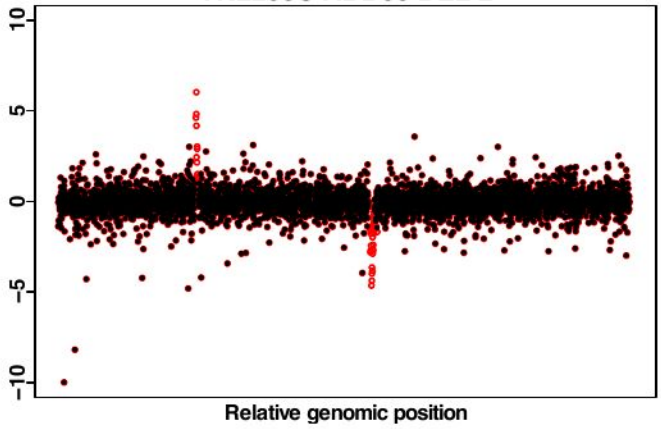


YKR019C IRS4 DEL 1

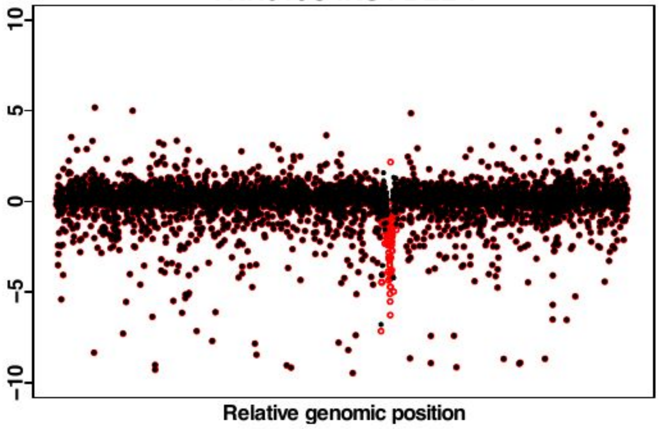


YKR019C IRS4 DEL 2

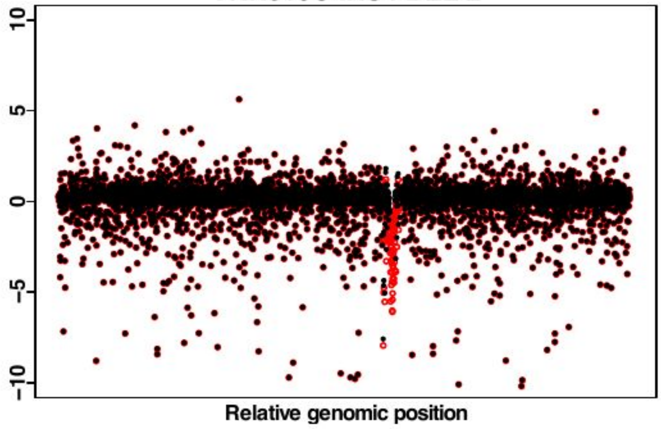


YKR048C NAP1 DEL 1

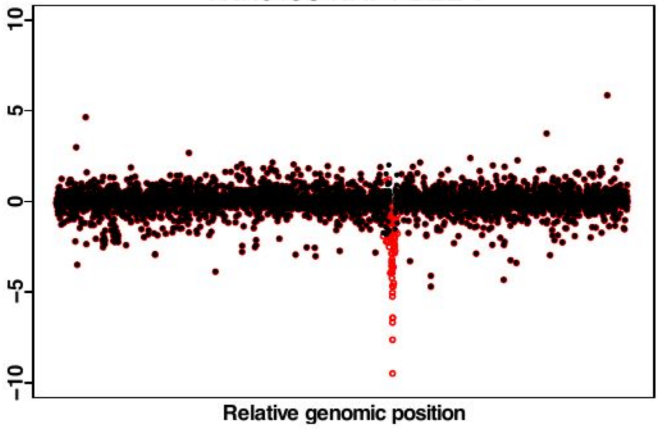

YKR048C NAP1 DEL 2

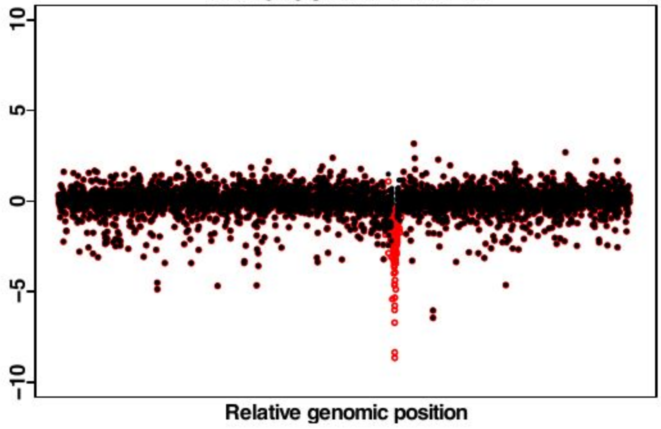


YKR084C HBS1 DEL 1

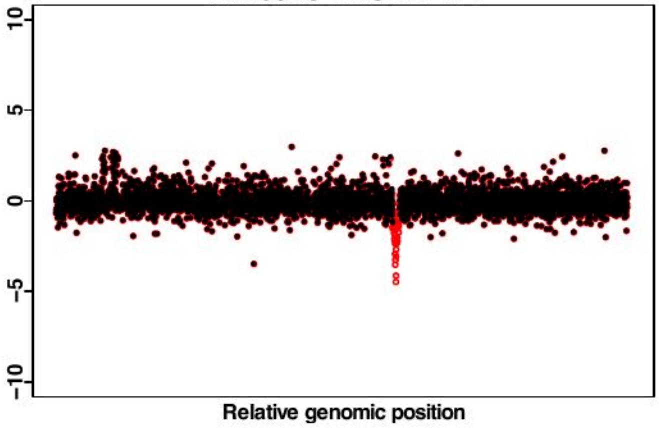


YKR084C HBS1 DEL 2

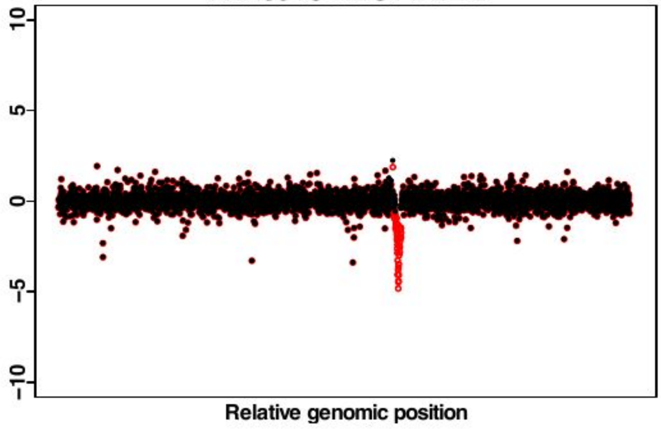


YLR021W IRC25 DEL 1

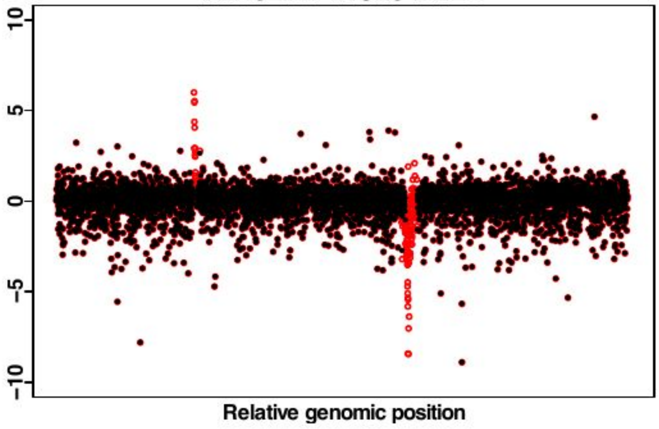


YLR021W IRC25 DEL 2

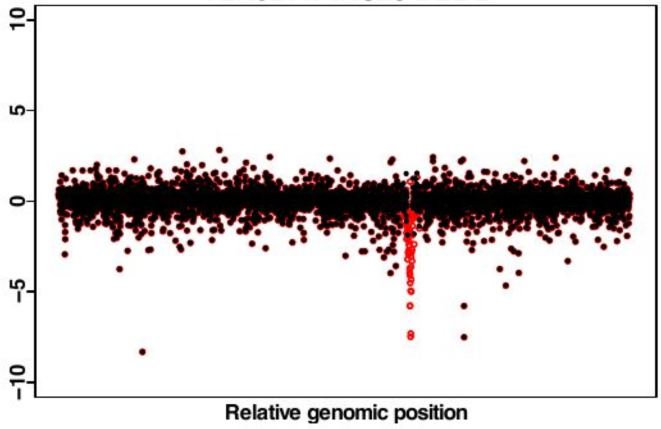


YLR058C SHM2 DEL 1

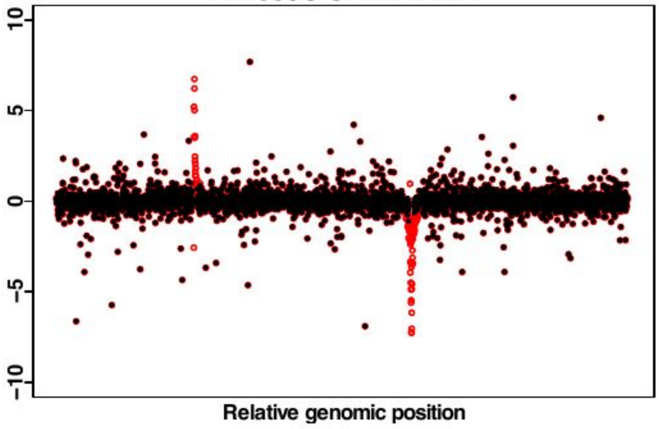


YLR058C SHM2 DEL 2

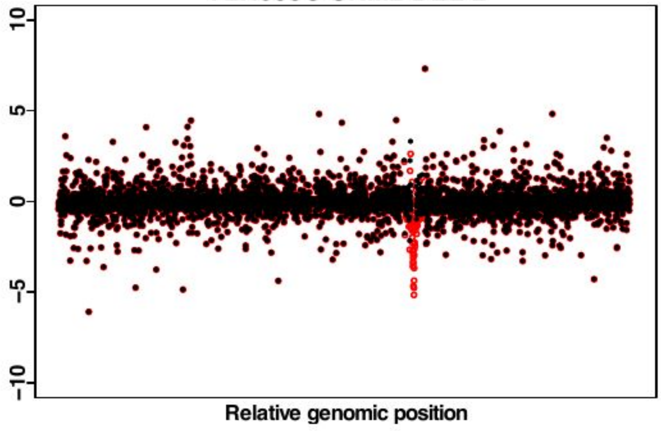


YLR059C REX2 DEL 1

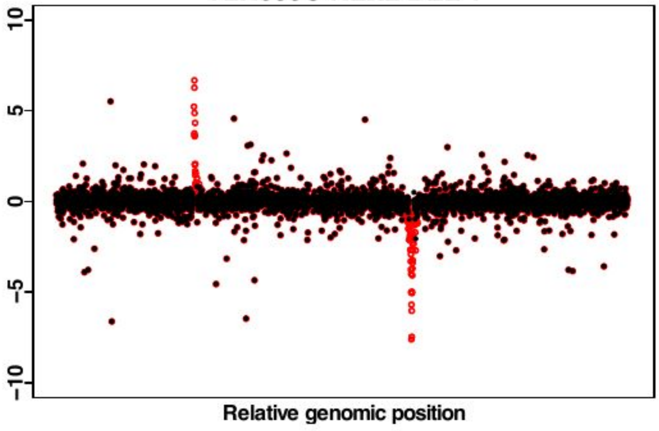

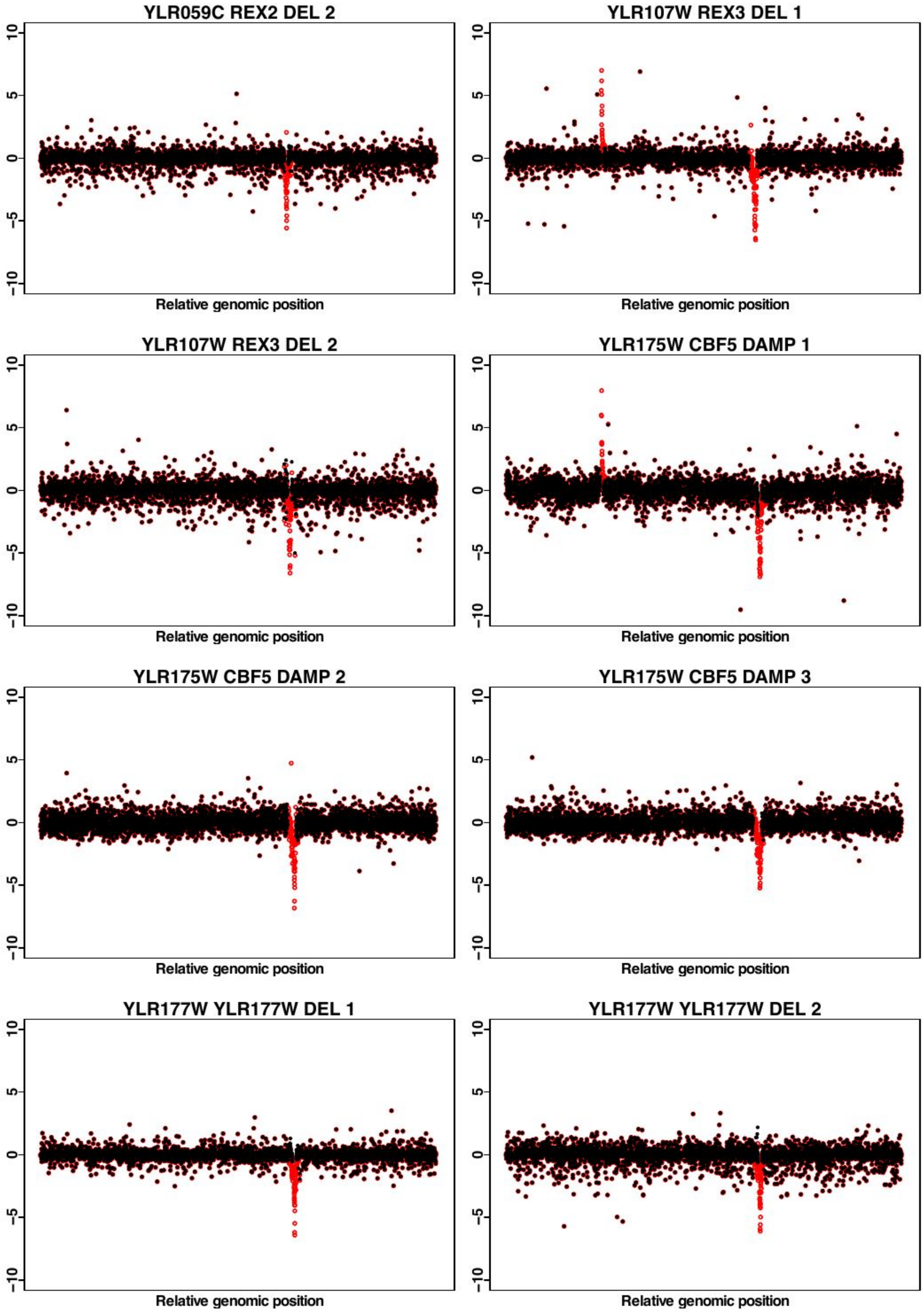
YLR059C REX2 DEL 2
YLR107W REX3 DEL 1
YLR107W REX3 DEL 2
YLR175W CBF5 DAMP 1
YLR175W CBF5 DAMP 2
YLR175W CBF5 DAMP 3
YLR177W YLR177W DEL 1
YLR177W YLR177W DEL 2
Relative genomic position
10
5
0
−5
−10

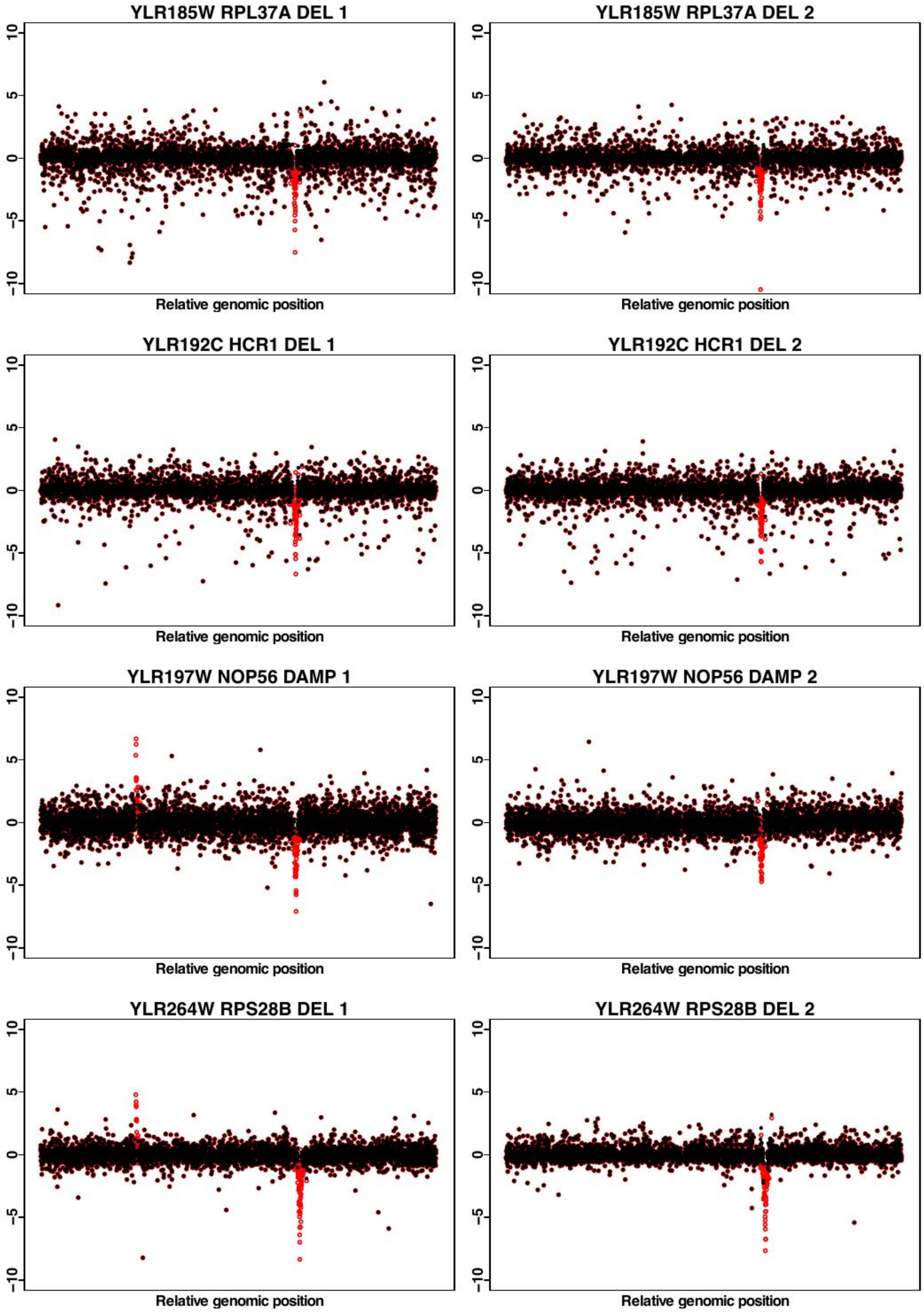
YLR185W RPL37A DEL 1
YLR185W RPL37A DEL 2
YLR192C HCR1 DEL 1
YLR192C HCR1 DEL 2
YLR197W NOP56 DAMP 1
YLR197W NOP56 DAMP 2
YLR264W RPS28B DEL 1
YLR264W RPS28B DEL 2
10
5
0
−5
−10
Relative genomic position

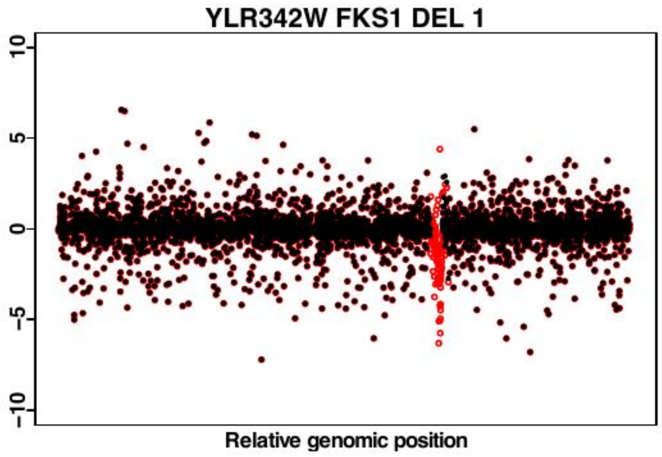
YLR342W FKS1 DEL 1
10
5
0
-5
-10
Relative genomic position

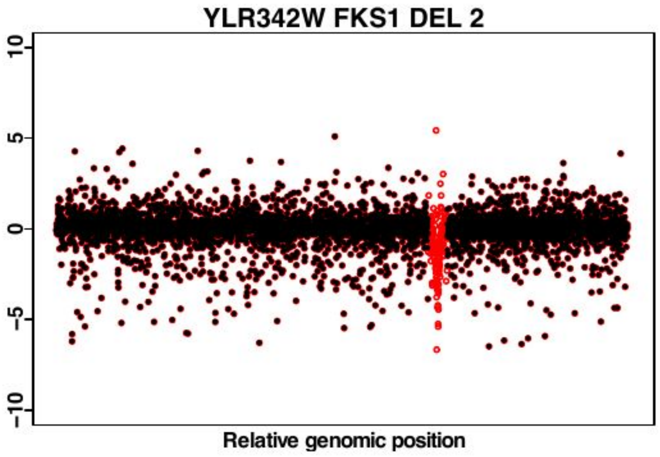
YLR342W FKS1 DEL 2
10
5
0
-5
-10
Relative genomic position

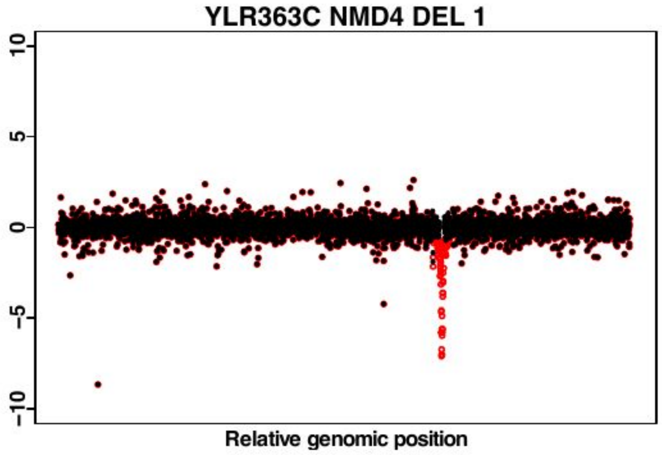
YLR363C NMD4 DEL 1
10
5
0
-5
-10
Relative genomic position

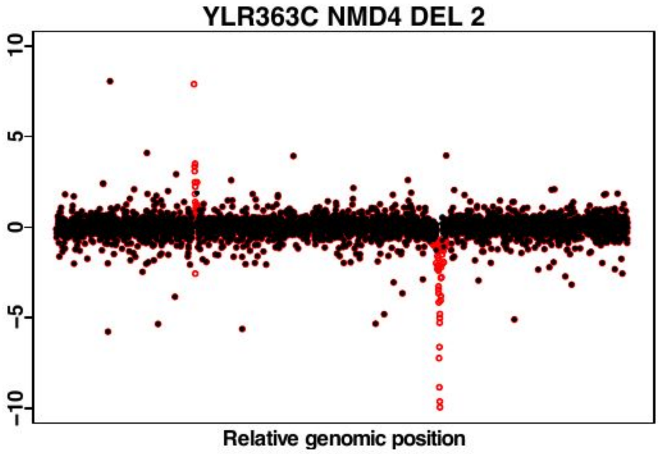
YLR363C NMD4 DEL 2
10
5
0
-5
-10
Relative genomic position

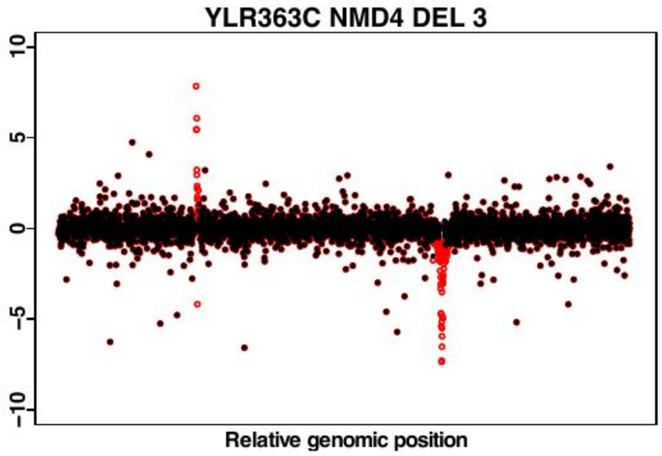
YLR363C NMD4 DEL 3
10
5
0
-5
-10
Relative genomic position

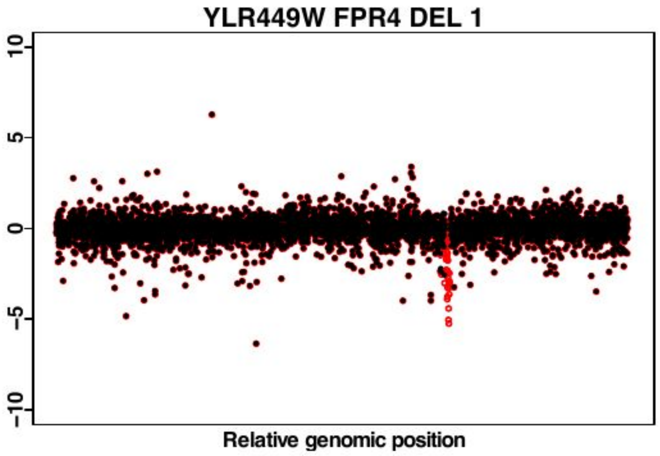
YLR449W FPR4 DEL 1
10
5
0
-5
-10
Relative genomic position

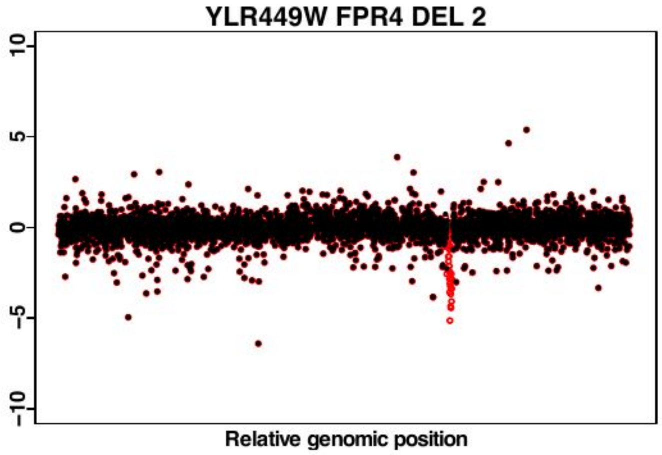
YLR449W FPR4 DEL 2
10
5
0
-5
-10
Relative genomic position

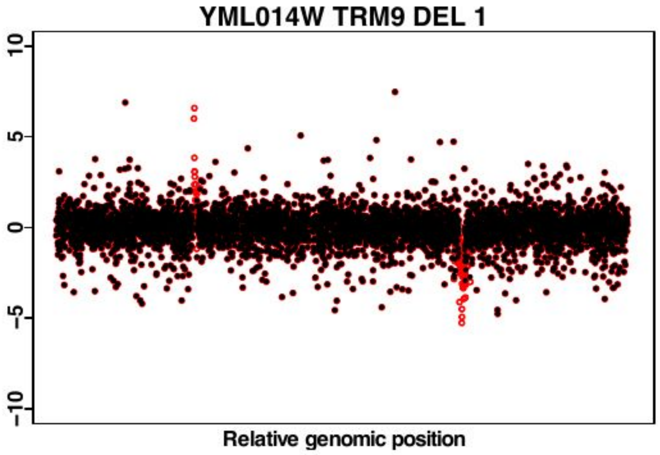
YML014W TRM9 DEL 1
10
5
0
-5
-10
Relative genomic position

YML014W TRM9 DEL 2

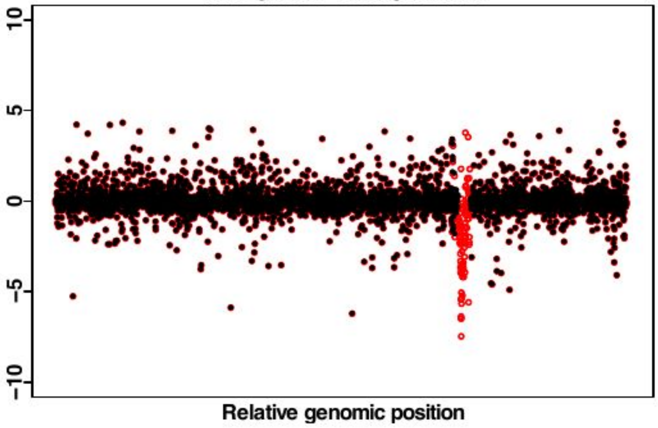


YML058W.A HUG1 DEL 1

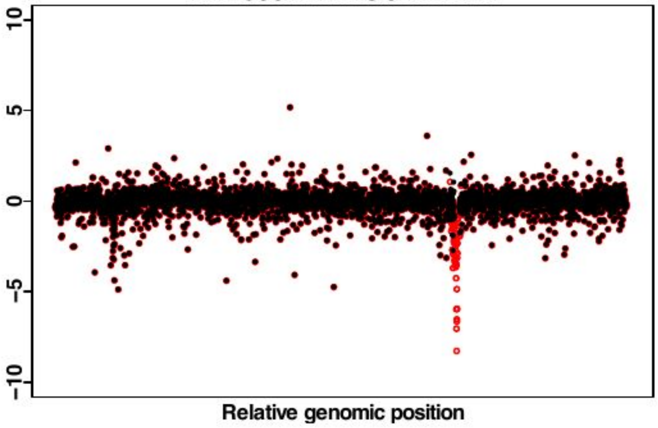


YML058W.A HUG1 DEL 2

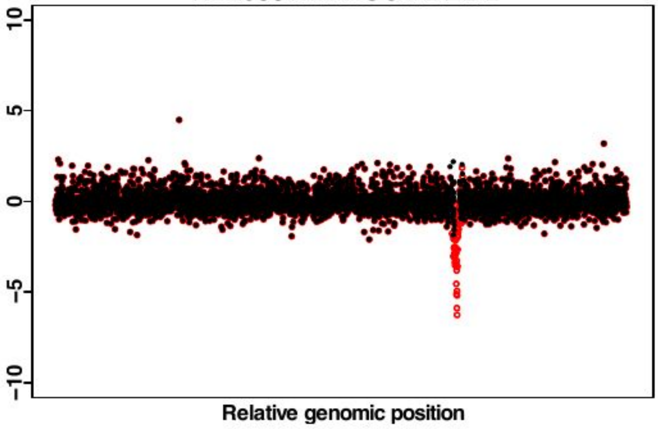


YML074C FPR3 DEL 1

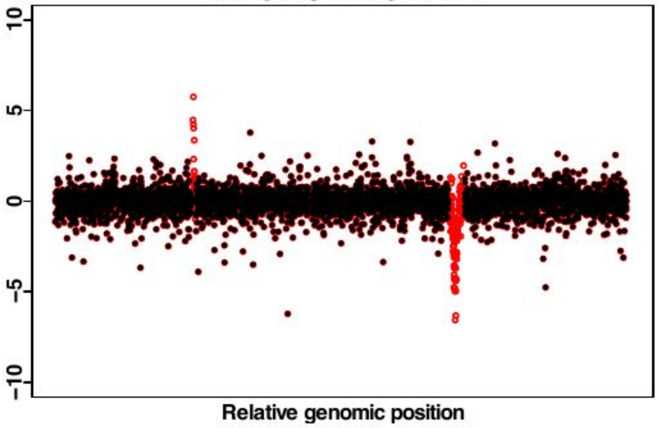


YML074C FPR3 DEL 2

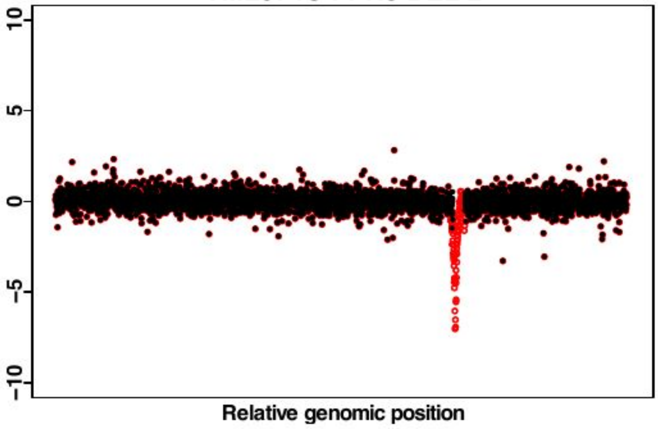


YML074C FPR3 DEL 3

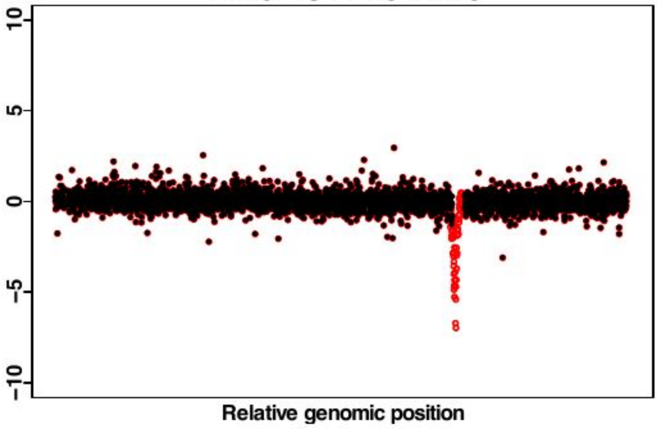


YMR037C MSN2 DEL 1

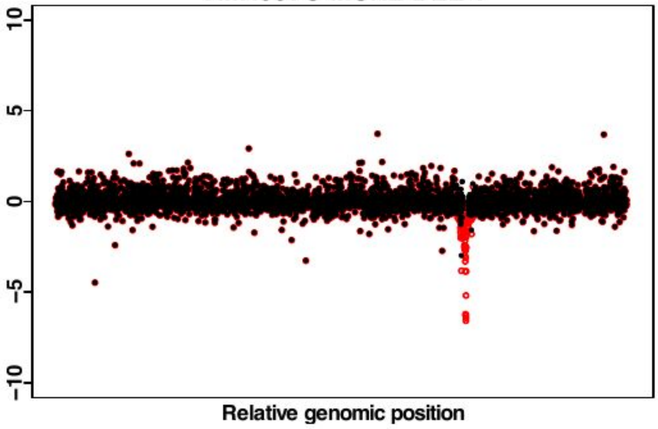


YMR037C MSN2 DEL 2

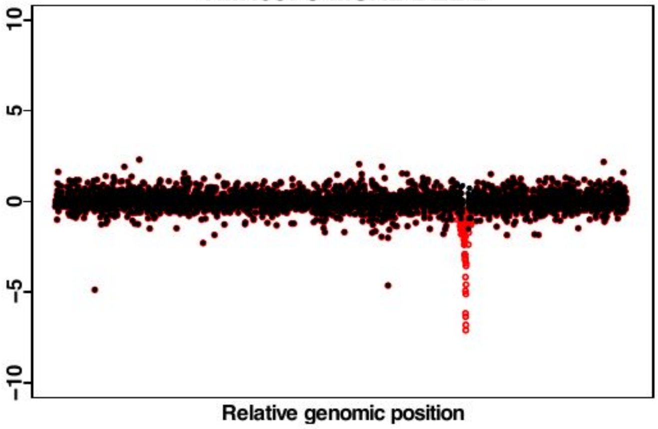

YMR075W RCO1 DEL 1

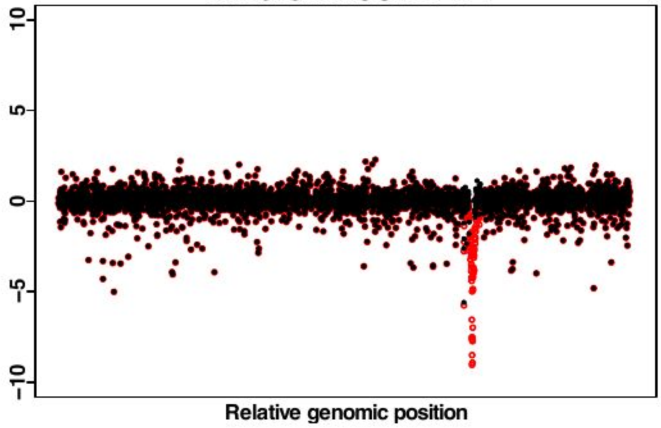


YMR075W RCO1 DEL 2

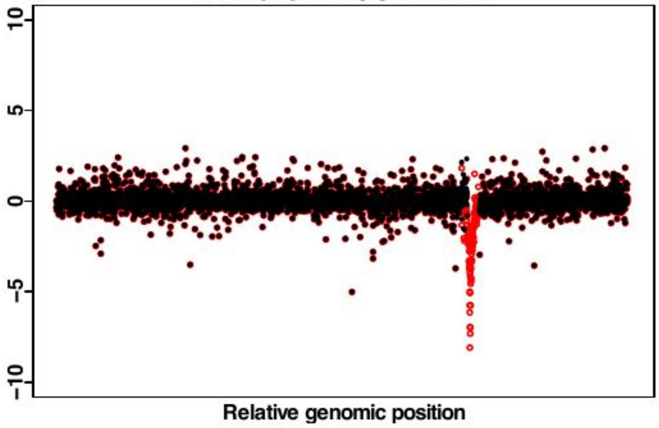


YMR080C NAM7 DEL 1

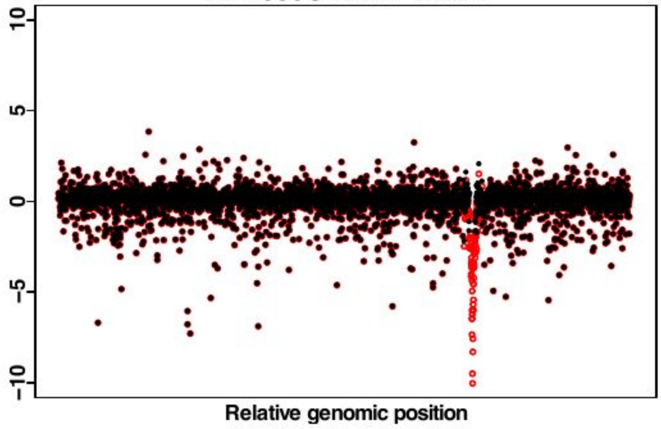


YMR080C NAM7 DEL 2

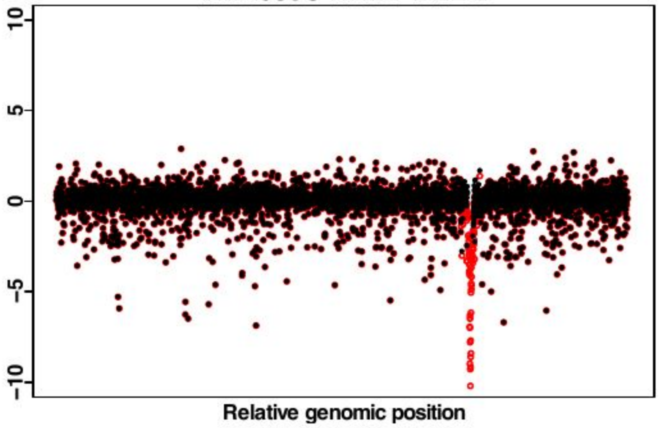


YMR080C NAM7 DEL 3

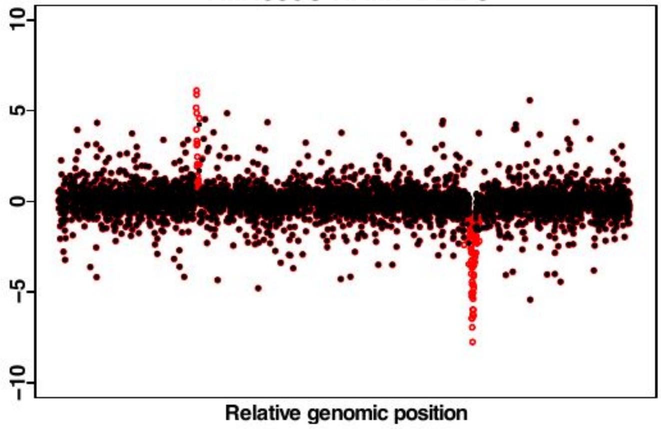


YMR153W NUP53 DEL 1

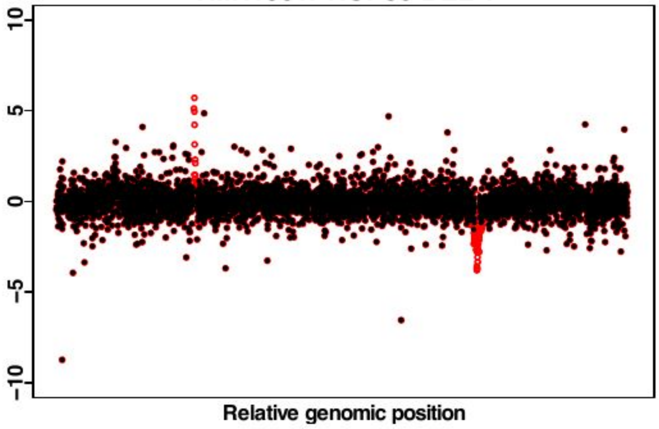


YMR153W NUP53 DEL 2

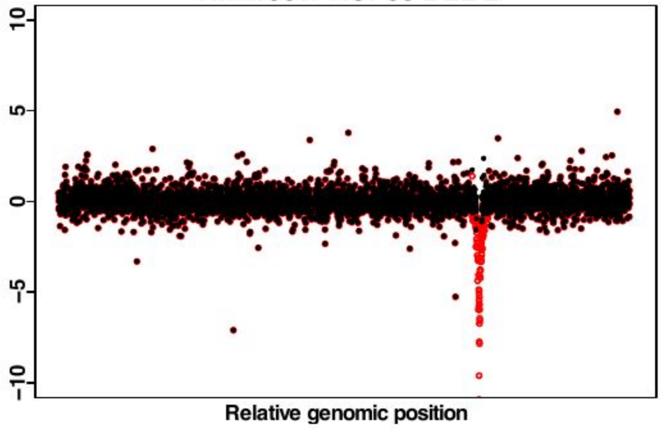


YMR162C DNF3 DEL 1

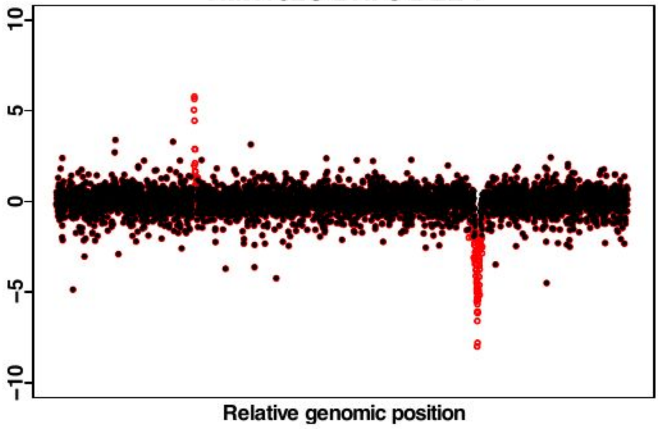

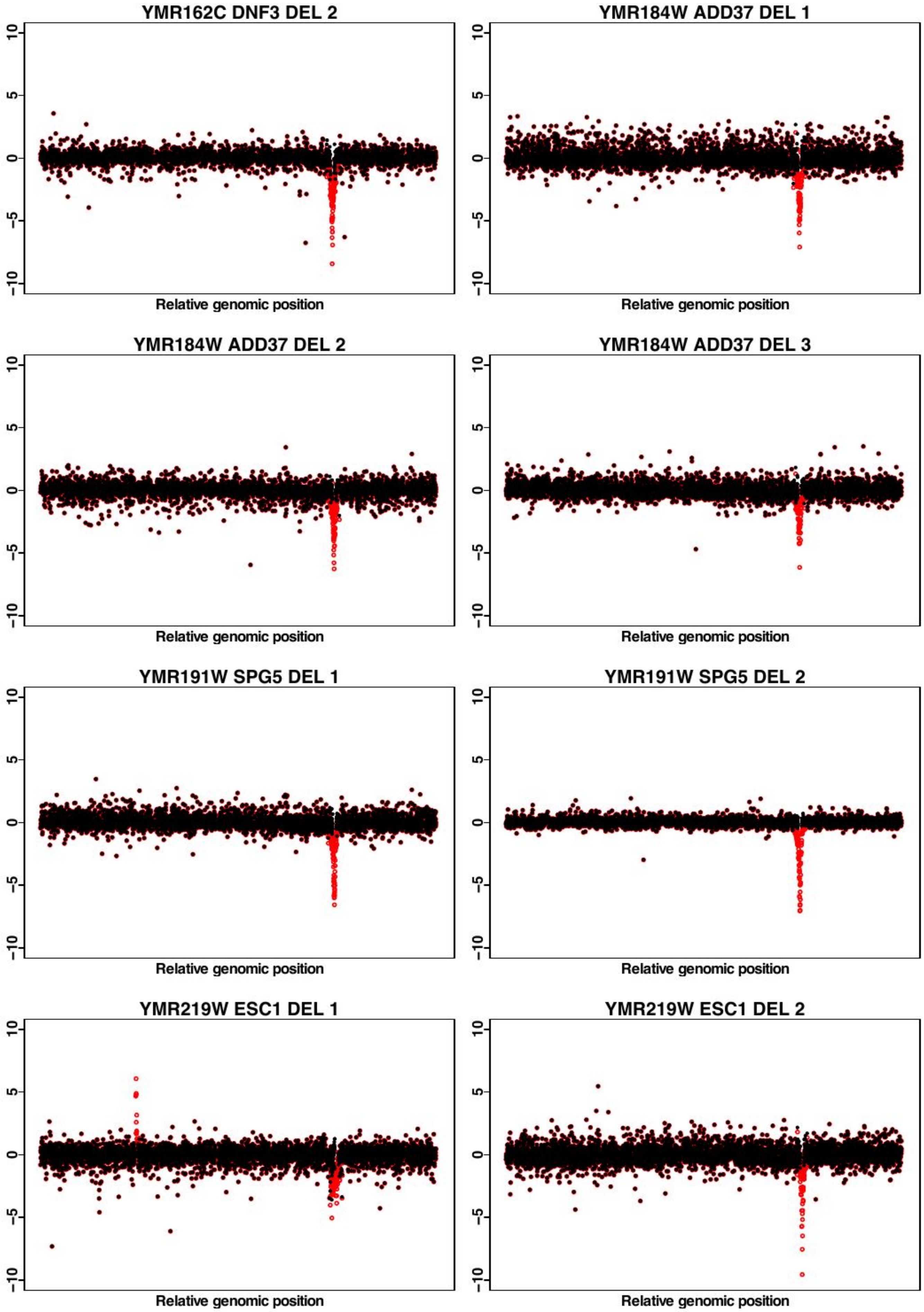
YMR162C DNF3 DEL 2
YMR184W ADD37 DEL 1
YMR184W ADD37 DEL 2
YMR184W ADD37 DEL 3
YMR191W SPG5 DEL 1
YMR191W SPG5 DEL 2
YMR219W ESC1 DEL 1
YMR219W ESC1 DEL 2
10
5
0
-5
-10
Relative genomic position

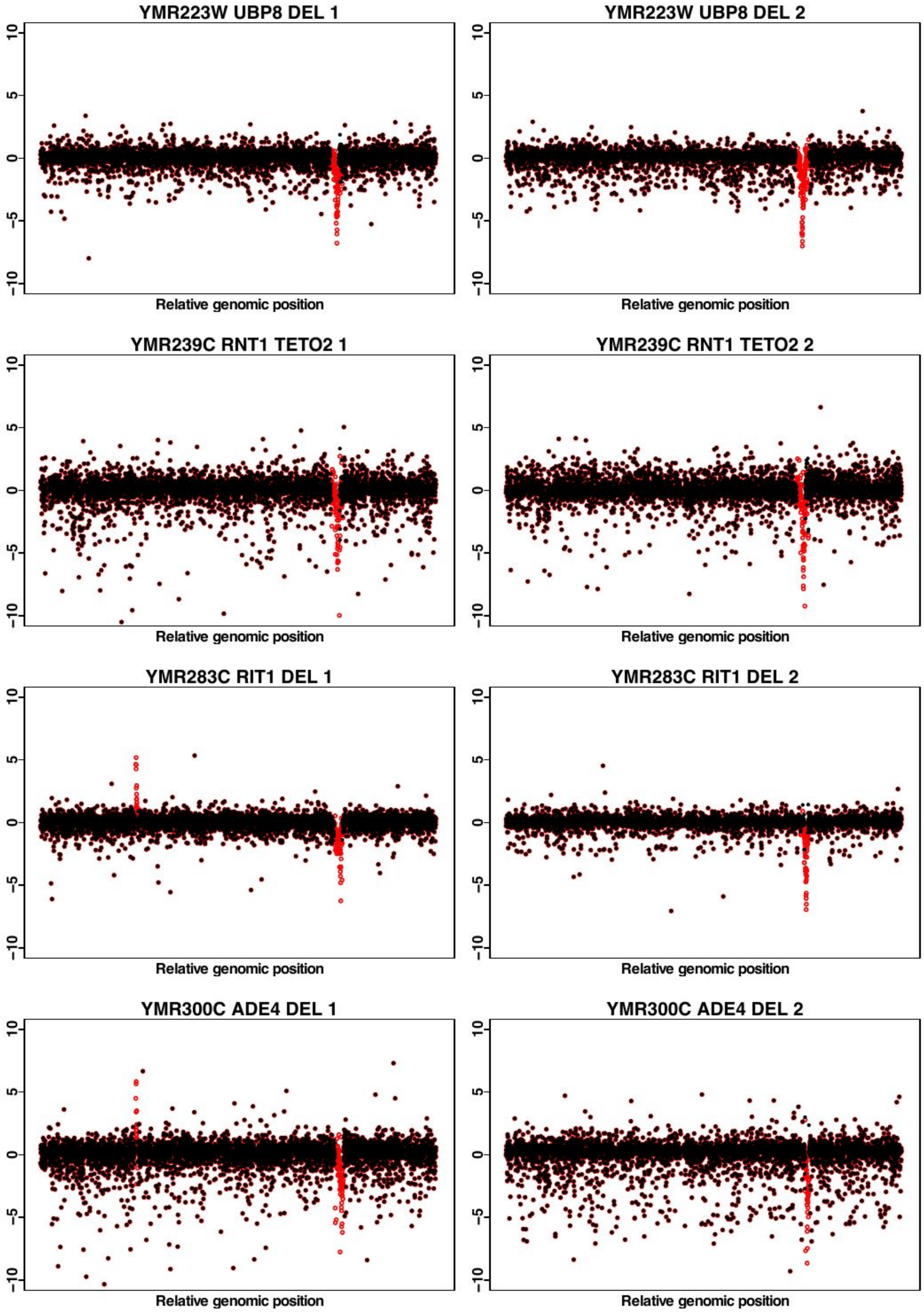

YMR223W UBP8 DEL 1
YMR223W UBP8 DEL 2
YMR239C RNT1 TETO2 1
YMR239C RNT1 TETO2 2
YMR283C RIT1 DEL 1
YMR283C RIT1 DEL 2
YMR300C ADE4 DEL 1
YMR300C ADE4 DEL 2
10
5
0
-5
-10
Relative genomic position

**YNL021W HDA1 DEL 1**

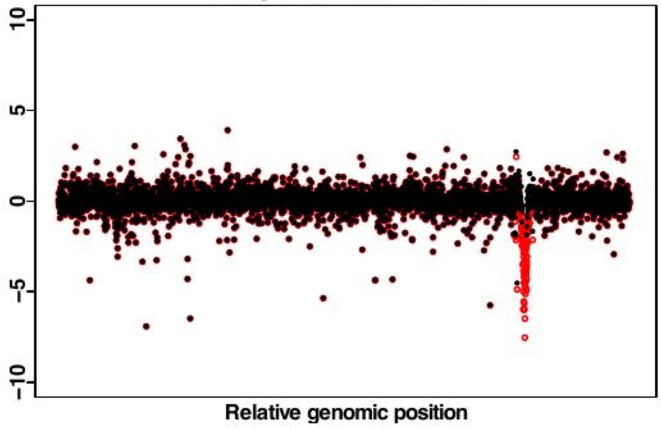


**YNL021W HDA1 DEL 2**

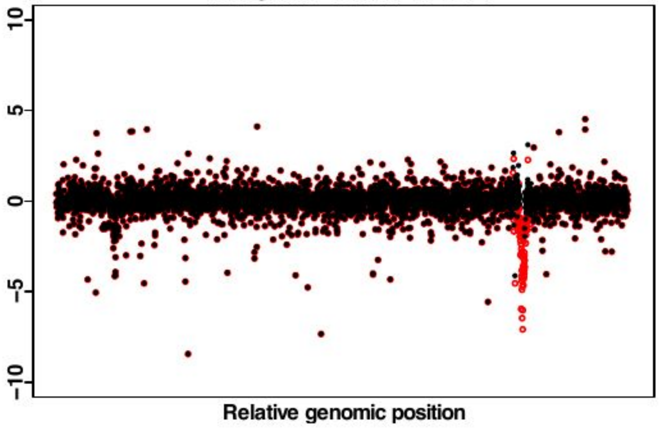


**YNL056W OCA2 DEL 1**

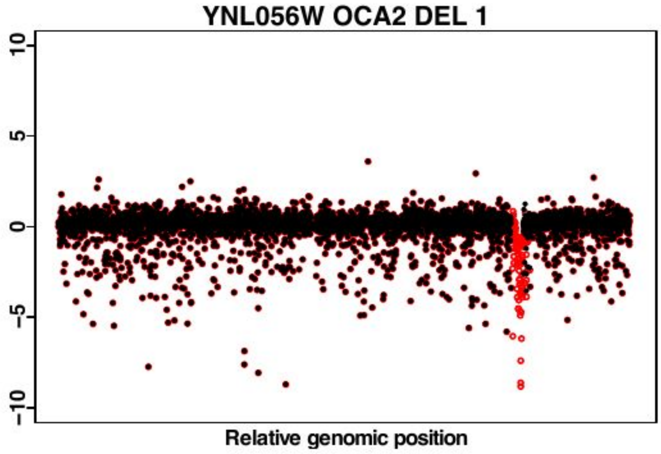


**YNL056W OCA2 DEL 2**

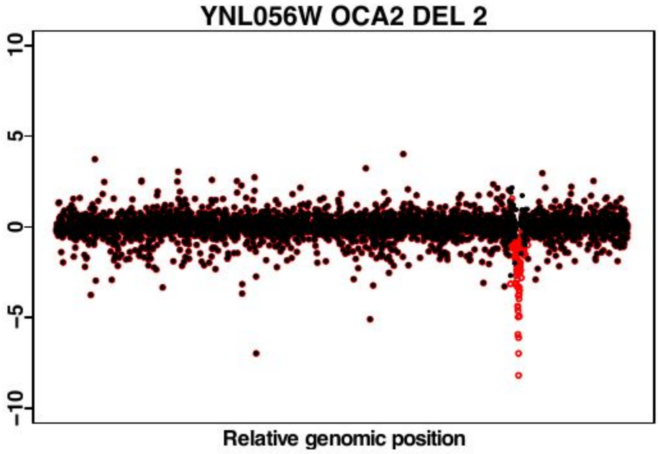


**YNL072W RNH201 DEL 1**

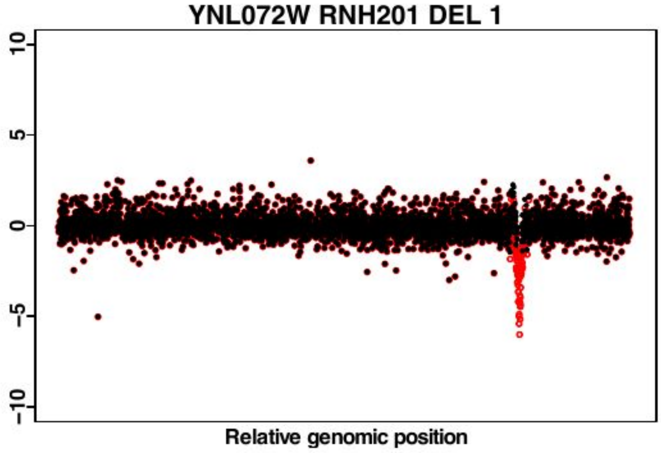


**YNL072W RNH201 DEL 2**

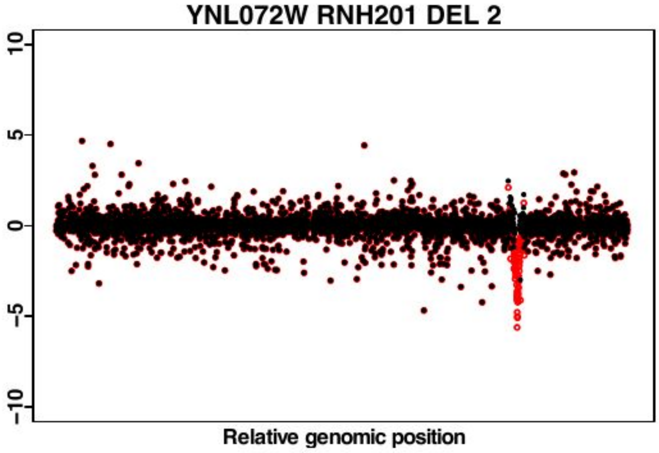


**YNL099C OCA1 DEL 1**

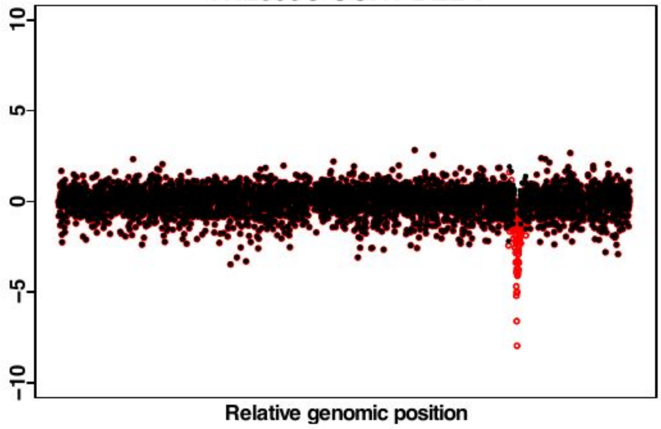


**YNL099C OCA1 DEL 2**

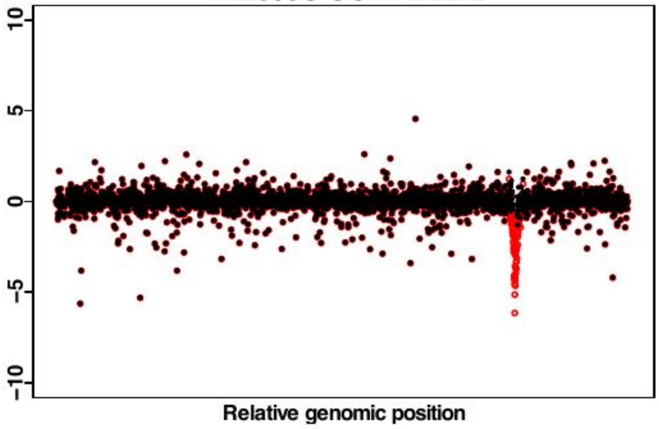

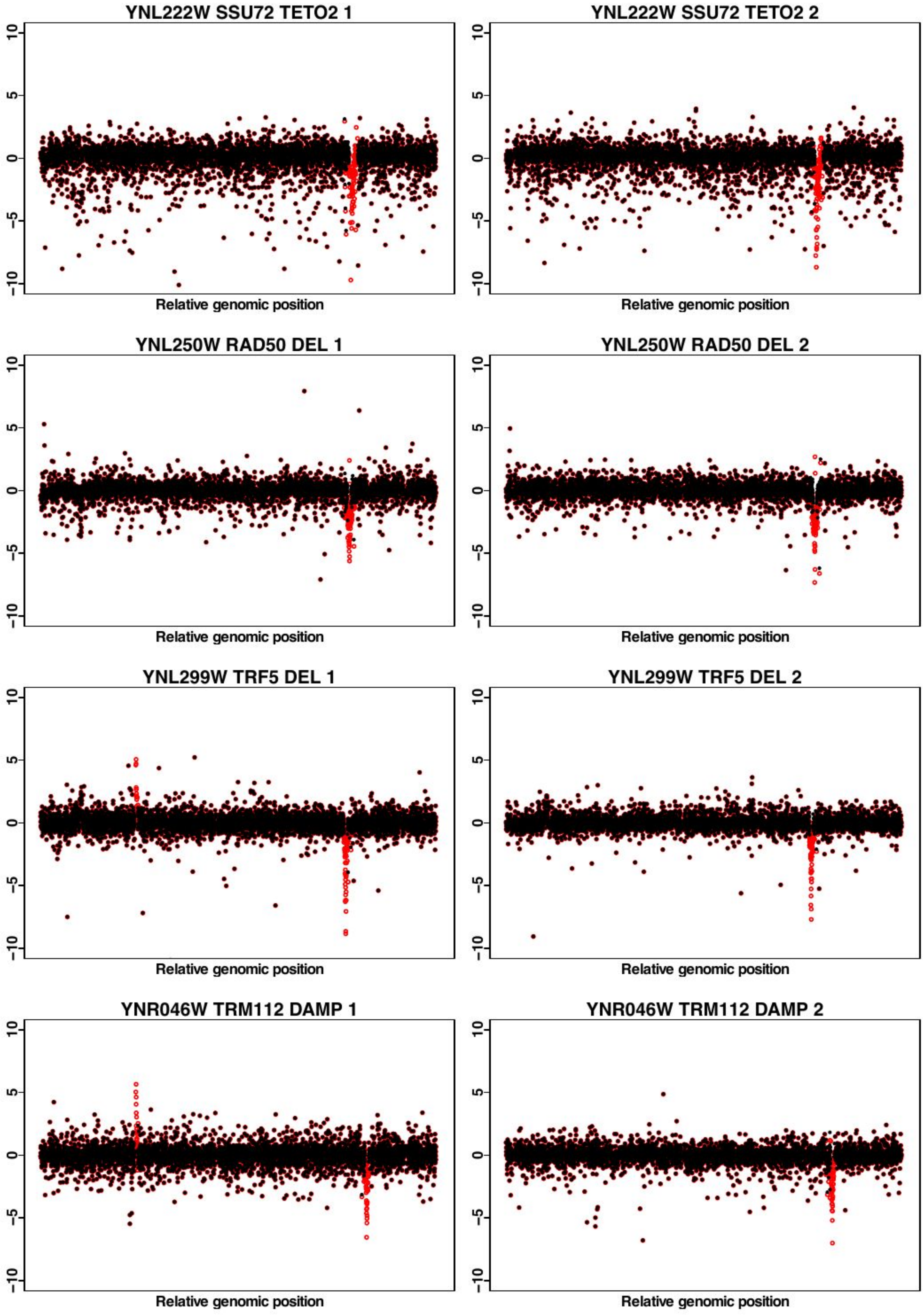
YNL222W SSU72 TETO2 1
YNL222W SSU72 TETO2 2
YNL250W RAD50 DEL 1
YNL250W RAD50 DEL 2
YNL299W TRF5 DEL 1
YNL299W TRF5 DEL 2
YNR046W TRM112 DAMP 1
YNR046W TRM112 DAMP 2
10
5
0
-5
-10
Relative genomic position

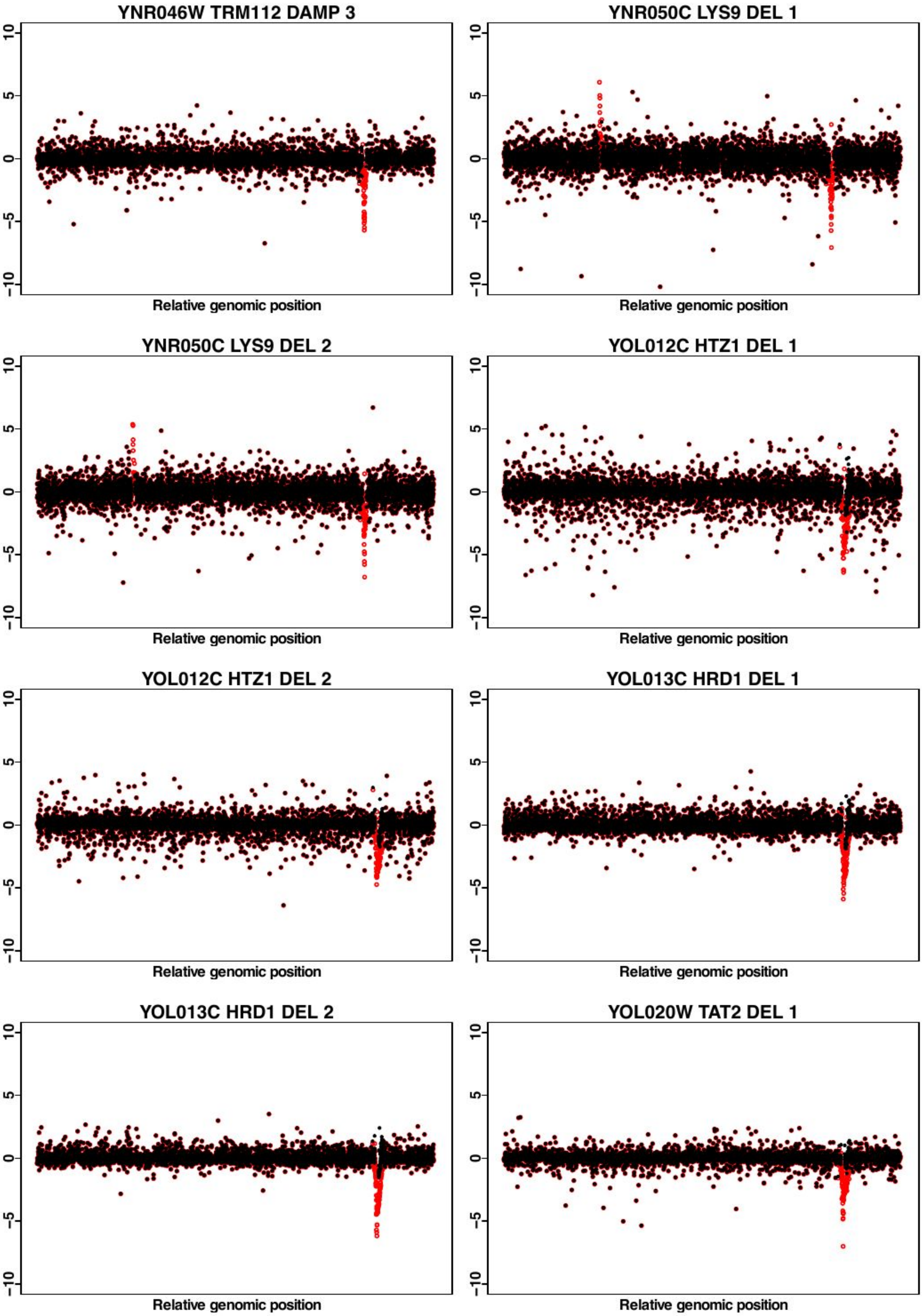
YNR046W TRM112 DAMP 3
YNR050C LYS9 DEL 1
YNR050C LYS9 DEL 2
YOL012C HTZ1 DEL 1
YOL012C HTZ1 DEL 2
YOL013C HRD1 DEL 1
YOL013C HRD1 DEL 2
YOL020W TAT2 DEL 1
10
5
0
-5
-10
Relative genomic position

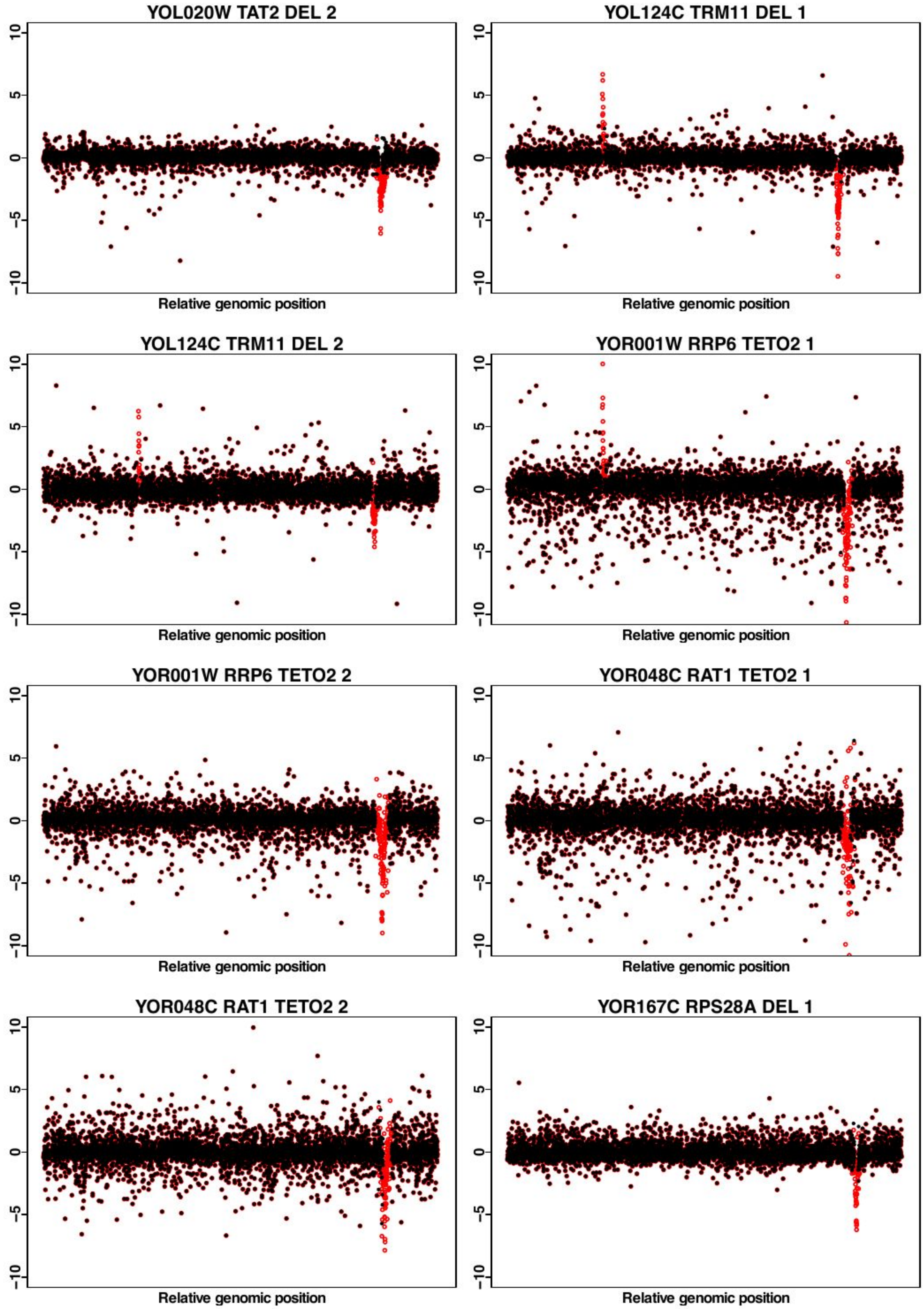

YOL020W TAT2 DEL 2
YOL124C TRM11 DEL 1
YOL124C TRM11 DEL 2
YOR001W RRP6 TETO2 1
YOR001W RRP6 TETO2 2
YOR048C RAT1 TETO2 1
YOR048C RAT1 TETO2 2
YOR167C RPS28A DEL 1
10
5
0
-5
-10
Relative genomic position

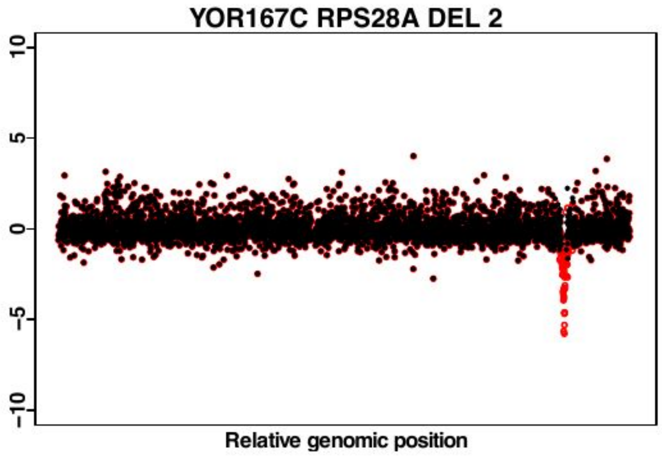
YOR167C RPS28A DEL 2
10
5
0
-5
-10
Relative genomic position

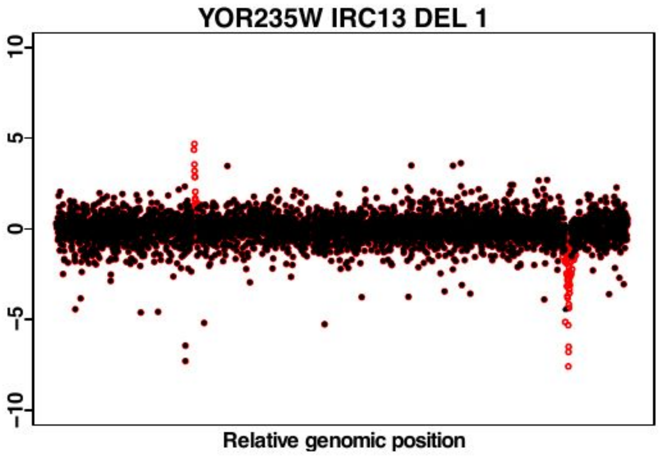
YOR235W IRC13 DEL 1
10
5
0
-5
-10
Relative genomic position

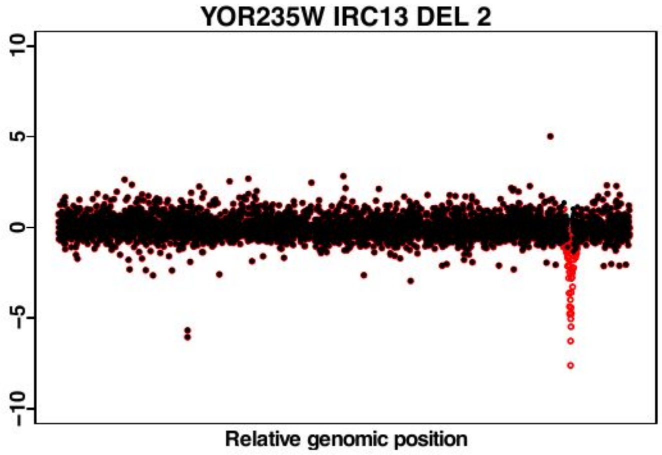
YOR235W IRC13 DEL 2
10
5
0
-5
-10
Relative genomic position

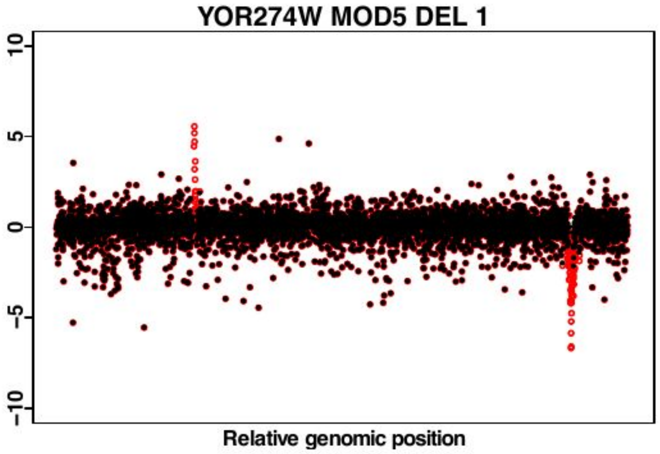
YOR274W MOD5 DEL 1
10
5
0
-5
-10
Relative genomic position

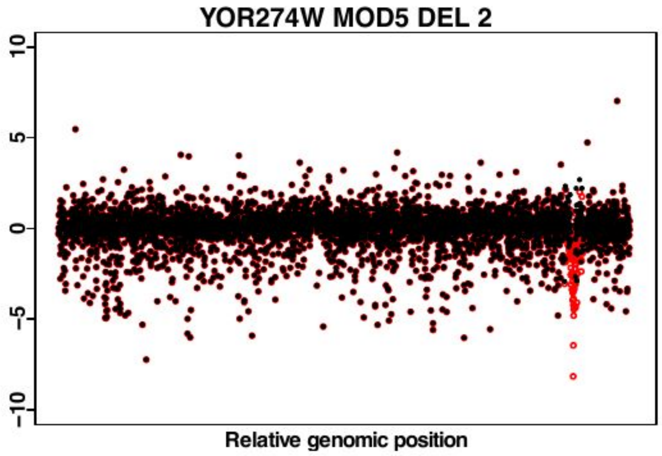
YOR274W MOD5 DEL 2
10
5
0
-5
-10
Relative genomic position

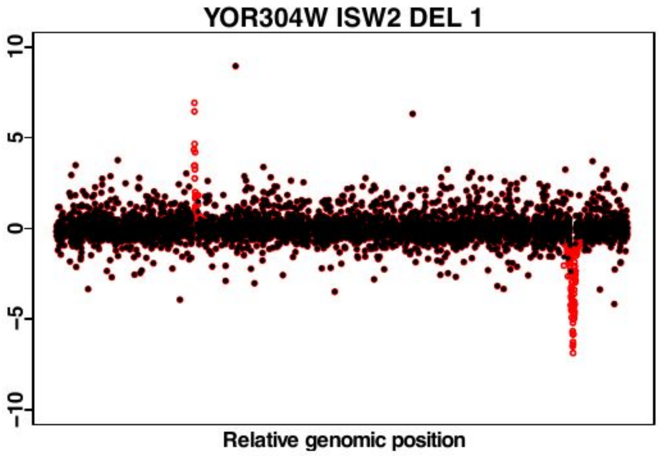
YOR304W ISW2 DEL 1
10
5
0
-5
-10
Relative genomic position

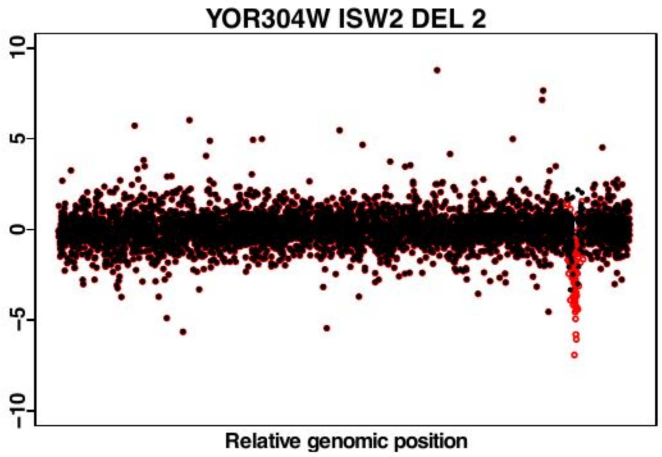
YOR304W ISW2 DEL 2
10
5
0
-5
-10
Relative genomic position

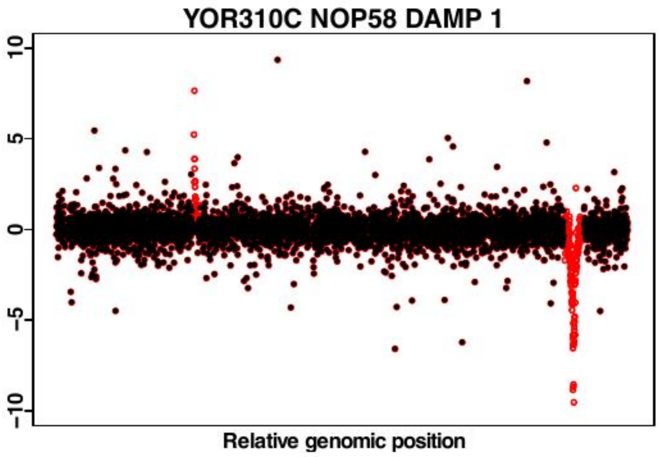
YOR310C NOP58 DAMP 1
10
5
0
-5
-10
Relative genomic position

YOR310C NOP58 DAMP 2

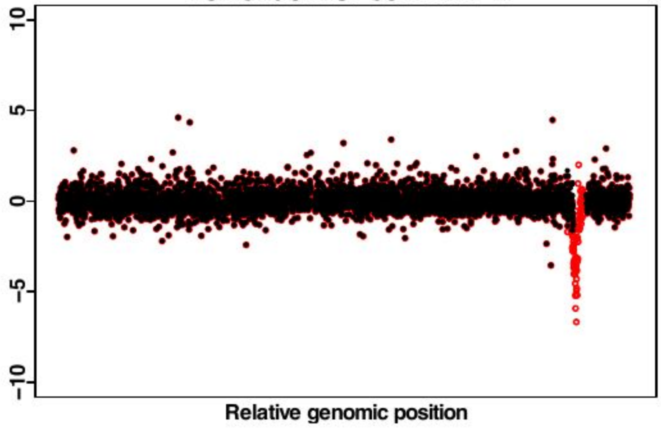


YOR339C UBC11 DEL 1

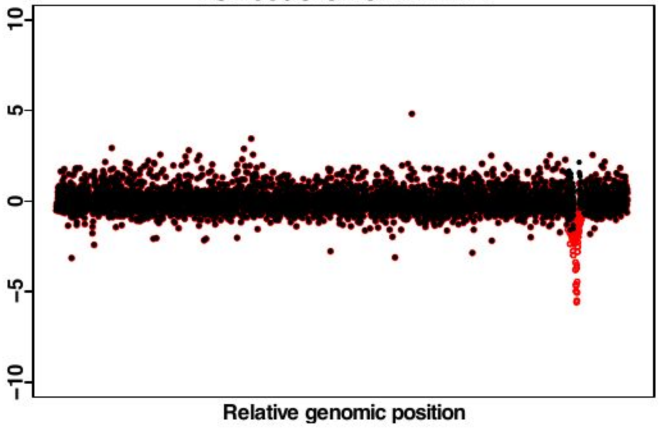


YOR339C UBC11 DEL 2

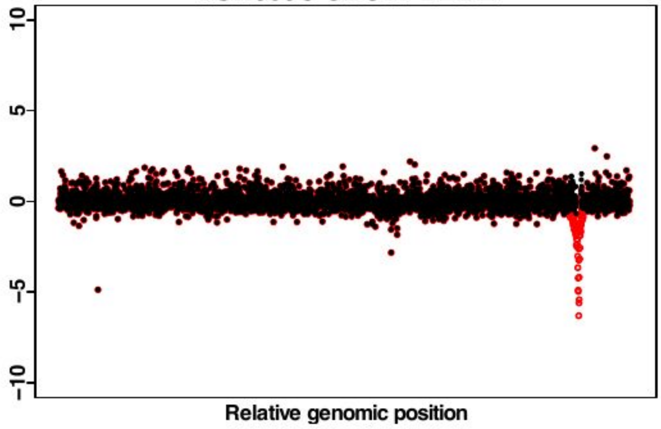


YPL093W NOG1 DAMP 1

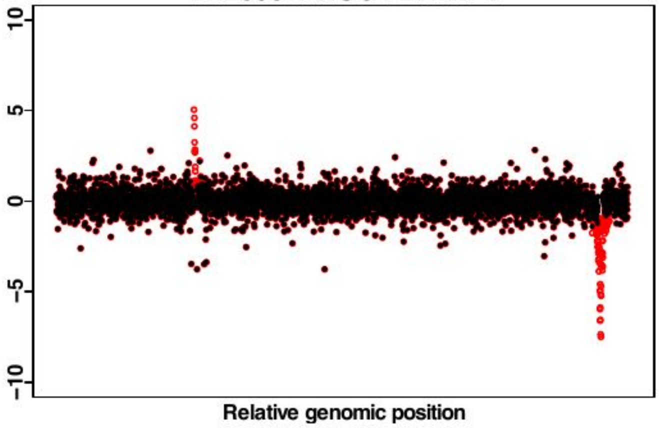


YPL093W NOG1 DAMP 2

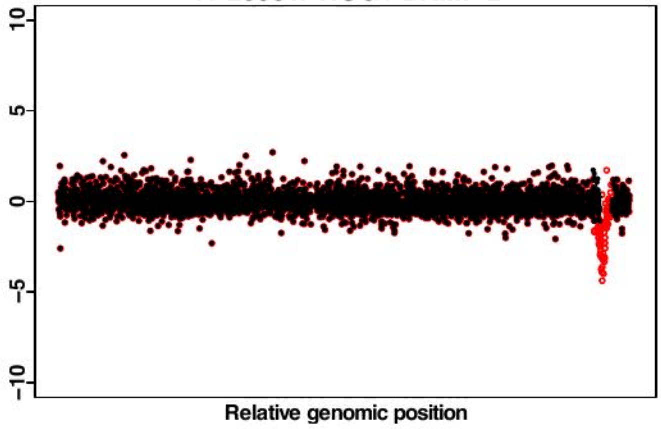


YPL144W POC4 DEL 1

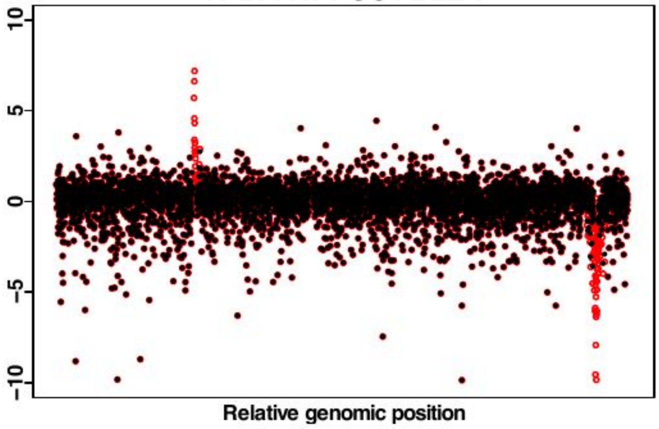


YPL144W POC4 DEL 2

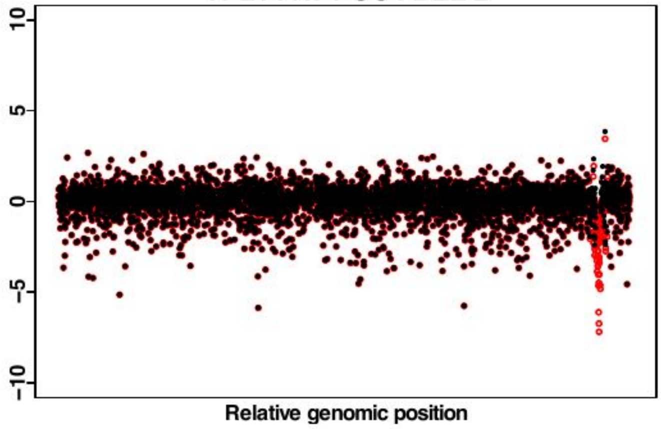


YPL184C MRN1 DEL 1

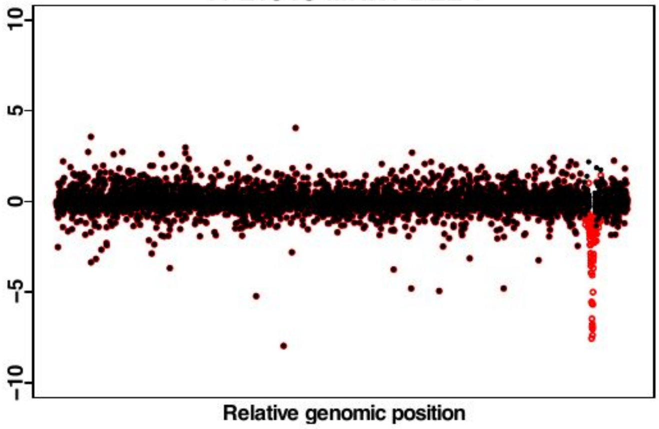

YPL184C MRN1 DEL 2

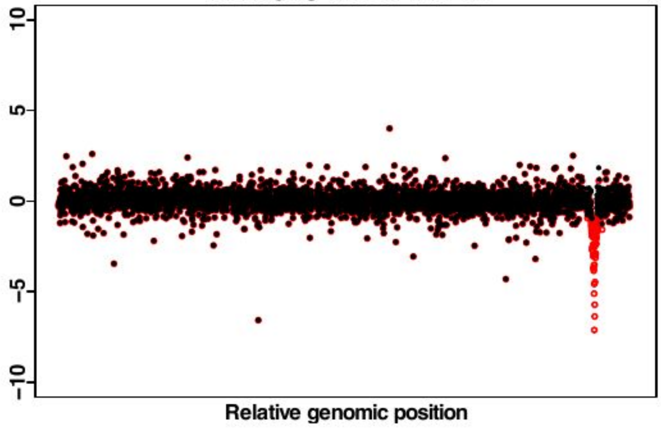


YPL198W RPL7B DEL 1

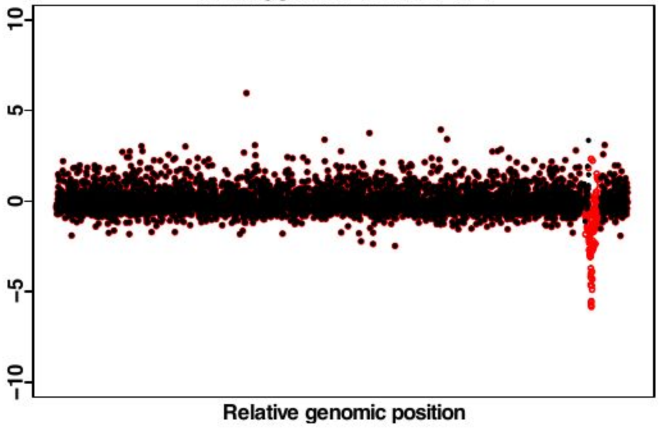


YPL198W RPL7B DEL 2

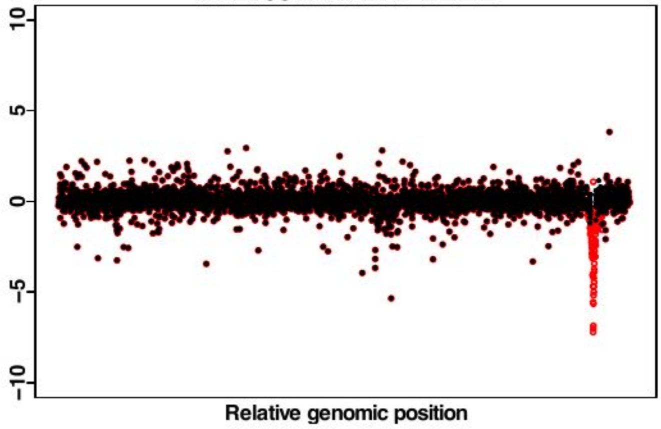


YPL200W CSM4 DEL 1

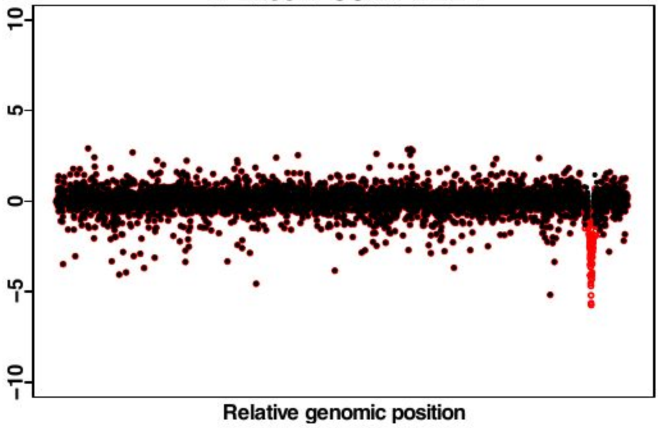


YPL200W CSM4 DEL 2

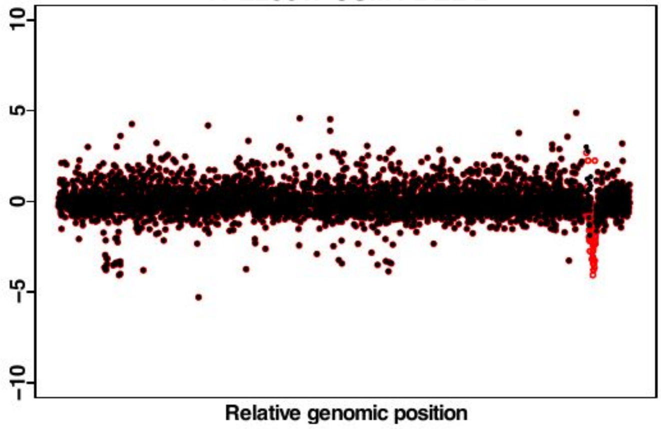


YPL212C PUS1 DEL 1

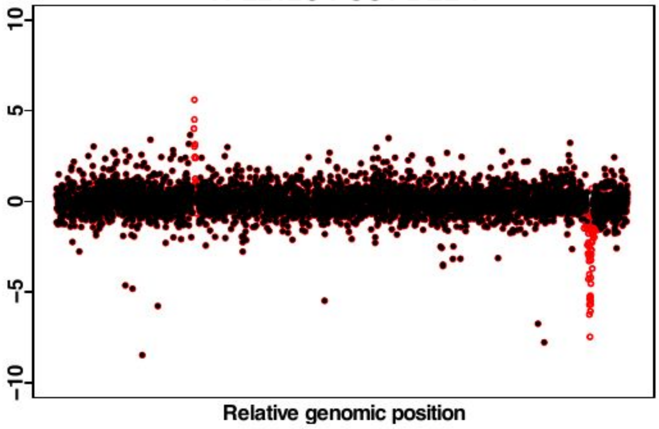


YPL212C PUS1 DEL 2

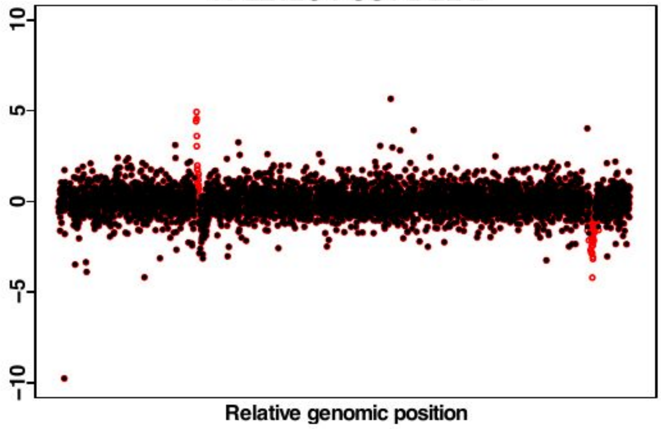


YPL212C PUS1 DEL 3

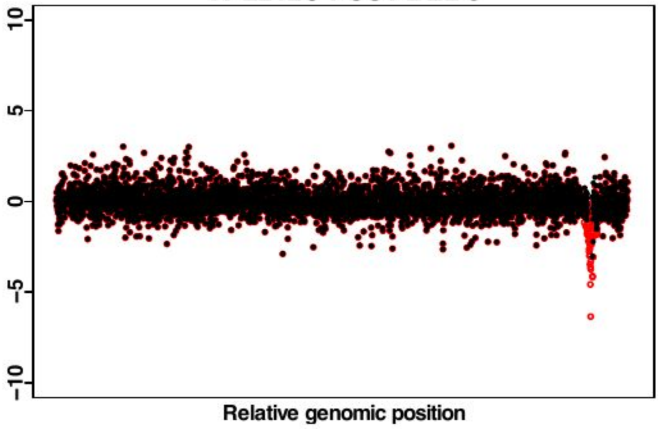

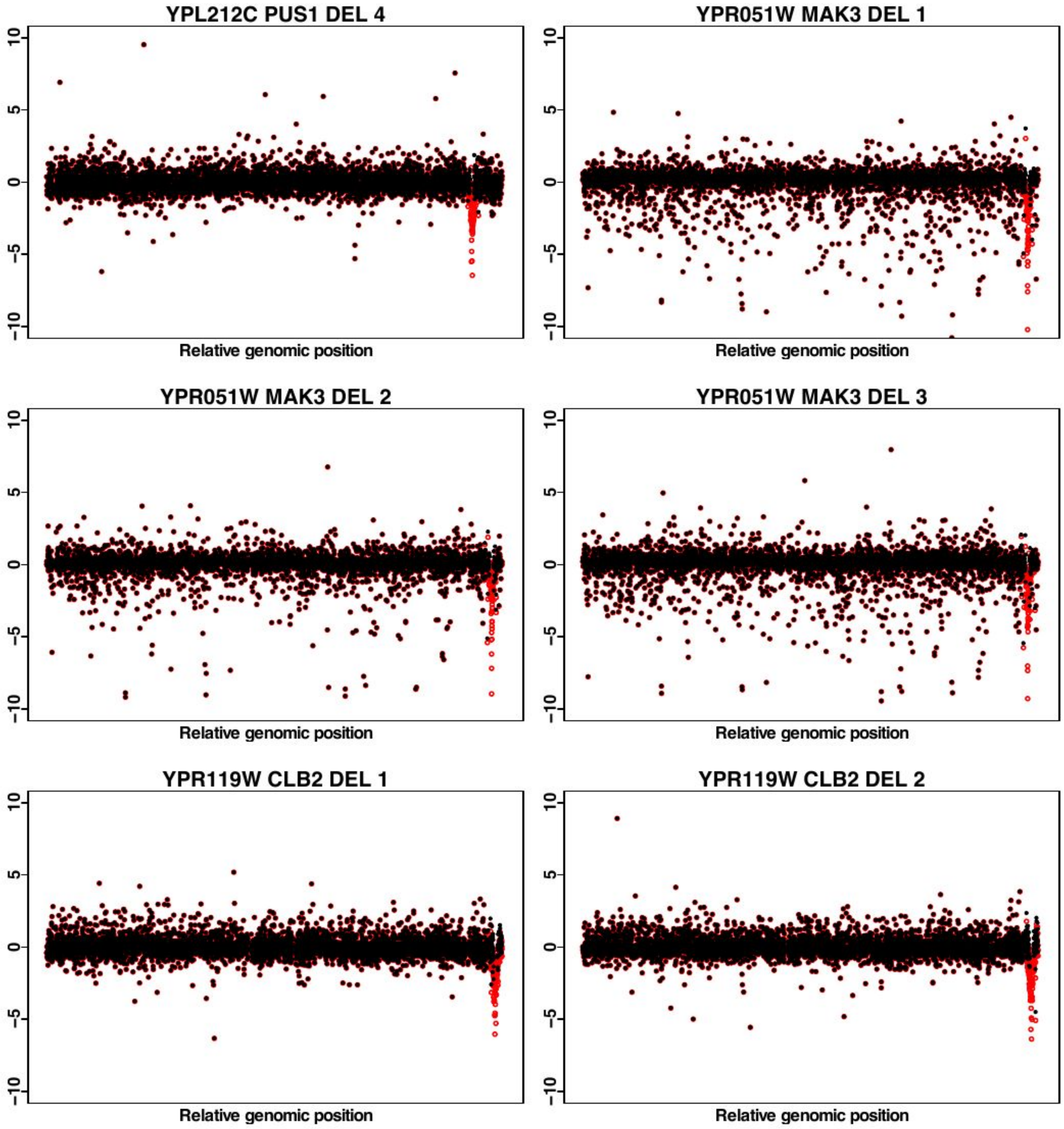
YPL212C PUS1 DEL 4
YPR051W MAK3 DEL 1
YPR051W MAK3 DEL 2
YPR051W MAK3 DEL 3
YPR119W CLB2 DEL 1
YPR119W CLB2 DEL 2
Relative genomic position
10
5
0
-5
-10
